# Ly6d expression delineates two putative postnatal thymus epithelial progenitor cells that are differentially affected by ageing

**DOI:** 10.1101/2025.08.28.672880

**Authors:** Irene Calvo-Asensio, Andreas Tarcevski, Fatima Dhalla, Anja Kusch, Thomas Barthlott, Saulius Zuklys, Georg A. Holländer, Michael D. Morgan

## Abstract

The thymus is a primary lymphoid organ which provides essential structural and functional support for the development of naïve T cells. Thymic epithelial cells (TECs), key components of the thymic stroma, are classified into cortical (cTEC) and medullary (mTEC) lineages based on their distinct molecular, structural, transcriptional, and functional characteristics. Advances in single-cell RNA sequencing (scRNA-seq) have revealed significant TEC heterogeneity, including the identification of intertypical TECs that share properties of both cTEC and mTEC and have been postulated to play a role in the development and maintenance of thymic function. To date, the identity and maintenance of postnatal TEPCs remain unclear, with debates on whether bipotent TEPCs persist after birth or if lineage-restricted progenitors independently maintain TEC compartments. Using an inducible lineage-tracing system based on β5t expression, we explored the early dynamics of the relationships between TEPC and mTEC progenitors and their progeny. Our results identified two potential lineage-biassed TEPC subpopulations, distinguished by Ly6d expression. Additionally, we observed that ageing disproportionately affects Ly6d-compared to Ly6d+ TEPCs, with implications for the rejuvenation of the ageing thymic epithelium. This study provides insights into the developmental pathways of TEC lineages and their maintenance, contributing to strategies for enhancing thymic function in ageing and disease.

## Introduction

The thymus provides as a primary lymphoid organ the necessary structural and functional support for the development of naïve T cells collectively expressing an antigen receptor repertoire purged of reactivity to an individual’s tissue antigens yet poised to react to potentially injurious foreign antigens^1–3^. Thymic epithelial cells (TECs) form together with mesenchymal, endothelial and hematopoietic cell types the major stromal components. TEC are categorized into separate cortical (c) and medullary (m) lineages based on their specific spatial, molecular, structural, transcriptional and thus functional characteristics. cTEC induce the commitment of blood-borne precursor cells to a T cell fate, foster their subsequent expansion, guide their maturation and control both positive and negative selection of antigen receptor bearing thymocytes^4–7^. In contrast, mTEC promote the terminal differentiation of thymocytes which includes the establishment of immunological tolerance to self-antigens via a deletional mechanism (negative thymocyte selection) but also promote the generation of natural regulatory T cells^8,9^. The critical need for T cell antigen receptor (TCR)-mediated negative selection depends on mTEC’s promiscuous expression of transcripts that encode proteins which are normally only detected in organs other than the thymus. Employing flow cytometry, limited number of markers have been identified that distinguish mTEC from cTEC, and that classify separate maturational TEC stages^10^ such as Ulex Europeaus Agglutinin 1 (UEA1) lectin reactivity and Ly51, respectively^11^.

Advances in single-cell RNA sequencing (scRNA-seq) have uncovered a previously unappreciated degree of TEC heterogeneity^12–17^. For clarity, we distinguish flow cytometry phenotypes from scRNA-seq cell types as subpopulations and subtypes, respectively. Using scRNA-seq, we recently described intertypical TEC^12^, so named because they share transcriptomic properties of both cTEC and mTEC. Intertypical TEC are characterized by the expression of genes previously associated with thymic epithelial progenitor cells (TEPCs) localized at the cortico-medullary junction (CMJ), such as Pdpn^15,18^, Ccl21a^18,19^, and Sca1^20,21^, as well as low cell-surface levels of expression of MHC class II^22–24^.

The thymus is the first organ to display several hallmarks of ageing, a process referred to as thymic involution characterized by a loss of cellularity, changes in gross anatomy and diminished T cell output with an altered antigen receptor repertoire^12,25^. Similarly, external and iatrogenic insults such as cytoablative treatments and infections lead to compromised thymic functions^23^ resulting in weakened T cell responses to pathogens, diminished immune surveillance, and an elevated susceptibility to autoimmunity and cancer^26,27^. Consequently, over the past few decades, considerable research has been directed towards uncovering strategies that can either preserve or restore thymic functionality^28^. Research to identify, characterise, and test the mechanisms that support and maintain thymic function by TEPCs suggests the presence of both bipotent progenitors of cTEC and mTEC, and corresponding unipotent progenitors for each lineage^18–22,24,29–37^. During embryogenesis, TEC arise from a common TEPC in the third pharyngeal pouch. Evidence suggests these embryonic progenitors are bipotent, giving rise to both the cTEC and mTEC lineages. Embryonic TEPCs phenotypically resemble cTECs due to the common expression of canonical cTEC-specific molecules including CD205, β5t and IL-7^38–40^. In contrast, there is a lack of consensus on the identity of postnatal TEPC and how they maintain TEC throughout life^19,21,22,29,30,34,41^. Some have suggested that self-renewing bi-potent TEPCs persist after birth^21,34,37^, while others advocate for a series of lineage-restricted progenitors that independently maintain the different TEC compartments during adulthood^14,30–33^. In addition, how the newly-described TEC subtypes, such as intertypical TEC, perinatal cTEC or tuft-like mTEC, fit into these different developmental models requires clarification^38^.

Lineage tracing studies have shown that β5t+ cTEC-like TEPCs located at the CMJ can serve as efficient progenitors for mTECs, although their contribution to the mTEC lineage becomes restricted in the postnatal thymus^18^. This raises the question of whether alternative TEPCs contribute to mTEC maintenance in the adult thymus, and what the developmental relationship is between adult and embryonic progenitor populations. We recently suggested that intertypical TEC may constitute a previous missing link between β5t+ progenitors and mature mTEC, based on their progressive age-related quiescence and limited differentiation into the medulla lineage^12^.

Here, we seek to resolve the relationship between postnatal TEPC and mature TEC lineages. Using an inducible lineage-tracing system that depends on β5t expression, we uncover the early kinetics of TEPC-mTEC progenitor-progeny relationships and resolve the contribution of these cells to cTEC, mTEC and intertypical TEC subsets. Moreover, by investigating the earliest stages of TEPC-TEC differentiation and lineage tracing with single cell transcriptomics, we identify 2 candidate lineage-biased TEPC subpopulations delineated by the expression of Ly6d that are spatially segregated in the thymus. Finally, we note the disproportionate impact of ageing on Ly6d-vs. Ly6d+ TEPC, which has consequences for rejuvenation of the ageing thymic epithelium.

## RESULTS

### mTEC-biassed precursors express β5t

Postnatal mTEC are derived from epithelial precursors that express the proteasome subunit β5t encoded by the *Psmb11* gene^39^. Ageing reduces the differentiation efficiency of β5t+ progenitors, raising the possibility that additional TEC progenitor cells with different developmental capacities could contribute to the maintenance of the postnatal thymus epithelium^12,18,30^. To explore this possibility, we used a previously described lineage tracing model to track the progeny of β5t-expressing TEC. This mouse model, designated 3xtg^β5t,^ relies on *Psmb11* promoter activity to drive expression of the reverse tetracycline transactivator (rtTA)^18,30^. Administration of doxycycline (Dox) therefore leads to Cre expression, the removal of a stop-cassette, and subsequent expression of the fluorescent reporter ZsGreen (Supplementary Figure 1A). Previous observations show that this transgenic model also labelled a small fraction of lineage committed mature mTEC, attributed to promiscuous expression of the *Psmb11* locus^18,42^.

We took advantage of this model to study differential fate choices of mTEC from β5t+ or β5t^-^ progenitors. We first investigated whether decreasing Dox doses would reduce the labelling of mature mTEC whilst maintaining the marking of TEC precursors and their immediate progeny. Treatment of 3xtg^β5t^ mice at 1 week of age with a single dose of either 0.3 mg, 0.02 mg, 0.004 mg, or 0.0008mg of Dox labelled a decreasing number of TEC in a dose-dependent fashion (Figure 1A, B and Supplementary Figure 1B). The lowest dose (0.0008 mg) resulted in a very limited labelling across all TEC populations (<10%) and was therefore not further considered. Analysed 48 hours after the Dox pulse (Figure 1C-H), the majority of cTEC were labelled due to their constitutive expression of *Psmb11*^43^. In contrast, only a small fraction of mTEC were labelled at this time-point (0.3 mg - 7.20 ± 3.57%, 0.02 mg - 3.52 ± 1.74%, 0.004 mg - 1.66 ± 0.97%; Supplementary Figure 1C), the majority of which had a mature CD80^hi^Aire^+^ phenotype (0.3 mg - 76.88% ± 4.2%, 0.02 mg - 86.78% ± 3.85%, 0.004 mg - 92.57% ± 4.40%; Figure 1C-F and Supplementary Figure 2A-C), a result attributed to the promiscuous expression of β5t. In addition, higher doses of Dox preferentially labelled, lineage immature Sca1^+^CD80^-^AIRE^-^MHCII^low^ mTEC without any apparent drug toxicity (Figure 1C-G and Supplementary Figure 2A-C). Therefore, we found that the main differences amongst Dox doses with regards to mTEC labelling was not due to differential labelling of mature mTEC as we originally suspected, but rather to a dose-dependent labelling of immature Sca1^+^CD80^-^ AIRE^-^MHCII^low^ mTEC.

**Figure 1:**
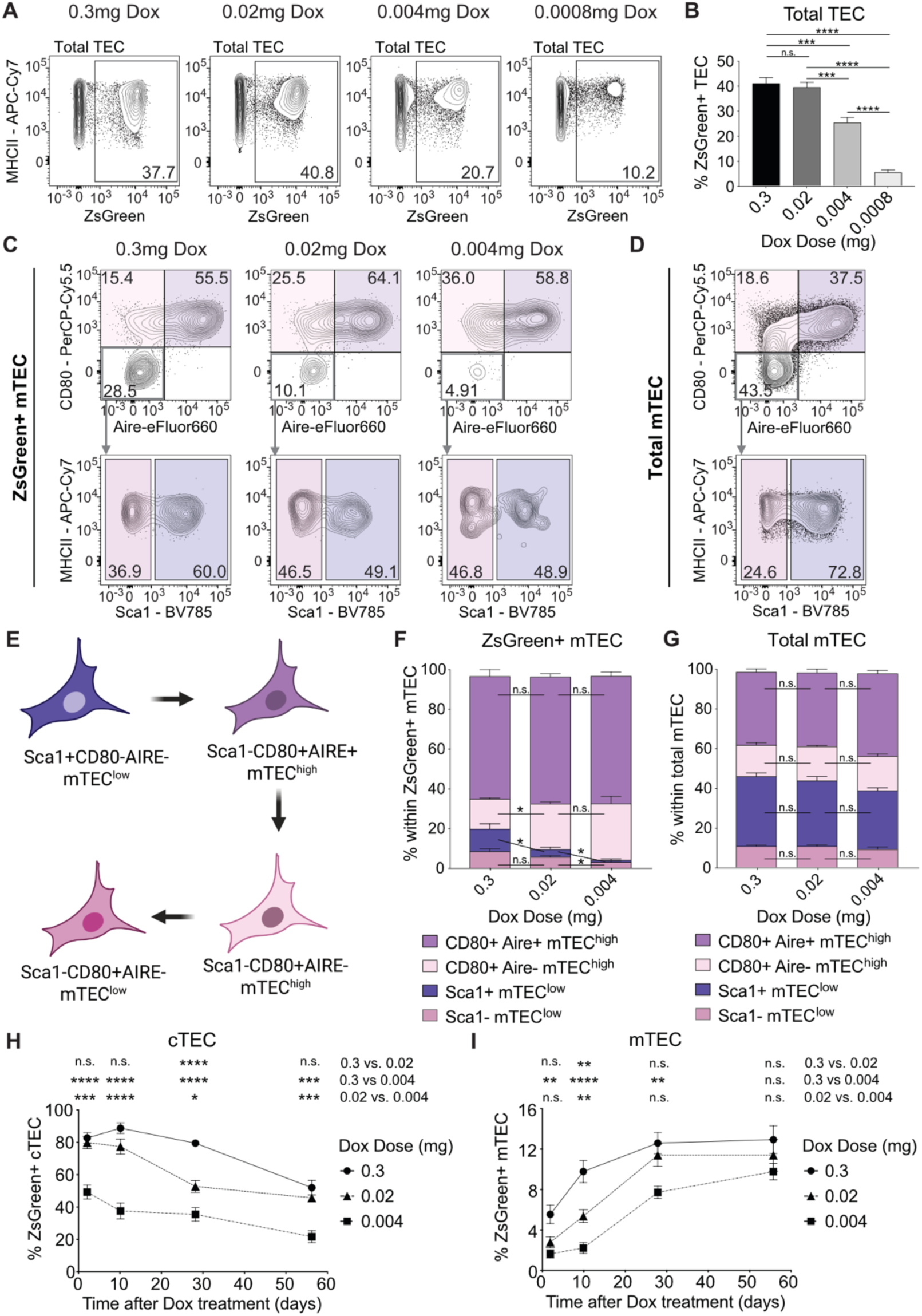
**(A)** Representative FACS plots of ZsGreen expression by total TEC (defined as CD45-EpCAM+ MHCII+) 48 h after 7-day old mice were injected i.p. with the indicated doses of doxycycline (Dox). **(B)** Quantification of the proportion of ZsGreen-labelled TEC, 48h after Dox injection (postnatal day 9). n = 11 mice per group, from 3 independent experiments, except for the 0.0008mg condition which consists of n = 6 mice from two independent experiments. Error bars represent the standard error of the mean. Statistical significance was determined by multiple unpaired Welsch t tests, * p < 0.05, ** p < 0.01, *** p < 0.001, **** p < 0.0001. **(C)** and **(D)** Representative FACS plots of mTEC (C) and total TEC (D) depicting their expression of MHC-II, CD80, AIRE, and Sca1 48h after Dox injection. (**E**) Schematic representation of mTEC subpopulation lineage relationships. **(F)** and **(G)** Quantification of the relative proportions of the different ZsGreen+ mTEC (E) and total mTEC (F); n = 6 mice per group, from 2 independent experiments. Statistical significance was determined by multiple unpaired Welsch t tests, * p < 0.05, ** p < 0.01, *** p < 0.001, **** p < 0.0001. Quantification of the relative proportions of ZsGreen-labelled **(H)** cTEC (Ly51+ UEA1-) and **(I)** mTEC (Ly51-UEA1+), over the course of 8 weeks after injection with decreasing doses of Dox. n = 11 mice per time-point and per treatment group, except for the 56-day time point of the 0.3mg dose, for which n = 10. Statistical significance was determined by mixed-effects model, using the maximum likelihood method. Correction for multiple comparisons was performed using the Turkey method, and p values were adjusted accordingly. The results from the multiple comparisons are depicted as follows: * p < 0.05, ** p < 0.01, *** p < 0.001, **** p < 0.0001. Error bars represent the standard error of the mean.

To investigate the long-term dynamics of TEC progenitor labelling, we studied 3xtg^β5t^ mice 10, 28 and 56 days after Dox treatment. Our analysis revealed differential labelling dynamics across individual TEC subpopulations. For example, the frequency of labelled cTEC decreased over time by ∼20%, which was independent of the Dox dose used (Figure 1H). In contrast, the frequency of labelled mTEC increased over the same time frame, resulting in equivalent proportions of labelled cells across the different Dox doses after 8 weeks (Figure 1I). These data both confirmed the contribution of β5t-positive progenitors to the mTEC lineage and suggested that any of the Dox doses used in our experiments were similarly effective in labelling mTEC long term. Therefore, we used a single Dox dose of 0.004 mg for all subsequent lineage-tracing experiments.

### Developmental dynamics of TEC differentiation

To gain insight into the temporal dynamics of TEC progenitor-progeny relationships, we examined how the frequency of labelled subpopulations changed during the 8 weeks following Dox treatment of 7-day old mice (Figure 2A). In agreement with our previous findings^12^, cTEC and mTEC contributed relatively equally to the total TEC number early in life (postnatal day 9: cTEC: 39.8% ± 4.2%; mTEC: 37.7% ± 2.9%; Supplementary Figure 3A). By 8 weeks post-Dox treatment, mTEC comprised a larger fraction of labelled TEC compared to either phenotype of cTEC (see below week 9; cTEC: 6.5% ± 2.1% vs. mTEC: 35.6% ± 5.7%; Figure 2A-B). In parallel, the frequency of intertypical TEC, which we defined based on the co-expression of CCL21 and Sca1, increased progressively over the same time period (postnatal day 9: 18.2% ± 2.6%; week 5: 43.9% ± 1.7%; week 9: 56.2% ± 3.4%; Supplementary Figure 3A, 4 and 5).

**Figure 2:**
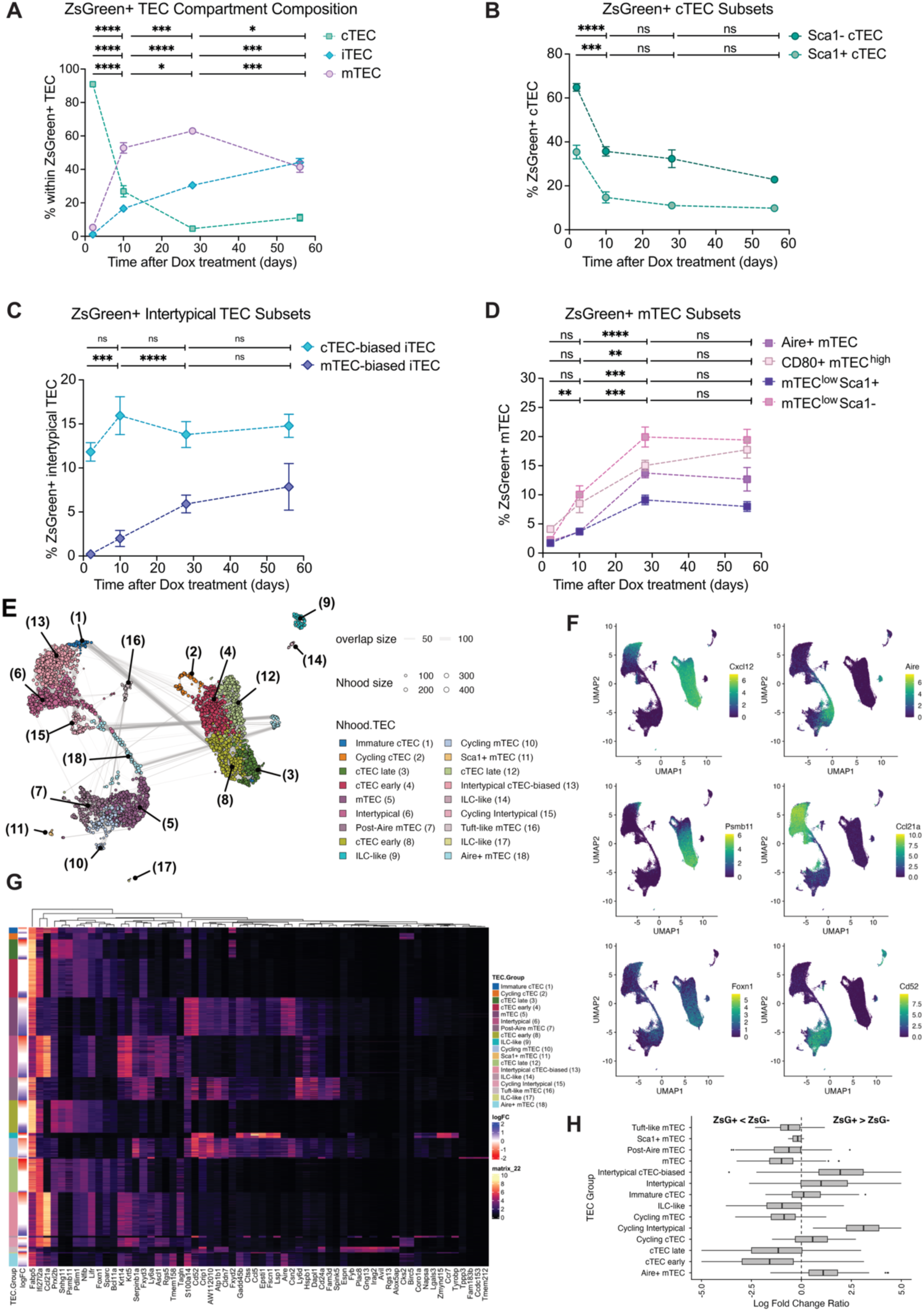
Differential progenitor-progeny relationships revealed by lineage tracing and scRNA-seq. **(A-D)** Analysis of ZsGreen+ TEC compartment composition among (A) total TEC, (B) cTEC, (C) Intertypical TEC and (D) mTEC following i.p. injection of 0.004 mg Dox at 7 days of age and a chase period of 2, 10, 28 and 56 days. n = 7 mice for the analysis at day 2 and n = 8 for that at 10, 28- and 56 after Dox treatment. Error bars represent the standard error of the mean. Statistical significance was determined by using multiple t tests to calculate differences in the TEC subset composition between pairs of consecutive time points. Correction for multiple comparisons was performed using the Holm-Sidak method. * p<0.05, ** p<0.01, *** p<0.001, and **** p<0.0001. **(E)** Neighbourhoods anchored by single-cell UMAP co-ordinates illustrating TEC heterogeneity. Points represent neighbourhoods with UMAP coordinates defined by the index cell for each. Colours denote TEC subpopulation annotations based on marker genes (see **G**). Point size displays the number of cells in a neighbourhood and connecting lines represent the number of overlapping cells between each pair of neighbourhoods. Numbers are annotated for each corresponding cluster. **(F)** UMAP of postnatal EpCAM+ single-cell transcriptomes. Each panel is coloured by markers of TEC lineages: mTEC (*Aire, Cd52*), cTEC (*Foxn1, Psmb11* and *Cxcl12*), intertypical TEC (*Ccl21a*). **(G)** Heatmap of TEC sub population marker genes. **(H)** Log fold change ratio (x-axis) across annotated neighbourhood group (y-axis). Box plots show the median LFC ratio, box edges are the lower and upper quartiles with whiskers extending to 1.5x the IQR. Outliers beyond this range are denoted as individual points.

We further examined the contribution of TEC progenitors to cTEC, mTEC and intertypical TEC, assessed by the frequency of ZsGreen-positive cells in each of these subpopulations over the indicated time course (Figure 2A-D). The initial high frequency of ZsGreen+ cTEC (day 9: 91.1% ± 1.5%) decreased rapidly over time reaching a low level 4 weeks after Dox treatment, which remained largely unchanged thereafter (11% ± 5.6%; Figure 2A, Supplementary Figure 5A). cTEC can be further classified into two separate subpopulations based on their MHCII and Sca1 expression. Based on these two markers, MHCII^hi^Sca1^-^ cTEC (designated cTEC^hi^) would represent a mature cTEC population, while MHCII^low^ Sca1^+^ (cTEC^lo^) have been proposed to constitute a more immature cTEC state^44^. cTEC^hi^ were highly abundant in 9 day old mice but their frequency reduced over time and matched that of cTEC^lo^ in 9 week old animals (Supplementary Figure 5B). Although the percentage of labelled cells differed between the two cTEC phenotypes, the decrease of ZsGreen positivity over time was comparable (Figure 2B), rendering a precursor-progeny relationship between these two cTEC subpopulations unlikely.

The frequency of intertypical TEC as a fraction of all TEC increased 3-fold over the time course, whereas the proportion of ZsGreen positive cells among intertypical TEC increased almost 50-fold (day 9: 0.96% ± 0.54% vs week 9: 44.1% ± 6.7%) (Figure 2A and Supplementary Figure 3A). Based on similar ageing dynamics, we previously hypothesised that intertypical TEC accumulate in a lineage-biased state that restricts their differentiation into mature mTEC^12^. To investigate the lineage bias of intertypical TEC, we used UEA1 and Ly51, respectively, to identify any skewing to either a medullary or a cortical TEC phenotype. We observed that the frequency of ZsGreen+ cTEC-biased Intertypical TEC (Ly51+CCL21+Sca1+UEA1-) remained relatively stable over the 56 days of our time course (Figure 2C). In contrast, and over the same time frame, we observed a sustained accumulation of ZsGreen+ mTEC-biased intertypical TEC (Ly51^-^CCL21^+^Sca1^+^UEA1+; Figure 2C). Our results show that the labelling kinetics of progeny derived from β5t+ progenitors shifts, preferentially give rise to mTEC lineage-biased precursors (Figure 2C).

Based on this observation, we next sought to trace the potential of ZsGreen+ mTEC-biased intertypical TEC. To achieve this, we re-classified mTEC into sequential stages of ontologically separate subpopulations based on MHCII, Sca1, CD80 and AIRE expression (Figure 1E). In this scheme, MHCII^low^Sca1^+^CD80^low^AIRE^-^ mTEC represent an immature stage that differentiates into a mature (MHCII^+^Sca1^-^CD80^+^AIRE^+^) and finally transition via a MHCII^high^CD80^+^AIRE^-^ state to a terminally differentiated subpopulation of MHC^low^Sca1^-^ CD80^+^AIRE^-^ mTEC^20,44–46^. Over time, the frequencies of all but the least mature mTEC decreased (Supplementary Figure 3D). However, the proportion of ZsGreen+ cells increased progressively up to 4 weeks after labelling, whereupon their relative frequency plateaued (Figure 2D). These dynamics suggest that mTEC-biased ZsGreen^+^ intertypical TEC eventually contribute to all stages of mTEC maturity. However, these observations could also be explained by the biased expansion of a particular subpopulation of ZsGreen+ intertypical TEC (which have not yet been further identified using convenient cell surface markers) or mTEC. To exclude this possibility, we determined the proliferation rate of TEC subpopulations 2 days post Dox treatment in relation to their ZsGreen-labelling status. We found that ZsGreen+ cells were less proliferative than unlabelled cells, as measured by Ki67 staining (Supplementary Figure 3E-G), ruling out biased proliferation in ZsGreen+ cells.

In summary, our results indicate that β5t+ progenitors progressively contribute to each of the TEC lineages examined here. Moreover, the inheritance of ZsGreen+ label from β5t^+^ mother to daughter cells suggests that labelled intertypical TEC preferentially give rise to mTEC, and that medulla lineage-biased intertypical TEC accumulate progressively over the first 9 weeks of the life course, a finding in agreement with our previous observations in aged mice^12^. Our results further exclude a continuous contribution of β5t^+^ cTEC precursors into the intertypical TEC pool. Finally, the large reduction in ZsGreen+ cTEC indicates a rapid turnover of these cells early in the postnatal mouse thymus despite the constitutive expression of *Psmb11* in the cortex^22,46^.

### TEC lineage progenitors differentially accumulate in the first weeks of life

The temporal analysis of ZsGreen-labelled Ly51-UEA1+ cells showed as early as 2 days post-Dox that most cells displayed a mature mTEC phenotype (MHC^hi^Sca1-CD80+) in accordance with the promiscuous expression of *Psmb11* (Supplementary Figure 3D). In contrast, 10 days was the earliest time point when a non-zero fraction of immature (MHCII^low^Sca1^+^ CD80^low^AIRE^-^) and post-Aire (MHC^low^Sca1^-^CD80^+^AIRE^-^) phenotypes were labelled (Supplementary Figure 2D). Therefore, to further investigate the early steps in progeny-progenitor relationships, we combined our inducible lineage tracing system with single-cell gene expression profiling at 2- and 10-days post Dox treatment. Based on our and others’ recent findings^14^, we reasoned that a progenitor TEC might express protein markers characteristic of a combined lineage phenotype (bipotent) or alternatively, restricted to either the mTEC or cTEC-specific lineages (unipotent), respectively. We therefore sorted and analysed equal numbers of both ZsGreen- and ZsGreen+ cTEC and mTEC. To prevent batch effects confounding our analysis, we stained the sorted cells with hashtag oligonucleotide-labelled antibodies and multiplexed across individual ZsGreen fractions and time points prior to droplet single-cell RNA-sequencing. Following quality control (Supplementary Figure 6), we integrated single cell gene expression profiles across 4 replicate experiments for each ZsGreen fraction and time point from 64,500 high-quality cells. Visualising marker gene expression on a low dimensional embedding of single cell transcriptomes using uniform manifold approximation and projection (UMAP) indicated the presence of multiple TEC lineages (Figure 2E, F, Supplementary Figure 7). These included mature cTEC (*Cxcl12, Psmb11*), mature mTEC (*Cd52*, *Aire*), and intertypical TEC (*Ccl21a, Lifr*)^12^. Because single-cell gene expression measurements are noisy and sparse, we constructed a set of k-nearest neighbour (kNN) graph neighbourhoods using Milo^47^, which represents the single cell data using overlapping cell states (neighbourhoods).

Clustering similar TEC neighbourhoods yielded 18 clusters, which we annotated based on canonical marker gene expression profiles (Figure 2E-G and Supplementary Figure 7). This analysis revealed the majority of previously described adult mouse TEC lineages^12^, including Tuft-like mTEC (cluster 16) and post-Aire mTEC (cluster 7). This analysis also identified innate lymphoid cells that displayed EpCAM on their surface but in which the corresponding mRNA was not detected (Supplementary Figure 8). These cells were excluded from further analysis. The transcriptional similarity between cells positioned far apart in the UMAP space was more clearly illustrated by visualising the neighbourhood graph (Figure 2E) which resolved the connection between Immature cTEC (cluster 1) and early cTEC (cluster 8), and between intertypical TEC (cluster 6) and Aire+ mTEC (cluster 18). Notably, our clustering analysis indicated multiple neighbourhood clusters that corresponded to intertypical TEC, which we previously hypothesised may contain TEC progenitors based on shared gene expression profiles e.g., *Ccl21a, Ly6a, Lifr* (Figure 2G)^12^.

To identify transcriptional TEC cell states that had different abundance between day 2 and day 10 post-dox treatment, we used differential abundance testing algorithm Milo ^47,48^, analysing ZsGreen+ and ZsGreen-fractions separately (10% false discovery rate; FDR). Considering just the ZsGreen+ cells, our analysis revealed across the 8 day time course an enrichment of intertypical TEC with a cTEC-biased transcription profile (cluster 13; *Cxcl12, Ccl21a, Krt5, Krt14*; Supplementary Figure 9A), a finding concordant with our flow cytometry data (Figure 2C) and the low but detectable expression of *Psmb11* in these cells (Figure 2F).

We next investigated the lineage biases in each neighbourhood cluster of ZsGreen+ and ZsGreen-TEC. To do this, we computed the ratio of the log-fold changes from the separate ZsGreen+ and ZsGreen-Milo analyses for each neighbourhood and grouped them by TEC neighbourhood cluster. This analysis showed an accumulation of ZsGreen-positivity in cycling cTEC-biased intertypical TEC, as well as Aire+ mTEC (Figure 2H), findings consistent with our flow cytometry data (Figure 2A-D). Also consistent with that analysis was the observation of a depletion of ZsGreen-positive cTEC populations, reflecting the relatively high postnatal turn-over of these TEC subtypes (Figure 2B & H)^49^. In contrast, the neighbourhoods representing the remaining mTEC subtypes, including Post-Aire, Tuft-like and cycling mTEC were skewed towards ZsGreen-negativity (Figure 2H).

We identified 3 clusters of dividing cells: cTEC cluster 2, mTEC cluster 10 and intertypical TEC cluster 15. A transit-amplifying population of mTEC has previously been described and given the name TAC-TEC^50^. To assess if any of the proliferating TEC clusters identified here correspond to TAC-TEC, we integrated the two single cell transcriptomic data sets (Supplementary Figure 10). The cross-data set mapping indicated that TAC-TEC best corresponds to a mixed subset of intertypical TEC cluster 6, cycling intertypical TEC (cluster 15) and Aire+ mTEC (cluster 18). This heterogeneity has been noted previously^50^, and our better separation of dividing TEC subtypes is likely explained by the larger number of cells in our data. Notably, cycling intertypical TEC (cluster 15) were strongly enriched in the ZsGreen+ fraction (Figure 2H), which suggests that either they arise from or contain a β5t+ progenitor state, or that the labelling efficiency is specifically skewed in dividing cells (Supplementary Figure 3F); Ki67 staining ruled out the latter explanation (Supplementary Figure 3F). Moreover, as cycling intertypical cells appeared to form a transitional population between intertypical TEC and Aire+ mTEC, and because intertypical TEC also accumulated the ZsGreen-label, our results further support the notion that intertypical TEC cluster 6 may contain the mTEC progenitor.

In summary, our neighbourhood analysis of perinatal TEC development and progenitor-progeny relationships are consistent with at least one β5t+ progenitor. Our results do not delineate between a single bipotent β5t+ progenitor, or two discrete unipotent β5t+ TEPC that give rise to the cTEC and mTEC lineages separately. However, our combined flow cytometry and single-cell data strongly support multiple progenitor populations that share characteristics with intertypical TEC, each likely displaying distinct TEC lineage biases (Figure 2E-G).

### Ly6d expression distinguishes lineage-biased TEC progenitors

To further investigate the possibility that the population of intertypical TEC includes one or more potential progenitor populations, we restricted our single-cell analysis to data corresponding to the least mature TEC (i.e. clusters 1, 6, 13, 15). This selection was supported by cross-mapping to single-cell RNA-sequencing data of human TEC (Supplementary Figure 11)^51^. Human cells corresponding to a poly-keratin signature (described by Ragazzini *et al*.^51^ as human neighbourhood cluster 7) had the highest transcriptional correlation with intertypical TEC and immature mTEC and cTEC in our data (Supplementary Figure 11). Therefore, we constructed a kNN-graph with a newly computed set of highly variable genes and reduced dimensions that captured the gene expression variation across the cells from clusters 1, 6, 13, 15 (19956 cells, Supplementary Figure 12). We then identified marker genes for a new set of 10 neighbourhood clusters and manually annotated these cells (Figure 3A, B). Our higher resolution analysis revealed a heterogeneous mix of TEC subtypes that expressed genes characteristic of intertypical TEC, mTEC and cTEC lineages (Figure 3A, B, Supplementary Figure 13). The immature mTEC (cluster 7) and cTEC states (cluster 5) were broadly separable in our neighbourhood graph, delineated by lineage markers (cTEC: *Ccl25, Prss16, Psmb11*; mTEC: *Fezf2, Aire*). In addition, our neighbourhood graph suggested that an mTEC fate choice was made prior to cell division, indicating that previously described TAC-TEC cells may already be committed to the mTEC lineage (Figure 3A, Supplementary Figure 10).

**Figure 3.**
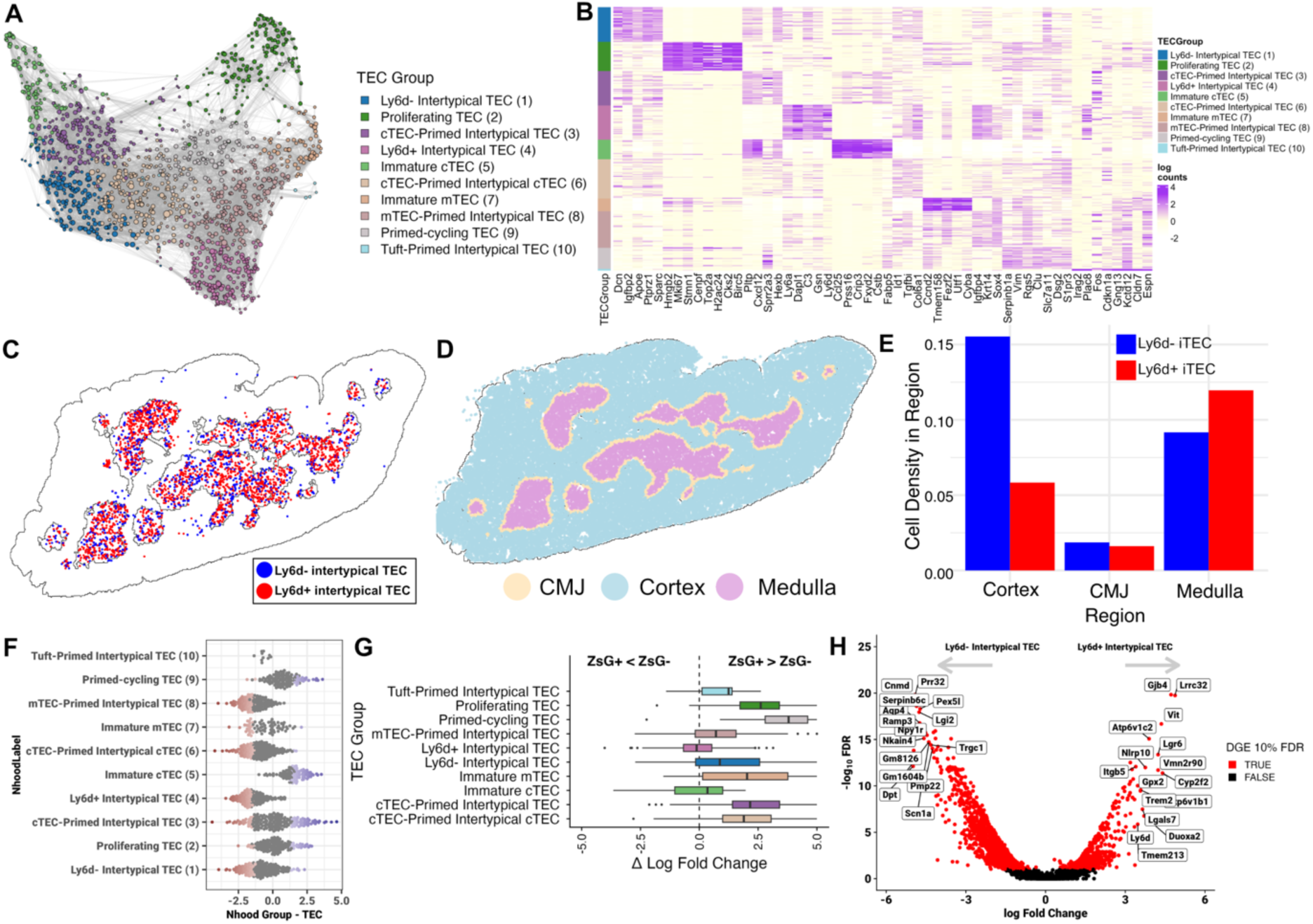
Stratification of progenitor TEC and differential lineage labelling over time. **(A)** UMAP of intertypical TEC neighbourhoods (19956 cells, 1466 neighbourhoods). Points are the 1466 neighbourhood index cells embedded in the single-cell UMAP of intertypical TEC (Supplementary Figure 12), coloured by annotated neighbourhood cluster. Edges represent the number of cells shared between neighbourhoods, and point sizes denote the number of cells in a neighbourhood. **(B)** Heatmap of top marker genes expression for intertypical TEC clusters shown in (A). Rows denote neighbourhoods and columns are marker genes. **(C)** Spatial positions of Ly6d+ and Ly6d-intertypical TEC within the thymus. Each point represents a segmented single intertypical TEC (Ccl21a+Lifr+). **(D)** All spatial transcriptome cells labelled by inferred thymic compartment – corticomedullary junction (CMJ; yellow), cortex (turquoise) or medulla (pink). **(E)** Bar chart showing the density (y-axis) of Ly6d+ (red) and Ly6d-(blue) intertypical TEC within spatial thymic compartments (x-axis) as in **(D)**. Each region is scaled by the total cell density. **(F)** Beeswarm plot of DA neighbourhoods between ZsGreen+ and ZsGreen-lineage traced cells. Points are coloured by log fold change (x-axis). **(G)** Boxplot of log fold change difference between ZsGreen+ and ZsGreen-DA neighbourhoods (10% FDR) for each annotated intertypical TEC state, as in (A). Values greater than zero indicate an accumulation of cells in the ZsGreen+ fraction from day 2 to 10, while values less than zero accumulate cells in the ZsGreen-fraction. Boxplot horizontal bars denote the median ΔLFC, box margins are the interquartile range (IQR), with whiskers extending to 1.5x the IQR; outliers are shown as individual points. Plot colours are as in (A). **(H)** Volcano plot of differentially regulated genes between Ly6d+ (cluster 4) and Ly6d- (cluster 1) intertypical TEC. Statistically significant DEGs (FDR 10%) are coloured in red. Labelled genes denote the top 15 up- and down-regulated genes.

We identified 2 intertypical TEC neighbourhood sub-clusters (1 & 4) which expressed the highest levels of genes previously described as markers for TEC progenitors (*Gas1, Ly6a, Itga6, Plet1*; Supplementary Figure 13)^21,29^ but differed in their expression of *Ly6d* (Figure 3B, Supplementary Figure 13). The Ly6d+ intertypical TEC displayed a pattern of lowly expressed genes shared with mTEC sub-clusters 7 and 8 (e.g., *Cd52, Aire*, *Fezf2;* Supp Figure 13), and expressed transcription factor genes associated with type I interferon signalling (*Stat1, Stat2, Irf6*; Supplementary Figure 14). Recent data shows that interferon signalling is critical for mTEC maturation during human foetal thymus development^52^, indicating Ly6d+ intertypical TEC might be precursors of, or biased towards, the mTEC linage. In contrast, the Ly6d-intertypical TEC shared gene expression patterns with cTEC sub-clusters 3 and 5, including *Dll4* and *Pdpn*, and are characterised by transcription factors associated with differentiation and stem cells (e.g. *Id3, Nfib, Peg3*; Supplementary Figure 14)^53^. We further identified 5 sub-clusters that we call Primed intertypical TEC, so called because they share transcriptional profiles with intertypical TEC and either mTEC (cluster 8), cTEC (clusters 3 & 6), Tuft-like mTEC (cluster 10) or proliferating TEC (cluster 9; Figure 3A-B, Supplementary Figure 13). We speculate that these cells may be in the initial stages of differentiation or making a fate choice towards a particular lineage or the cell cycle. Finally, we note that *Psmb11* is highly expressed in the immature cTEC sub-cluster 5, and more lowly expressed in the cTEC-Primed intertypical TEC and Ly6d^-^ intertypical TEC (Supplementary Figure 13). Together, these analyses suggest the presence of up to two possible progenitor TEC states, where their lineage bias is delineated by the expression of *Ly6d*.

To explore how Ly6d expression marks cortical vs. medullary biased progenitors, we mapped the marker gene profiles for each on to single-cell spatial transcriptomics profiles of an 8-week-old thymus (Fig3C-D; Methods). Using marker genes identified by differential gene expression analysis we computed a combined module score to classify each intertypical TEC (Ccl21a+Lifr+) in our spatial transcriptomics data set as either Ly6d+ or Ly6d-intertypical TEC (Supplementary Figure 15A-D). Re-mapping the module score back onto our original scRNA-seq data using the same genes validates that this approach is selective for Ly6d+ and Lyd6-intertypical TEC (Supplementary Figure 15E-F). Our spatial analysis shows that Ly6d+ intertypical TEC are enriched within the medullary regions, while the Ly6d-intertypical TEC are more likely to be in the cortico-medullary junction (CMJ) or adjacent in the cortex (Fig3E, Supplementary Figure 15G). Together, these complementary data demonstrate that two intertypical TEC populations, delineated by Ly6d expression, are both spatially segregated and characterised by transcriptome profiles indicative of different TEC lineages.

Given the different spatial compartmentalisation of Ly6d+/- intertypical TEC, we wanted to understand their lineage relationship to β5t+ progenitors. To do this, we probed the composition of the ZsGreen+ and ZsGreen-fractions within the intertypical and immature TEC states in our scRNA-seq data. Using Milo^47,48^, our analysis revealed that Immature cTEC, cTEC-primed intertypical TEC (sub-cluster 3) and both primed-cycling (sub-cluster 9) and proliferating TEC (sub-cluster 2) were enriched within the ZsGreen+ fraction (10% FDR, Figure 3C). In contrast, Ly6d-intertypical TEC (sub-cluster 1), Ly6d+ intertypical TEC (sub-cluster 4), and mTEC-primed intertypical TEC (sub-cluster 8) were depleted in that fraction. This analysis indicated that both Ly6d- and Ly6d+ intertypical TEC precede a β5t+ state, irrespective of their apparent lineage biases. To explore this further, we used Milo to compare the abundance of TEC neighbourhoods between 2 and 10 day time points following Dox treatment (FDR 10%; Supplementary Figure 16). Over the course of 8 days, both Ly6d+ and Ly6d-intertypical TEC were enriched over time regardless of whether these cells were ZsGreen positive or negative (Supplementary Figure 16). Additionally, we observed for the ZsGreen+ fraction a relative enrichment of primed-cycling TEC (sub-cluster 9) and cTEC (sub-cluster 5; Supplementary Figure 16) and a corresponding lower frequency of cTEC was noted at 10 days within the ZsGreen-fraction. We reasoned that if intertypical TEC either contained or were the progeny of β5t+ progenitors then a relative accumulation of ZsGreen+ cells should be observed over time. Conversely, should either one or both intertypical TEC populations precede the developmental stage of β5t+ progenitors, a change in the ZsGreen fractions would not occur. Therefore, we computed for each neighbourhood the difference between the Day 2 vs. Day10 log fold changes for the ZsGreen+ and ZsGreen-fraction of TEC (Δ Log Fold Change; ΔLFC, Figure 3D). Our analysis revealed that cTEC-primed intertypical TEC (sub-clusters 3 & 6) accumulated over time in the ZsGreen-labelled fraction, just like proliferating (sub-cluster 2) and primed-cycling TEC (sub-cluster 9). The latter are TEC with a transcriptional profile concordant with both dividing cells (*Mki67, Top2a, Ccnd2*) and an mTEC-like expression profile (*Cd52, Cd40, Aire*; Supplementary Figure 13). While there was a higher percentage of Ly6d^-^ intertypical TEC accumulating in the ZsGreen+ fraction, this was not observed for Ly6d^+^ intertypical TEC (Figure 3D), suggesting Ly6d^+^ intertypical TEC may arise from an unlabelled progenitor, or, alternatively, precede a state of β5t-positivity.

Despite the fact that ZsGreen does not preferentially accumulate in Ly6d^+^ intertypical TEC over our time course, we observed a small fraction that were ZsGreen-positive despite a lack of detectable *Psmb11* expression (Supplementary Figure 12). We hypothesised that this may be due to a stochastic activation of the *Psmb11* promoter. To explore this premise further, we correlated the proportion of ZsGreen+ cells in each neighbourhood with the proportion that expresses TEC progenitor and lineage genes (cortex: *Psmb11, Prss16, Ccl25,* medulla: *Aire, Fezf2,* Progenitor: *Ly6a, Ccl21a, Ly6d*; Supplementary Figure 17A). This analysis revealed that the proportion of ZsGreen^+^ Ly6d^+^ intertypical TEC was positively, albeit weakly, correlated with the proportion of Foxn1^+^ (ρ=0.20, 95% CI [0.06-0.33]) and Psmb11+ cells (ρ=0.28, 95% CI [0.15-0.41]). Correlations between ZsGreen expression and other Foxn1-regulated genes^54^ were notably weaker in comparison (*Ccl25:* ρ=0.12, 95% CI [-0.02-0.26]; *Prss16:* ρ=0.12, 95% CI [-0.02-0.26]). These correlations appeared to be driven by a small subset of neighbourhoods containing ≥ 50% ZsGreen^+^ and ≥25% Foxn1^+^ cells, because the number of Psmb11^+^ cells was low (mean 0.72%; Supplementary Figure 17B). Therefore, weak activation of the *Psmb11* promoter during Dox treatment in Foxn1+ cells, despite the absence of detected *Psmb11* mRNA transcripts, may lead to the stochastic labelling of some TEC that are not the progeny of a β5t+ progenitor.

### Differential signalling cues between Ly6d+ and Ly6d-intertypical TEC

To understand the extracellular signals and gene regulatory programs that define the Ly6d^+^ and Ly6d^-^ intertypical TEC, respectively, we first performed differential gene expression and functional enrichment analyses. We compared gene expression levels between Ly6d^+^ and Ly6d^-^ intertypical TEC across the mouse protein-coding transcriptome and identified 1992 differentially expressed (DE) genes (1% FDR, Figure 3E). These included, in addition to *Ly6d*, the R-spondin receptor *Lgr6*, Leucine rich repeat containing 32 (*Lrcc32*), also known as GARP, and Galectin 7 encoded by *Lgals7*, which had previously been observed in Hassal’s corpuscles^55^. This expression profile indicated both paracrine and cell-matrix interactions by Ly6d^+^ intertypical TEC. To gain a broader insight into the gene regulatory programs in each cell type, we performed gene set enrichment analyses (GSEA, Supplementary Figure 18). Ly6d-intertypical TEC had up-regulated genes related to mitosis and cell cycle regulation (e.g. Myc Targets, E2F Targets, G2M Checkpoint), indicating the cells’ license for cell division, and identifying them to be likely immediate progenitors of the primed-cycling TEC (sub-cluster 9). Retinol metabolism genes were enriched in the Ly6d^+^ intertypical TEC (Supplementary Figure 18), which is required for *in vivo* and *in vitro* thymus organogenesis^45,56^, suggesting an ongoing role for retinoic acid in postnatal TEC progenitors.

Based on the observed differences in expression of cell-cell and cell-ECM interaction genes between Ly6d+ and Ly6d-intertypical TEC, we next sought to identify the paracrine signalling pathways active in the postnatal thymic epithelium that could be important for TEPC. To achieve this we first merged the annotation of all TEC clusters (Figure 2E) with the higher resolution intertypical TEC sub-clusters (Figure 3A, Supplementary Figure 19A). Using *CellChat* we then identified 91 signalling pathways (Supplementary Figure 19B), including Wnt, BMP, and Notch which have known roles in thymus function, organogenesis and TEC lineage differentiation^44,57–61^. Using non-negative matrix factorisation of the cell-cell communication network we identified a complex landscape of incoming and outgoing signalling modules that were largely distinct for intertypical TEC, cTEC and mTEC (Figure 4A-B).

**Figure 4.**
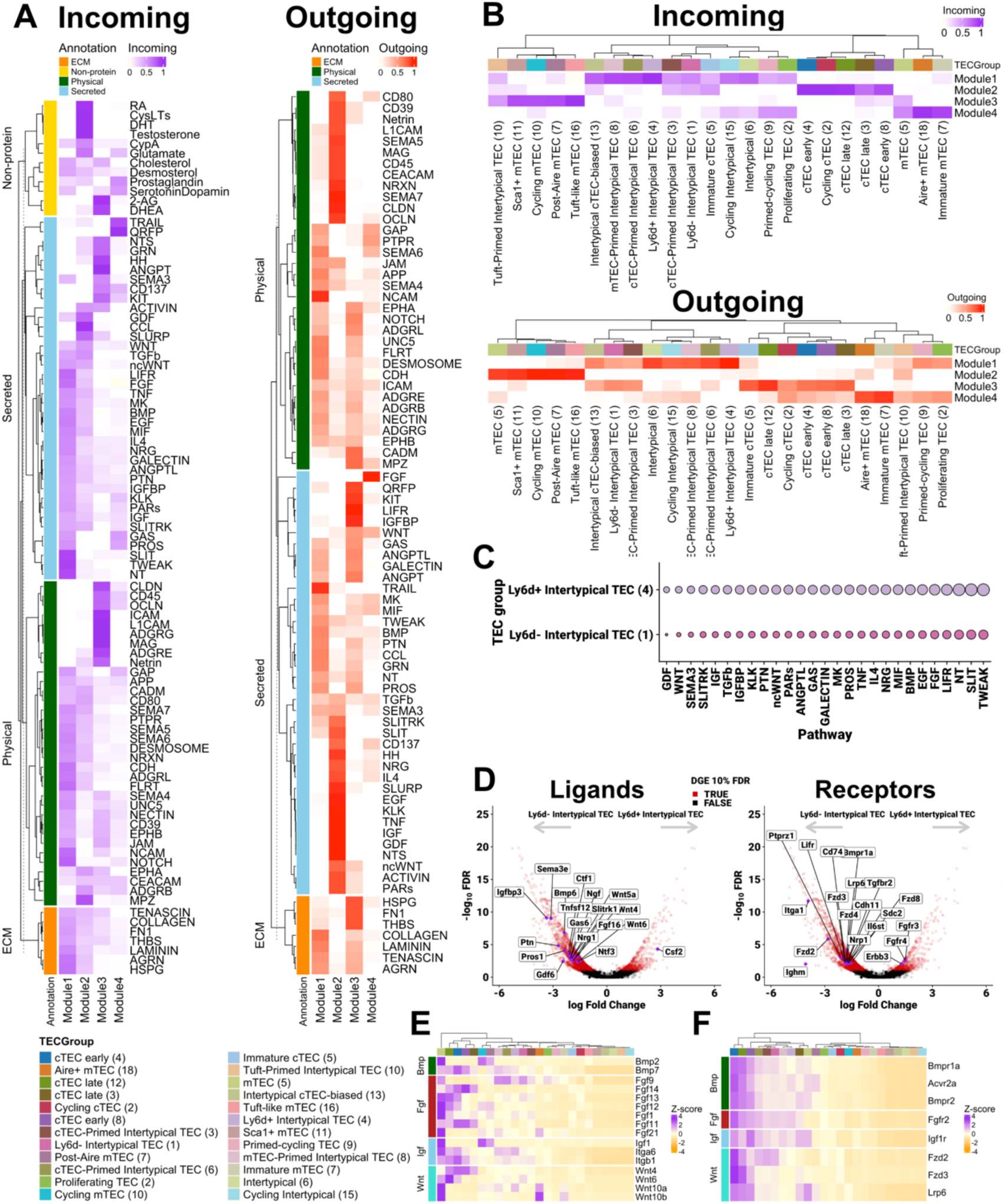
Cell-cell communication identifies TEC progenitor signalling cues. **(A)** Heatmap of NMF weights for each signalling pathway on coordinated modules. Each row is a CellChat signalling pathway for one of 4 identified modules (columns). Each pathway is annotated based on mechanism of action; either physical interaction (dark green), secreted paracrine (light blue) or ECM interactions (orange). **(B)** Weighting of each TEC state on the coordinated signalling modules as in (A). **(C)** Module 1 specific signalling weights for Ly6d+ and Lyd6-intertypical TEC. **(D)** Differential gene expression between Ly6d+ and Ly6d-intertypical TEC highlights differential signalling cues. **(E-F)** Gene expression levels (Z-scores; y-axis) of DE signalling components shown as violin plots. Each row is a DE ligand (E) or receptor (F) for Wnt, Fgf or Igf signalling receptor for TEC states (x-axis). The colour legend denotes the annotated TEC subtypes and corresponds to panels B, E & F.

Focussing first on the input signal module 1 which had the highest score in Ly6d+ and Ly6d-intertypical TEC, the ligand-receptor interactions with the greatest weighting included *Tnfsf12*-*Tnfrsf12a* (encoding Tweak-TweakR), multiple Notch-Jag/Dll pairs, *Slit2*-*Robo1/2*, *Ntf5*-*Ngfr*, and *Lif*-*Lifr* (Figure 4A-C); *Lifr* is a marker gene of intertypical TEC (Figure 2G). In addition, *Pros1* and *Gas6* share the receptor, *Axl*, and had high contributions to incoming signal module 1 and 4. The latter module was enriched in the mTEC lineage, suggesting these pathways may be involved in mTEC vs. cTEC fate choices, or their lineage maintenance. Moreover, TNF signalling (*Tnf*-*Tnfrsf1a*), which is known to promote TEC fate choices, was also strongly represented in module 1. The pattern of *Tnfrsf1a* expression, which encodes TNF receptor 1 (TNFR-I), was widespread across TEC subtypes, with lowest levels in mature, Tuft-like and Aire+ mTEC, and highest levels in Intertypical cTEC-biased and Ly6d-intertypical TEC. *Tnfrsf1a* expression in Ly6d+ Intertypical TEC was intermediate between Ly6d-Intertypical and mature mTEC (Supplementary Figure 19C).

Module 2 was also enriched amongst Ly6d-intertypical TEC and cTEC states, suggesting common signalling inputs. The ligand-receptor pairs contributing most to this module included testosterone, dihydrotestosterone, cysteinyl leukotrienes, retinoic acid, and *Ccl19/21/25-Ackr4*. Our single-cell data were derived from both male and female mice and are balanced across cell types or experimental conditions (Supplementary Figure 20). Therefore, the observation that sex hormone signalling, which is a known early contributor to thymic involution^62^, appears to differentially signal between TEC subtypes is interesting. Indeed, there is a rapid turnover of cTEC early in mouse life^46^, which we have confirmed in our experiments (Figure 2A-B, Supplementary Figure 8) and may therefore hint at a mechanism by which castration induces transient thymic renewal in male mice^63^.

Finally, incoming signal modules 3 and 4 were strongest in multiple mTEC subtypes, further illustrating the separation of cell-cell communication pathways between TEC lineages (Figure 4B). The top 5 ligand-receptor interactions on module 3 involved adhesion molecules L1 adhesion molecule (L1CAM), G protein coupled receptor G1 (ADRG) and intercellular adhesion molecule 1 (ICAM1), as well as myelin associated glycoprotein (MAG) and angiopoietin 1 (ANGPT1; Figure 4A). In contrast, the top 5 incoming signals on module 4 were the neuropeptide and receptor *Qrfp-Qrfpr*, Trail-Trail receptor, prostaglandin via *Ptge3*-*Ptger3/4*, *Gas*-*Axl*, and serotonin/dopamine reception via *Htr1b* and *Htr7*. The highest expression for the receptors for these 5 paracrine and endocrine signals was in the Aire+ mTEC (Supplementary Figure 20A-E) suggesting they are either involved in mTEC maturation, or their expression is a consequence of promiscuous gene expression. To explore the latter, we reasoned that promiscuous gene expression would manifest as highly variable expression as opposed to constitutive expression within Aire+ mTEC. We compared the proportion of cells expressing top ligand-receptor genes from module 4 against a set of control mTEC genes (*Aire, Cd52, Epcam, Krt10, Fezf2, Ascl1*) and Aire-dependent peripheral tissue antigen genes from Sansom *et al*^42^. Our analysis showed that while ligand and receptor genes were expressed heterogeneously across Aire+, mature and post-Aire mTEC (Supplementary Figure 20F), they were detected in a much higher proportion of cells than would be expected based on promiscuous expression of peripheral tissue antigen genes.

In contrast to signal reception, we also examined the outgoing signalling from each TEC subtype, identifying a further 4 outgoing modules (Figure 4A-B). Like the incoming signals, we observed a separation of TEC subtypes by outgoing signalling modules. For instance, the outgoing module 1 which was enriched amongst intertypical TEC subtypes, independent of Ly6d expression, was strongly associated with TRAIL signalling (*Tnfs10*-*Tnfrsf10b;* Figure 4A). The corresponding incoming signal was strongest amongst mTEC subtypes. TRAIL induces apoptosis upon binding to the TRAIL receptor, and a high rate of apoptosis has been described for immature mTEC as a potential quality control checkpoint during mTEC maturation^55^. These findings suggest that progenitor TEC may signal to immature mTEC to regulate mature mTEC abundance.

To prioritise signalling cues important for TEC progenitor biology we focussed on the pathways with both strong incoming signals to the Ly6d+ and Ly6d-intertypical TEC from module 1 (Figure 4C) and that were differentially expressed between Ly6d+ and Ly6d-intertypical TEC (Figure 4D-E). Notably, Wnt receptors *Fzd2, Fzd3, Fzd4* and *Lrp6* were up-regulated in Ly6d-intertypical TEC. Likewise, Wnt ligand genes *Wnt4, Wnt5a* and *Wnt6*, implicating both canonical and non-canonical Wnt pathways were up-regulated in Ly6d-intertypical TEC. Wnt4 regulates the thymus master regulator Foxn1, which is required for embryonic TEC development^58^, early postnatal mTEC expansion^57^ and adult thymus maintenance and function^64^.

In contrast, Ly6d+ intertypical TEC up-regulated the genes for fibroblast growth factor receptor 3 (*Fgfr3)* and 4 (*Fgfr4)* which bind fibroblast growth factors (Fgf) 8/9^65^ and 1^66^, respectively (Figure 4F). Fgf signalling is required for thymus organogenesis^67–69^ and has been implicated in TEC maintenance^70^. Moreover, keratinocyte growth factor (KGF, also known as FGF7), along with Fgf10 binds to Fgfr2IIIb and induces transient thymus rejuvenation^70–73^. *Fgfr2* is expressed broadly across intertypical TEC, cTEC and immature mTEC subtypes (Figure 4F), suggesting that KGF treatment acts on multiple TEC subtypes. Additionally, we observed that IGF signalling components are more highly expressed in Ly6d+ intertypical TEC, including potential trans-activation between *Igf1r* and *Itga6*^74^ (Figure 4E-F). *Itga6* is a marker of previously identified TEPC^21^, supporting the notion that Ly6d+ intertypical TEC represent the progenitors previously described. Together, our analysis of the paracrine and autocrine signalling landscape across perinatal TEC subtypes highlights candidate signalling pathways that might separate the biological fates of Ly6d+ vs. Ly6d-intertypical TEC.

### Ly6d-intertypical TEC are more affected by ageing than Ly6d+ intertypical TEC

We have previously described an age-dependent accumulation of intertypical TEC^12^. Therefore, we sought to understand the dynamics of our newly identified intertypical progenitor TEC across the mouse life course. First, we matched neighbourhoods between our perinatal and ageing datasets based on the correlation of their transcriptomes (Supplementary Figure 21A). We then performed differential abundance testing to compare cells from younger to older mice separately on the ZsGreen+ and ZsGreen-ageing neighbourhoods, which identified 1132 and 640 DA neighbourhoods, respectively (10% FDR). This analysis showed that the abundance of both labelled and unlabelled cells are altered by ageing (Supplementary Figure 21B,C). However, when focussing on specific TEC subtypes we found that Ly6d-intertypical TEC subsets were more depleted than the Ly6d+ intertypical TEC subsets in both the ZsGreen+ and ZsGreen-fractions (Figure 5A, Supplementary Figure 21D-F, p=5.73x10^-28^, Mann-Whitney test), suggesting that Ly6d-intertypical TEC might be more sensitive to the effects of ageing. We additionally observed in older thymi a depletion along both the cTEC (Immature cTEC, cTEC early, cTEC late, Cycling cTEC) and mTEC lineages (Sca1+ mTEC, Aire+ mTEC, mTEC) as reported previously.

**Figure 5:**
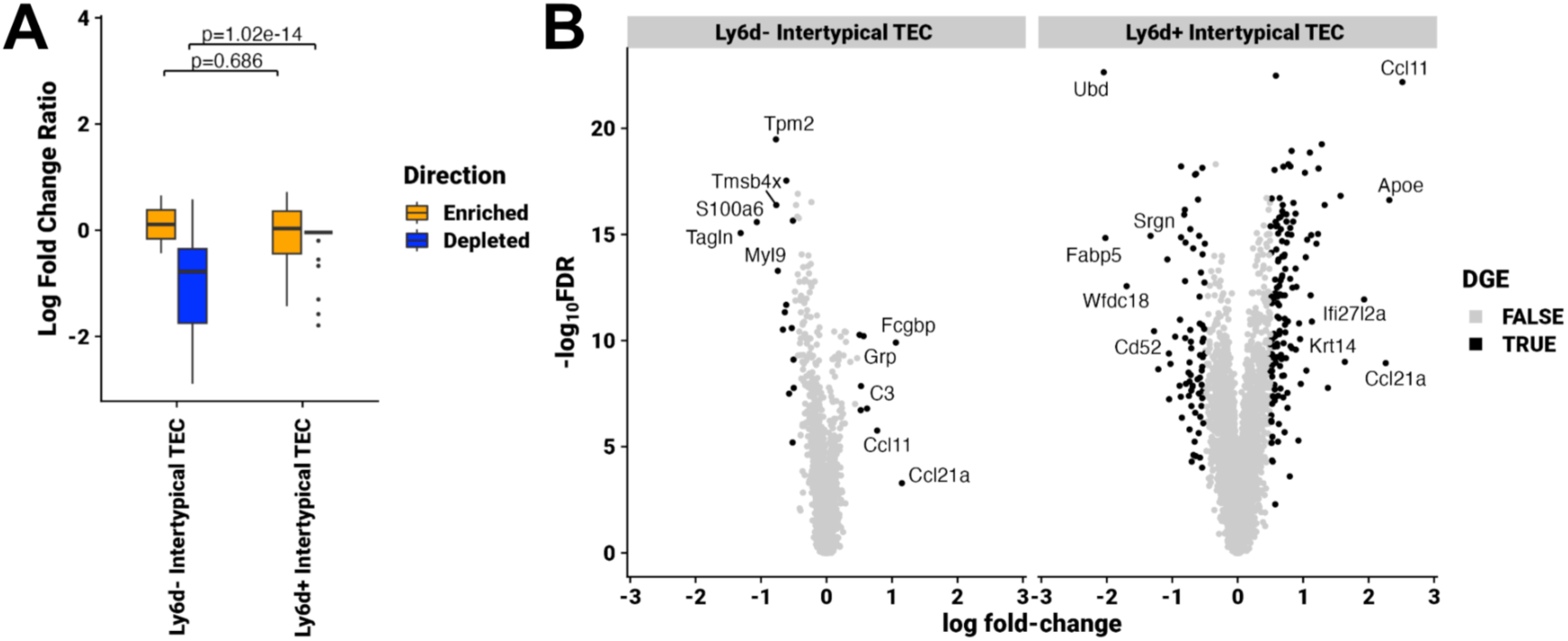
(A) Differential abundance log-fold change ratio between ZsG+ and ZsG-cells across the ageing time course. Positive values denote a greater change in the ZsG+ and vice versa. Box plots are coloured by whether the neighbourhoods are concordantly enriched (orange), depleted (blue) or are discordantly DA (orange). Boxes show the median and box limits are the lower and upper quartiles. Whiskers extend to 1.5x the IQR with outliers beyond this range shown as individual points. (B) Volcano plots denoting DEGs across the mouse life course. Points are coloured black with FDR<0.1 or not DE (grey).

To understand the gene expression differences between younger and older intertypical TEC we compared the transcriptomes of more vs. less abundant Ly6d+ and Ly6d-intertypical TEC separately, which identified 183 and 437 differentially expressed genes (DEGs), respectively (Figure 5B, Supplementary Figure 22, 1% FDR). Co-ordinately up-regulated genes in more abundant neighbourhoods of both intertypical TEC subtypes include *Nfib, Txnip* and *Ackr4*; the latter sequesters *Ccl21a*^75,76^, the canonical marker for intertypical TEC, while *Nfib* serves as an additional marker identifying Ly6d-intertypical TEC (Supplementary Figure 13B). Genes expressed in the less abundant neighbourhoods were enriched for antigen presentation, including MHC II and which are generally up-regulated during TEC differentiation, in line with a restriction in lineage differentiation^12,44^. Gene set enrichment analysis (Supplementary Figure 23) identified several cellular gene expression programs affected by ageing, including cytokine signalling (IL6/JAK/STAT signalling, Interferon alpha, Interferon gamma, TNF alpha), response to mitogen and cell division (E2F targets, G2-M checkpoint, Myc targets) and cell-cell communication (TGF-beta signalling, Wnt-beta Catenin). Based on Mohammed *et al.*^52^, the disruption of both type I and II interferon signalling in older thymi may contribute to the decline in TEC maintenance due to diminished haematopoietic cell input into the involuting thymus. Collectively our results demonstrate the wide-ranging impact of ageing on TEPC in-keeping with inflammageing, replicative senescence, and remodelling of paracrine signalling.

In sum, our analysis of TEPC, across the mouse life course has identified the differential impact that ageing has on Ly6d+ and Ly6d-putative progenitors providing molecular evidence for the disproportionate loss of mTEC during ageing and thymic involution.

## Discussion

TEC development and the search for thymic progenitor cell populations has attracted considerable attention in recent years due to its potential to unlock thymic rejuvenation. However, there is little consensus with regards to the identity and developmental potential of postnatal TEPC. Here, we have identified and quantified the contributions of β5t+ TEPC to postnatal TEC lineages in the mouse combining genetic lineage tracing with flow cytometry and single-cell transcriptomics. Our experiments reveal over the course of the first 8 weeks of life the disproportionate accumulation of lineage-trace labelled intertypical TEC compared to cTEC and mTEC (Figure 1-2). These results suggest an early lineage bias switch in β5t+ TEPC from foetal cTEC to postnatal mTEC. Supporting this conclusion, we resolve the identity of not one, but two postnatal TEC progenitors, each distinguished by the expression of the surface marker Ly6d and their separate lineage biases (Figure 3). Our transcriptome and spatial analyses of these cells extends to the impact of ageing on each, in which we discover that Ly6d-TEPC are more affected than Ly6d+ TEPC, leading to their greater depletion over 32 weeks of mouse life; a period that encompasses the rise and fall of thymic epithelium during involution^12^.

We and others have previously shown that adult mTEC are derived from both foetal and newborn-derived TEC progenitors expressing the cTEC-specific protein β5t^18,30^. Additionally, ageing reduces the efficiency with which postnatal β5t+ TEPC contribute to the maintenance of the thymus medulla^18,30^. However, the phenotypic similarity between cTEC-like progenitor cells and other *bona fide* cTEC populations has limited the isolation and characterisation of postnatal TEPC that contribute to the medullary linage^36^. Recently, we described intertypical TEC as a postnatal precursor to mature mTEC that arise from or contain β5t-expressing TEPCs^12^. Intertypical TEC share marker genes with previously described TEPC populations, such as Ccl21a, Pdpn and Sca1^19,20,29,44^. In this study, we utilised intra-cellular Ccl21 and surface Sca1 staining to flow cytometrically quantify the dynamics and contribution of β5t+ TEPC to intertypical TEC, cTEC and mTEC subpopulations.

Detailed analysis of the labelling kinetics of cTEC, mTEC and intertypical TEC showed a strong accumulation of the ZsGreen label in the intertypical and mTEC compartments, while the proportion of ZsGreen+ cTEC decreased with age, indicating (a) a fast turnover of foetal-to-postnatal cTEC^22^ and (b) an increased contribution from β5t-low or negative progenitors postnatally (Figure 1G). The constitutive expression of *Psmb11* during cTEC maturation and maintenance ^16,77^ prevents a definitive analysis using our transgenic system. However, using single-cell RNA-sequencing of ZsGreen+ and ZsGreen-sorted cells we observed the same dynamics up to 10 days post-dox treatment of 1 week old pups, corroborating our findings.

Our model of *Psmb11*-dependent lineage tracing has the best discriminative power to discover progenitor-progeny relations for lineages that lack constitutive β5t expression yet are derived from *Psmb11* expressing precursors, such as mTEC and intertypical TEC. Indeed, we observed a steady increase in ZsGreen+ mTEC across multiple developmental stages, which plateaus 4 weeks after Dox treatment. These results indicate β5t+ TEPC contribute equally to intermediate and mature mTEC subpopulations. We also observed a striking accumulation of ZsGreen signal in intertypical TEC, equivalent to a 50-fold increase over 9 weeks of lineage tracing (Figure 2). More detailed flow cytometry analysis suggests these accumulating cells are mTEC-biassed as the relative representation of ZsGreen+ cTEC biassed intertypical TEC remained constant (Figure 2C). To explore these dynamics further, we applied single-cell RNA-sequencing to our system, focused on the earliest time points with the greatest changes across TEC compartments.

Our single-cell transcriptome analysis identified 2 putative TEPC populations, delineated by the differential expression of *Ly6d*. Surface expression of Ly6D marks a number of progenitor populations in haematopoietic (plasmacytoid dendritic cells, lineage-restricted common lymphoid progenitors & thymocytes)^78–80^ and stromal cell types (prostate luminal cells)^81^. Using bioinformatic analysis and differential abundance testing, we found that Ly6d-intertypical TEC accumulated in the ZsGreen+ fraction over time. In contrast, we observed no such bias for Ly6d+ intertypical TEC (Figure 3). These analyses suggest that the latter subtype may precede a β5t+ state, and which would either place Ly6d+ intertypical TEC as the earliest postnatal TEPC identified to date, or as a parallel lineage that arise from a β5t-TEPC. However, we observed a small fraction of ZsGreen+ Ly6d+ intertypical TEC, suggesting these cells can be labelled. To explore this further, we found this small proportion of Ly6d+ intertypical TEC express Foxn1 and Psmb11 transcripts which can lead to the activation of our transgene after Dox treatment. Therefore, the balance of evidence indicates that Ly6d+/- intertypical TEC contains the previously described β5t+ TEPC. Further lineage-tracing studies will be required to validate that these two TEC subtypes are able to give rise to mature postnatal cTEC and mTEC, and to establish the relationship with each other.

We have previously described the acquisition of an inflammaging-like transcriptional signature in intertypical TEC, which we hypothesised might explain their reduced contribution to the mTEC compartment across the mouse lifecourse^12^. We further explored the impact of ageing on our putative TEPC progenitors, and found that Ly6d-intertypical TEC are more strongly depleted across the mouse life course than Ly6d+ intertypical TEC. The down-regulated Ly6d+/- intertypical TEC expressed genes were indicative of TEC maturation, adding weight to previous observations that thymic involution is due to the restricted differentiation of progenitors into mature TEC^12,18,30,44^. These results are important for strategies that seek to rejuvenate the ageing thymus, as they point to the dysfunction that requires correcting to restore the thymic epithelium and thus rejuvenate T cell development. Our results indicate these should target inflammation (e.g. IL6/JAK/STAT signalling), mitogen response (e.g. Myc targets) and cell-cell communication (e.g. TGF-β and Wnt - β-catenin signalling). Current experimental models to reinvigorate aged thymic cellularity include androgen depletion^63,82,83^, Foxn1-overexpression^84,85^ or growth factor treatment^67,70–73^. The latter notably uses KGF (FGF7) which binds to Fgfr2-IIIb, and acts as a mitogen that causes transient thymus growth and increased cellularity^67,70–73^. Our data indicate that growth factor treatment may have wide-ranging effects on cTEC, immature mTEC and intertypical TEC; whether the transient rejuvenation is caused by depleting the intertypical TEC is not clear and requires further investigation before KGF treatment can become a viable option for thymus renewal.

Our study makes extensive use of the 3xtg^β5t^ transgenic model for lineage tracing, and single-cell RNA-sequencing. This assumes that ZsGreen+ labelling is efficient and specific. Here we show that efficient, yet differential, labelling is already achieved with low doses of dox-treatment - a feature we use to our advantage to label maturing mTEC. However, the labelling of cells with no detectable *Psmb11* mRNA indicates that (i) scRNA-seq is limited at capturing very low expression levels and/or (ii) promoter accessibility without mRNA may be sufficient to permanently label cells. Indeed, we observed labelled cells without *Psmb11* transcripts that correlated with the expression of *Foxn1*, the main regulator of *Psmb11* in TEC. Therefore, promoter accessibility may prime cells for the cTEC lineage without them irreversibly committing to it, restricting the 3xtg^β5t^ model.

Finally, a true test of a bipotent TEPC is the ability of a single cell to give rise to both cTEC and mTEC lineages, given the right differentiation cues. Our experimental, single-cell and spatial data all point to the presence of 2 unipotent or lineage-biassed TEPC, though a progenitor-progeny relationship between these TEC subtypes could not be firmly excluded. In sum, our detailed molecular and cellular characterisation of Ly6d+ and Ly6d-intertypical TEC now enables their further study to resolve their role in thymic maintenance across the life course.

## Methods

### Mice

3xtg^b5t^ mice [β5t-rtTA::LC1-Cre::CAG-loxP-STOP-loxP-ZsGreen] mice have been previously described^18^. Animals of both sexes were used. Mice were kept under specific pathogen-free conditions, with food and water *ad libitum,* at a temperature of 22 ± 2 °C, 60 ± 15% humidity, and a 12 h light cycle with light phase beginning at 6 a.m. with a 30 min sunrise and ending at 6 p.m. with a 30 min sunset. Both the caring for the mice and the experiments were performed in compliance with permissions and regulations of the Cantonal Veterinary Office of Basel-Stadt (licence 2321).

### Doxycycline treatment

7-day old pups were treated with a single i.p. injection if decreasing doses of doxycycline (0.3 mg, 0.02mg, 0.004mg or 0.0008mg) diluted in 100 μl of sterile HBSS. Mice older than 4 weeks were treated with two i.p. injections of doxycycline (2mg, each) diluted in 200 μl of sterile HBSS. In the course of 24 h in between Dox injections, the mice were also exposed to drinking water supplemented with the drug (2 mg/mL in sucrose (5% w/v))^18^.

### Thymic epithelial cell isolation

Fatty and connective tissue was cleaned off the isolated thymi. Thymic lobes were cut into 0.1-0.2 cm cuboids using a scalpel and incubated either with Liberase^TM^ and DNaseI (Roche Diagnostics; 200 and 30 μg/ml, respectively; for 45 min at 37 °C)^86^ or with Papain (Sigma Aldrich), Liberase^TM^ and DNaseI (500, 200 and 30 μg/ml, respectively; for 30 min, at 37°C)^87^ to obtain a single cell suspension. Tissue digestion was mechanically aided by repeated pipetting of the cell suspensions at 15-minute intervals. Finally, cell suspensions were washed with Iscove’s Modified Dulbecco’s Medium (IMDM) (Life Technologies) containing 10% Foetal Calf Serum (FCS) (HyClone, Thermo Fisher Scientific) and 2mM EDTA (Invitrogen) to stop the enzymatic reaction and filtered through a nylon mesh (70 μm pore size, Sefar Nitex) to remove debris. Cells were pelleted by centrifugation and resuspended in PBS containing 2% v/v heat-inactivated FCS (designated FACS buffer) for downstream analysis.

### Flow Cytometry

Whole thymus or spleen single cell suspensions were typically stained with fluorochrome-conjugated antibodies against the desired surface epitopes for 30 minutes, at 4°C, in the dark. When required, cells were washed with PBS and stained with the appropriate Zombie^TM^ fixable viability dye (Biolegend) for 30 minutes at 4°C prior to fixation. Fixation and permeabilization for intracellular staining were performed with either Cytofix/Cytoperm kit (BD Biosciences) or eBioscience transcription factor staining buffer set (ThermoFisher Scientific), according to the manufacturer’s guidelines. Specific antibodies and reagents, as well as the optimal conditions for their use (including dilutions), can be found in Supplementary Table 1.

### TEC sorting

Whole thymus cell suspensions were generated by enzymatic digestion with Papain, Liberase^TM^ and DNaseI as described above. Cell suspensions were either enriched for EpCAM positivity or depleted of CD45+ cells prior to FAC-sorting using an automatic magnetic cell separator (AutoMACS Pro Separator, Miltenyi). Enriched cells were incubated for 30 minutes at 4 °C in FACS buffer containing the required fluorescently labelled antibodies for the sort. After washing, cells were resuspended in FACS buffer and filtered through a 40 μm mesh into tubes containing DAPI (Invitrogen) for dead cell exclusion. Cells were sorted using a FACSAria II (BD Bioscience) into 1.5 ml Eppendorf tubes containing the most appropriate buffer for the desired downstream application.

### Droplet-based single cell RNA sequencing

#### TEC processing and sorting

Whole thymus single cell suspensions from thymi isolated either 2 days or 10 days post treatment with 0.004mg of Doxycycline were obtained by enzymatic digestion using Papain, Liberase and DNaseI and enriched for TEC by EpCAM positivity, as described above.

TotalSeq-A oligonucleotide-conjugated antibodies (BioLegend) were used to allow for barcoding and pooling of different TEC subpopulations, as well as to assess surface protein levels of different markers (Supplementary Table 2). Enriched cells were simultaneously stained with the required fluorescently labelled antibodies for the TEC sort as well as with the TotalSeq-A antibodies. TEC were subsequently sorted into four different subpopulations: ZsGreen+ Ly51+ UEA1-cTEC, ZsGreen- Ly51+ UEA1-cTEC, ZsGreen+ Ly51-UEA1+ mTEC, ZsGreen-Ly51- UEA1+ mTEC. Post-sorting cell viability and concentration in each of the different samples was determined using a Nexcelom Bioscience Cellometer K2 Fluorescent Viability Cell Counter (Nexcelom Bioscience).

#### Library generation

Each well of a Chromium Single Cell B Chip (10X Genomics) was loaded with a total of 24,000 cells, pooled equally from each of the four sorted samples. The fact that the pooled samples had been barcoded with four different TotalSeq-A hashtag antibodies allowed to overload the 10x wells with 24,000 cells per well, aiming for a recovery of approximately 10,000 single cells (∼40%) per well, and enabling to overcome the increase in doublet rate by subsequently eliminating any cell barcode containing more than one single hashtag sequence from further analysis, as described in^12^. The Chromium Single Cell 3’ GEM, Library and Gel Bead Kit v3 and Chromium i7 Multiplex Kit (10X Genomics) was used for library preparation, according to the manufacturer’s instructions. In short, well number 1 on the Chromium Chip B was loaded with a mixture of the cell suspension with the GEM Retrotranscription Master Mix. Wells 2 and 3 were loaded with gel beads and partitioning oil, respectively. Nanoliter-scale Gel Beads-in-emulsion (GEMs) containing the single cells to be analysed were subsequently generated using the Chromium Controller (10x Genomics). Inside of each individual GEM occurs the simultaneous production of barcoded full-length cDNA from poly-adenylated mRNA as well as barcoded DNA from the cell surface protein-bound TotalSeqA antibodies. Silane magnetic beads were used for the recovery and clean-up of the pooled fractions following the fragmentation of the GEMs, after which the recovered DNA was amplified and cDNA products, Antibody-Derived Tags (ADT) and Hashtag oligonucleotides (HTO) were separated by size selection. To optimise the amplicon size for the generation of the 3’ libraries, the full-length cDNA generated from polyadenylated mRNA was enzymatically fragmented and subjected to size selection. Subsequently, addition of P5, P7, a sample index, and TruSeq Read 2 (read 2 primer sequence) via End Repair, A-tailing, Adaptor Ligation, and PCR were performed to construct the libraries to be sequenced. In parallel, generation of the ADT and HTO libraries involved the addition of P5, P7, a sample index, and TruSeq Read 2 (read 2 primer sequence) by PCR. The sequences of the oligonucleotides and primers employed for this purpose can be found in Supplementary Table 3.

#### Library quality control, pooling and sequencing

Capillary electrophoresis on a Fragment Analyser (AATI) was used to assess the quality of the libraries generated prior to sequencing. The three sets of libraries generated were pooled as follows: 85% cDNA + 10% ADT + 5% HTO, and the pooled libraries were sequenced using the NovaSeq 6000 S2 Reagent Kit (100 cycles) on an Illumina NovaSeq 6000 sequencing system (Illumina).

#### Single cell RNA-sequencing quality control and processing

CITE-seq libraries were processed using Cellranger v3.10 using mouse genome build GRCm38. Non-empty droplets were called using *emptyDrops*^88^ on each multiplexed sample with 20,000 permutations, a lower UMI threshold of 100, and 1% FDR. Across all sequenced library samples, 101,255 non-empty droplets were defined (Supplementary Figure 6). We defined low-quality single cells as those with <1000 UMIs, and excessive mitochondrial RNA content (Supplementary Figure 6). The latter was defined based on departure from a normal distribution with mean set to the median mitochondrial RNA proportion of the sample and standard deviation as twice the median absolute deviation. Excessive mitochondrial RNA content cells were then defined as those with a 1-tailed FDR adjusted p < 0.05; 2627 cells were excluded from downstream analysis. HTO demultiplexing was performed as described in^89^. Specifically, for each sequenced library, HTO UMI counts were scaled by HTO sequencing depth and used as input to k-means clusters, with k set to the number of expected clusters, i.e. the number of combinations of single and doublet HTO. For each HTO, we first assigned the candidate k-means cluster as the one with the highest normalised counts over cells. Excluding these cells and the top 5% of cells across other clusters, we then estimated a background distribution of counts by fitting a negative binomial with the *fitdist* function from the *fitdistrplus* package to the counts across the remaining cells. To assign cells to the HTO, we selected those with UMI count > 99th percentile. After repeating this for each HTO, cells were assigned to either singlet (a single HTO), multiplet (>1 HTO) or dropout (0 HTOs). For all downstream analyses we excluded multiplet and dropout HTOs, leaving 64,500 high-quality cells.

#### Single cell normalisation, clustering and visualisation

Single-cell gene counts were combined across all samples and were rescaled using pooled-estimate size factors prior to log + 1 transformation, as implemented in *scran*^90^. Highly variable genes (HVGs) were estimated by modelling the mean-variance relationship with *modelGeneVar* in the scran package, with a 10% FDR threshold - a total of 3327 HVGs were identified. Centred and scaled normalised gene counts for HVGs were used as input to randomised PCA, implemented in the R package *irlba*. Uniform manifold approximation and project (UMAP) coordinates were computed using 50 PC dimensions as input with 30 neighbours, min_dist=0.2 and random initialised (seed=42) as implemented in the *umap* R package. Single-cell clusters were calculated from a k-NN graph computed using 30 PC dimensions and k=21 using the Walktrap algorithm implemented in *igraph*^91,92^.

#### Differential abundance and neighbourhood analysis

*Milo* was used to build k-nearest neighbour graphs, construct refined neighbourhoods (k=50, props=0.05) using the graph-only^48^ method, and perform differential abundance testing.

Neighbourhood counts were modelled using the *Milo* negative binomial generalised linear model (GLM), with an FDR 10% threshold, unless otherwise stated. Neighbourhood clusters were defined by first constructing an adjacency matrix defined by an overlap threshold (overlap=10), where an edge is drawn between pairs of neighbourhoods sharing ≥ *overlap* numbers of cells. Louvain clustering was then performed on the corresponding graph using *igraph*^87^. Neighbourhood group marker genes were identified using the *findNhoodGroupMarkers* function in *Milo* with a 10% FDR threshold.

#### Cell-cell communication analysis

We inferred cell-cell communication between TEC subtypes using *CellChat*^93^. We used the mouse CellChat DB to annotate expressed ligand and receptor genes across single cells. Gene expression data were averaged over TEC subtype labels using the trimean function, and clusters with fewer than 10 cells were excluded from analysis. Significantly communicating pathways were defined based on a network out degree > 10^-4^, which yielded 91 active signalling pathways in total. To estimate co-ordinated signalling modules, we used the cophenetic and silhouette indices to select a k value for subsequent non-negative matrix factorisation (NMF), with k=4 used for both incoming and outgoing signalling module estimation. NMF factors were visualised as heatmaps using the ComplexHeatmap package^94^.

#### Cross-study integration and neighbourhood mapping

Gene counts data were downloaded for Baran-Gale *et al.*^12^, Ragazzini *et al.*^51^ and Wells *et al*.^50^ and normalised gene expression data as described above. For Ragazzini *et al*. we further used the *multiBatchNorm* function from the Bioconductor package *batchelor* to re-normalise across batches. HVGs were computed on each (Ragazzini: 8093, Wells: 2111, Baran-Gale: 3614) and used as input for PCA using *multiBatchPCA* in the *batchelor* package. For Ragazzini *et al*. and Wells *et al.* data we also integrated cosine-normalised principle components across study batches using mutual nearest neighbours^95^ (k=50 and k=21, respectively). kNN-graphs were computed for each data set (Ragazzini k=50, Wells k=21, Baran-Gale k=50). We then computed refined neighbourhoods using *Milo*^47,48^ with the graph-only algorithm. To identify cross-mapping neighbourhoods, we computed the Pearson correlation coefficient between neighbourhood pairs, one each from our perinatal data and the corresponding study. We then identified the best matching neighbourhood pairs as those with the highest Pearson correlation. Cross-mapping between TEC subtypes was visualised using *ggalluvial*. For human to mouse cross-mapping, we first identified the 1-to-1 homologous genes from Ensembl, accessed through the Bioconductor package *biomaRt*^96^.

#### Differential gene expression analysis

Single cell gene counts were aggregated over groups, e.g. TEC subgroups, neighbourhood clusters, etc, for each replicate sample in each condition to generate pseudo-bulk counts. Differential expression testing was then performed by modelling pseudobulk gene expression counts using a negative binomial GLM, implemented in *edgeR*^97,98^. Differentially expressed genes were defined based on statistical significance with a 10% false discovery threshold, unless otherwise stated. Gene set enrichment analysis was performed on up- and down-regulated genes separately using the *enrichR* package^99^. MSigDB Hallmark gene sets were used as input. Statistically significantly enriched/depleted gene sets were defined with a 10% FDR threshold. DGE analysis results were visualised using *ggplot*.

### Spatial transcriptomic profiling

A thymic lobe from an 8-week-old female C57BL/6 wildtype mouse was fixed in 10% neutral buffered formalin (Sigma Aldrich) at 4°C for 24 hours and then embedded in paraffin.

Sections at 5µm thickness were loaded onto a Xenium slide (10x Genomics) and processed according to the manufacturer’s instructions for spatial transcriptomics using the Xenium Prime 5k Mouse Pan Tissue and Pathways Panel and the Cell Segmentation Staining kit (10x Genomics).

#### Spatial transcriptomics analysis

Cell types were initially annotated using label transfer from Steier et al.^100^ before manually curating the final cell types to match the unsupervised clustering. The annotations were split into a preliminary cortical (DP (Q1), DP (Q2), DP (Sig.), DP (P), DN, cTEC, tc_stromal_mix(cor), CapFb, Adipo) and a medullary compartment (mTEC, TlTEC, MedFb, Mature CD4, tc_stromal_mix(med), SP4_resting / Treg-like) before progressing with the nearest-neighbour-based smoothening approaches (see below) to arrive at the final compartment definition.

#### Spatial transcriptome processing and analysis

The experiment was transformed into a Seurat object^101^, filtered for cells with more than 50 and less than 600 unique features as well as cells with more than 100 and less than 900 total transcripts and subjected to Seurat’s *SCTransform* ^102^. *Lifr* and *Ccl21a* double-positive stromal cells (those that have non-zero raw counts for both these genes) were retained as intertypical TEC. To remove potential contamination from imperfect segmentation and transcript contamination from neighbouring immune cells we further filtered cells with non-zero expression for at least 4 of the following genes: *Ptprc, Cd8a, Cd8b1, Cd4, Cd3e, Cd3g, Rag1, Cd5, Cd69, Cd68, Sirpa, Lck, Tcf7, Gzma, Ighm, Ighd, Itgax*.

Next, we took the intersection of the Ly6d+ and Ly6d-intertypical TEC marker genes (Supplementary Table 4) with the genes covered by the Xenium 5k panel. Module scores for both gene sets were computed per cell using the Seurat *AddModuleScore* function^103^ and cell-wise subtracted from each other to create a joint score. The difference between module scores was normalized to the largest absolute number. Ly6d+ cells were classified based on a score > 0.1 and Ly6d-intertypical as a score < -0.1; all other cells were labelled as “Unclassified”.

#### Thymic spatial compartment definition

A region definition for the thymus tissue section was developed exploiting the location of well-known cortical and medullary cell types (Supplementary Figure 24). To obtain a global, cortex/medulla definition and overcome the mispositioning of few cortical cell types scattered in the medulla and vice versa, a nearest neighbour network (FNN package v1.1.4.1^104^) was used to smoothen out inaccuracies in initial compartment allocation. This was done by querying the closest 11 cells to each other cell for their initial compartment annotation.

Subsequently, the compartment annotation of the closest 501 cells was used to filter out small medullary islands. From this final cortex and medulla definition, we assigned cortical cells to the CMJ as those within 50um of a medullary cell. To compute the density of cells in each of the cortex, medulla and CMJ we counted the number of cells of each type (Ly6d+ and Ly6d-intertypical TEC) and scaled these based on the total cell area of each compartment. To calculate the total cell area for each histological compartment (cortex, CMJ, medulla), the shortest distance from each cell centroid to the nearest neighbour cell was computed in each compartment separately and then averaged. This average shortest distance served as an estimate of the average diameter of each cell in each compartment. Next, we assumed hexagonal packing of cells and estimated the area using the following formula: 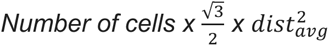, *where* 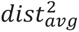 is the average squared physical distance between cell centroids.

## Code & Data availability

Analytical code can be accessed at https://github.com/MorganResearchLab/ProgenitorTEC2024. All single-cell RNA-sequencing data are publicly available at ArrayExpress (E-MTAB-15523), and Xenium spatial transcriptomics are accessible from BioImage archive (S-BSST2125). Publicly available scRNA-seq data were downloaded from GEO for Wells *et al.* (GSE137699) and Ragazzini *et al.* (GSE220830).

## Author contributions

ICA – conceptualization, formal analysis, investigation, methodology, validation, writing – original draft, writing – review & editing.

AT – formal analysis, data curation, investigation, validation, writing – review & editing.

FD – formal analysis, investigation, validation, writing – review & editing.

AK – investigation. TB – investigation. SZ – investigation.

GH – conceptualization, funding acquisition, resources, project administration, supervision, writing – original draft, writing – review & editing.

MDM – conceptualization, data curation, formal analysis, methodology, project administration, software, validation, writing – original draft, writing – review & editing.

