## Supplementary Figures for "Ly6d expression delineates two putative postnatal thymus epithelial progenitor cells that are differentially affected by ageing"

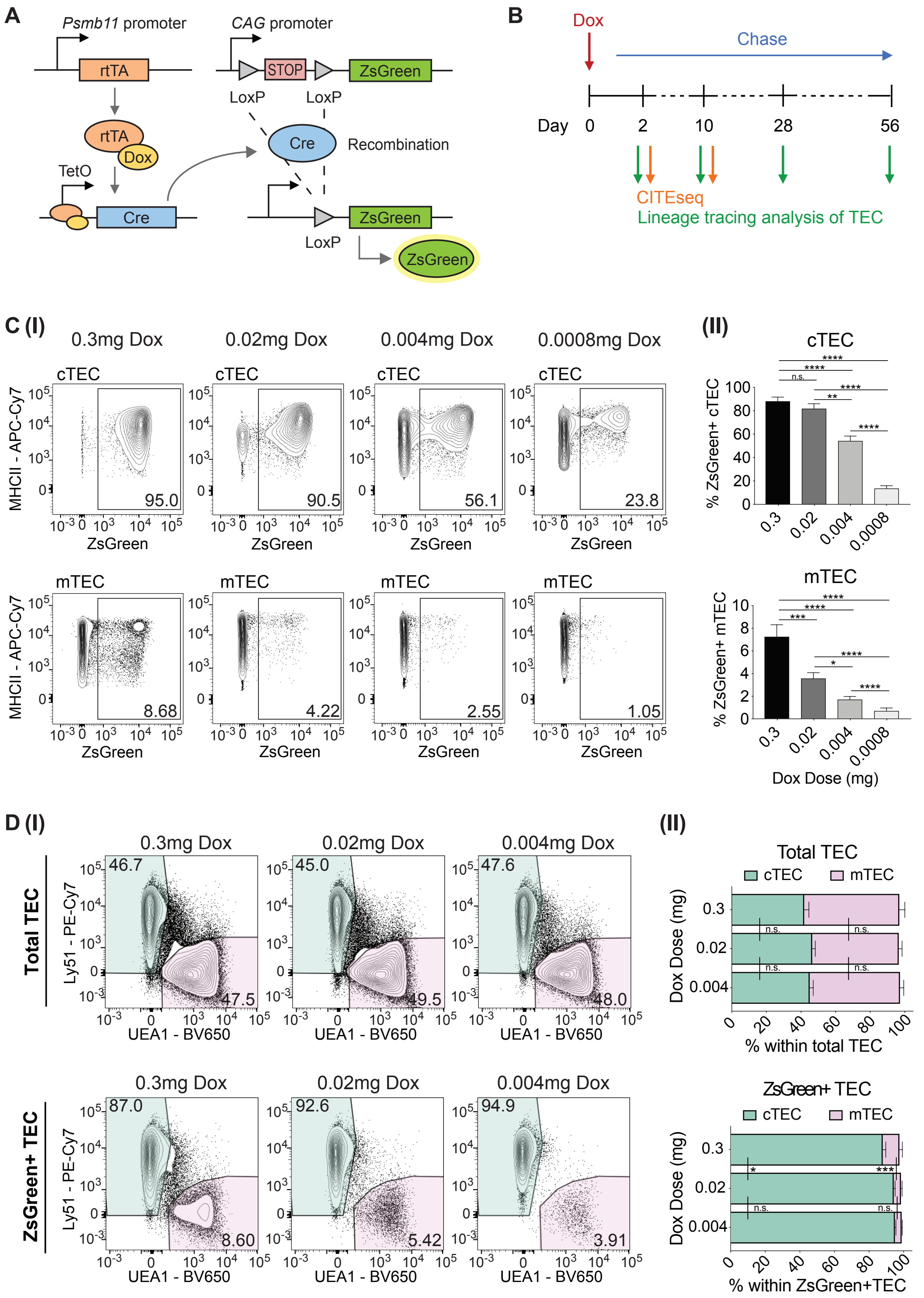


**Supplementary Figure 1.** (A) Cartoon depicting the TEC-specific, time-controlled and irreversible labelling of TEC from 3xTg mice, driven by the Psmb11 promoter. (B) Schematics of the treatment and data collection schedule. The time of doxicycline (Dox) i.p. injection of 7-day old pups is considered as Day 0 for the purpose of this report. (C) (I) Representative

FACS plots and (II) quantification of ZsGreen+ cTEC (top) and mTEC (bottom), in mice injected with decreasing doses of Dox 48h prior. n = 11 mice per group, except for the 0.0008mg dose which consisted of only n = 6 mice. Average proportion of labelled cTEC vs. mTEC: 87.2% ± 12.1% vs. 7.2% ± 3.6% for the 0.3 mg dose; 80.8% ± 14.1% vs. 3.5% ± 1.7% for the 0.02 mg dose; 53.5% ± 13.9% vs. 1.7% ± 1.0% for the 0.004 mg dose; and 13.2% ± 6.1% vs. 0.7% ± 0.7% for the 0.0008 mg dose) (D) (I) Representative FACS plots and (II) quantification of TEC compartment composition within total TEC (top), as a reference, and ZsGreen+ TEC (bottom) in mice injected with decreasing doses of Dox 48h prior. n = 11 mice per treatment group. Error bars represent the standard error of the mean. Statistical significance was determined by multiple unpaired Welsch t tests, * $p < 0.05$, ** $p < 0.01$, *** $p < 0.001$, **** $p < 0.0001$.

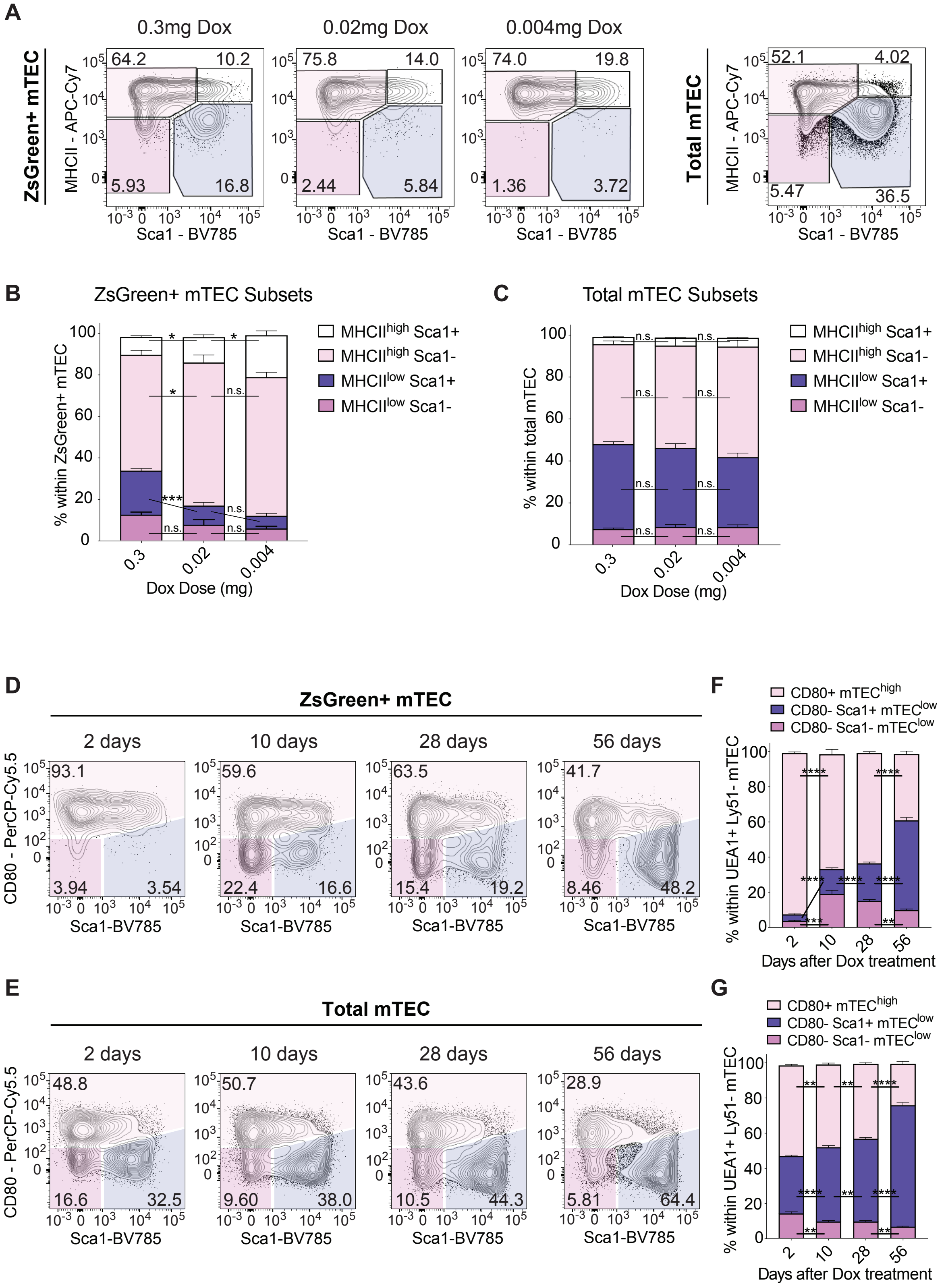


**Supplementary Figure 2.** (A) Representative FACS plots comparing the surface expression of MHCII and Sca1 on ZsGreen-labelled mTEC in mice injected with decreasing doses of Dox 48h prior. The panel on the right depicts total mTEC as a reference. Quantification of the relative proportions of different mTEC subsets as defined by their surface expression of

MHCII and Sca1, within (B) ZsGreen-labelled mTEC and (C) total mTEC, 48h after Dox injection. n = 8 mice per treatment group. Error bars represent the standard error of the mean. Statistical significance was determined by multiple unpaired Welsch t tests, * $p < 0.05$, ** $p < 0.01$, *** $p < 0.001$, **** $p < 0.0001$. Representative FACS plots comparing the surface expression of CD80 and Sca1 on (D) ZsGreen-labelled mTEC and (E) total mTEC; at the time-points of 2, 10, 28 and 56 days after treatment with 0.004mg of Dox. Quantification of the relative proportions of different mTEC subsets as defined by their surface expression of CD80 and Sca1, within (F) ZsGreen-labelled mTEC and (G) total mTEC; at the time-points of 2, 10, 28 and 56 days after treatment with 0.004mg of Dox. n = 8 mice per time-point, except for the first group (2 days) which consisted only of 7. Error bars represent the standard error of the mean. Statistical significance was determined by multiple unpaired Welsch t tests, * $p < 0.05$, ** $p < 0.01$, *** $p < 0.001$, **** $p < 0.0001$.

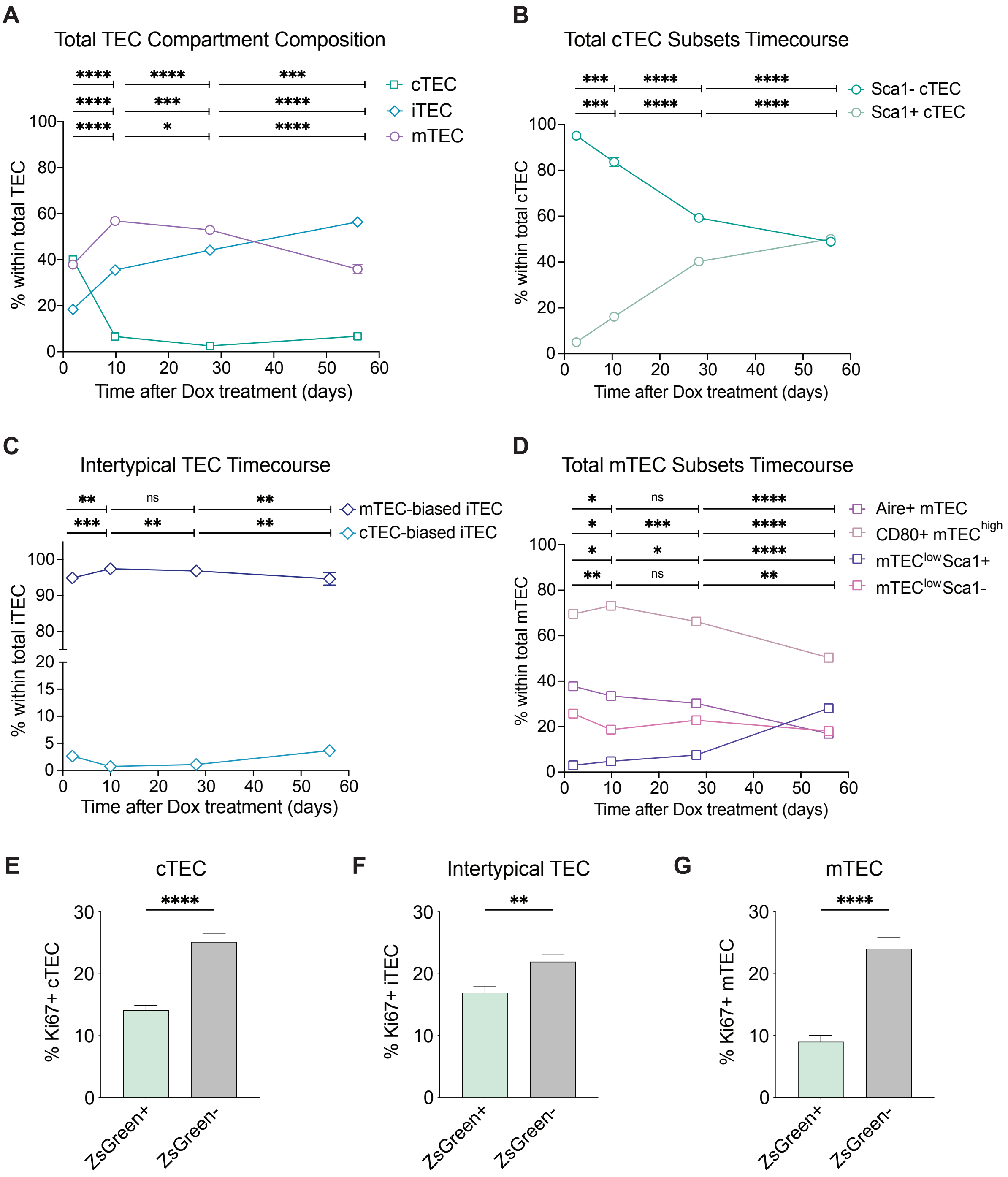


**Supplementary Figure 3.** Analysis of age-related changes in TEC compartment composition within (A) the whole TEC compartment, (B) cTEC subsets, (C) iTEC subsets and (D) mTEC subsets, at the time-points of 2, 10, 28 and 56 days following IP injection of 0.004 mg doxicycline at 7 days of age. n = 7 mice for the 2-day time-point, and n = 8 for the 10- 28- and 56-day time-points. Error bars represent the standard error of the mean. Statistical significance was determined by using multiple t tests to calculate differences in TEC subset composition from one time-point to the next. Correction for multiple comparisons was performed using the Holm-Sidak method. * $p<0.05$, ** $p<0.01$, *** $p<0.001$, and **** $p<0.0001$. Quantification of the proportion of Ki67+ cells within the ZsGreen positive and negative fractions of the (E) cTEC, (F) intertypical TEC and (G) mTEC compartments, 2 days after treatment with 0.004mg of Dox at 7 days of age. n = 7. Error bars represent the standard error of the mean. Statistical significance was determined by using unpaired t tests with Welsch's correction. * p<0.05, ** p<0.01, *** p<0.001, and **** p<0.0001.

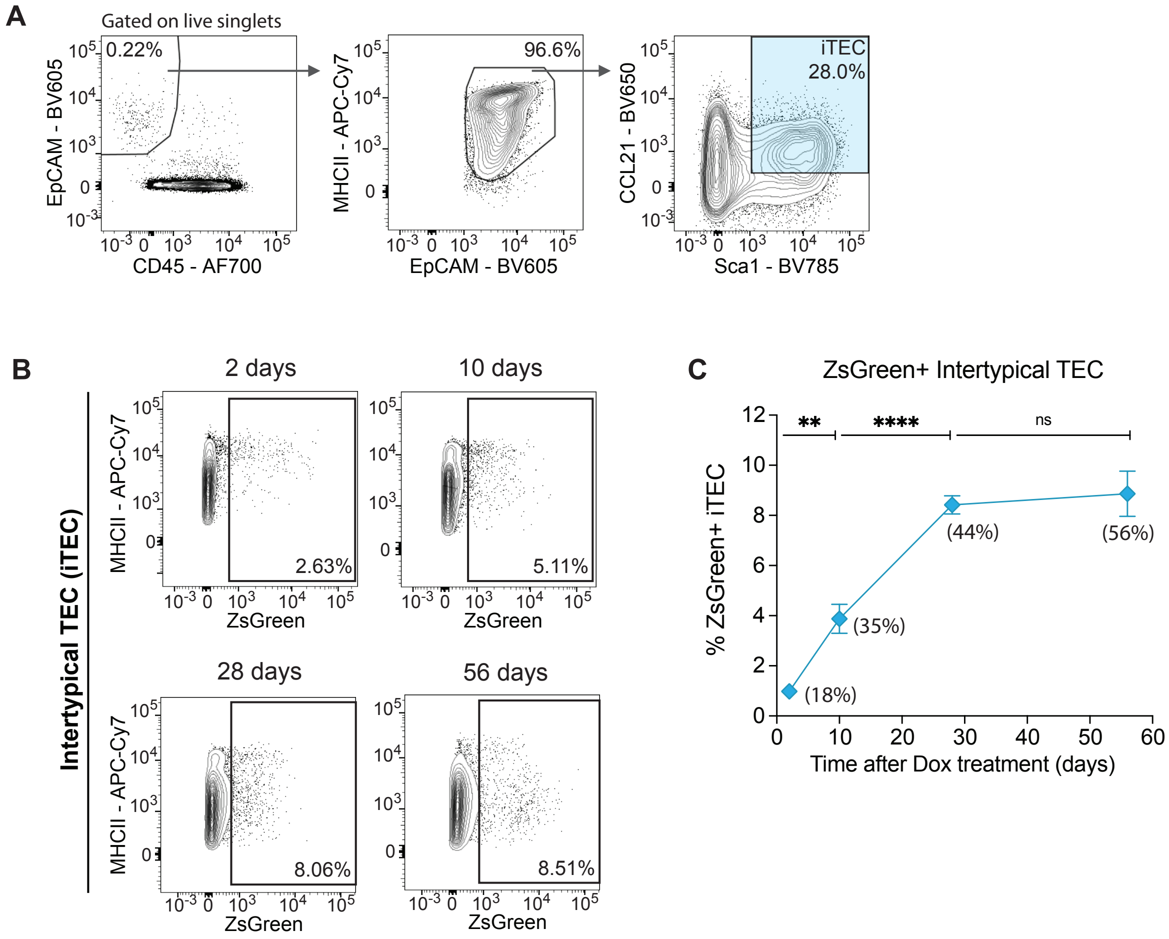


**Supplementary Figure 4.** (A) Representative FACS plots depicting the gating strategy followed for flow cytometric identification of intertypical TEC (iTEC, defined as EpCAM+, CD45-, MHCII+, Sca1+ and CCL21+). This example corresponds to a 17-day old mouse, 10 days post-treatment with a single dose of 0.004mg of Dox. (B) Representative FACS plots and (C) quantification of the proportion of ZsGreen+ intertypical TEC at the time-points of 2, 10, 28 and 56 days after injection with 0.004mg of Dox. The average proportion of total iTEC at each time-point is shown between brackets for reference. n = 7 mice for the 2-day time-point, and n = 8 for the 10- 28- and 56-day time-points. Error bars represent the standard error of the mean. Statistical significance was determined by multiple t tests to determine differences in TEC subset composition from one time-point to the next. Correction for multiple comparisons was performed using the Holm-Sidak method. * $p<0.05$, ** $p<0.01$, *** $p<0.001$, and **** $p<0.0001$.

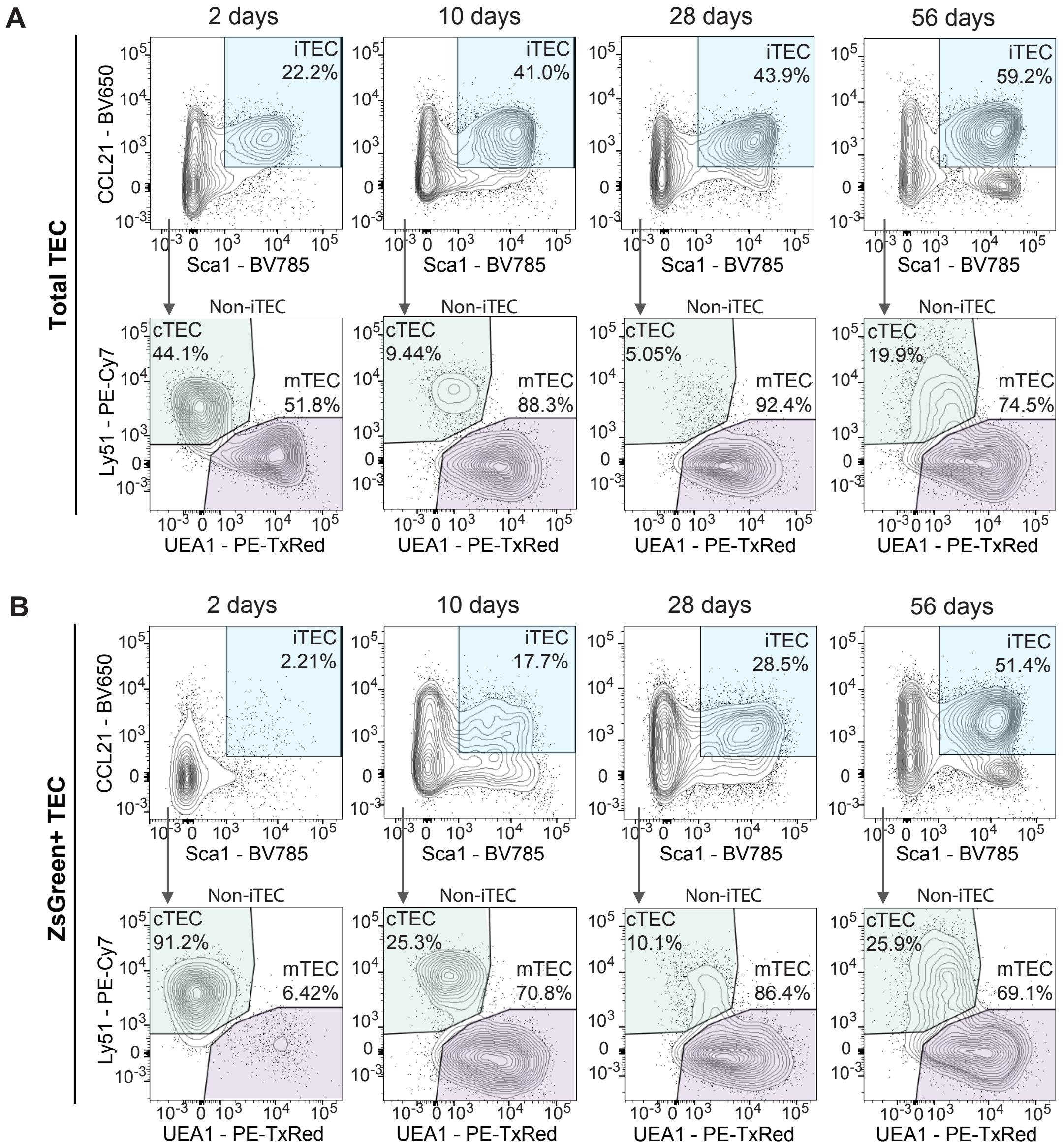


**Supplementary Figure 5.** Representative FACS plots of (A) total TEC subsets compared to (B) ZsGreen+ TEC, identifying intertypical TEC (iTEC, defined as Sca1+ and CCL21+), cTEC (defined as Ly51+ UEA1- out of non-iTEC) and mTEC (defined as Ly51- UEA1+ out of non-iTEC) at the time-points of 2, 10, 28 and 56 days after injection with 0.004mg of Dox.

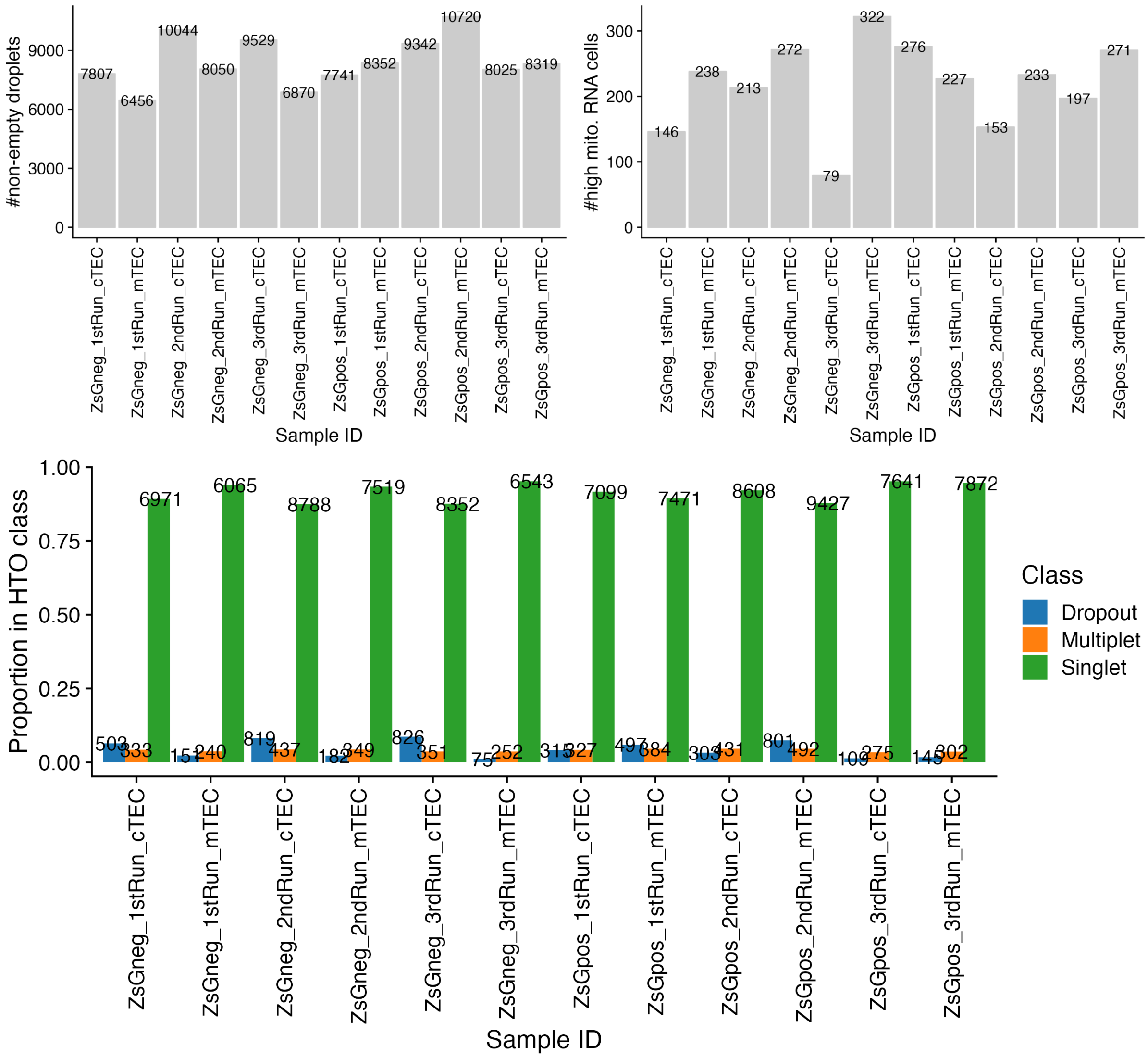


**Supplementary Figure 6.** Single cell RNA-sequencing quality control. (A) Cell calling non-empty droplets for each sample library. (b) Number of cells/droplets removed in each sample due to excessively high mitochondrial RNA content. (C) Multiplet and cell dropout detection using cell hashtags across all multiplexed sample libraries.

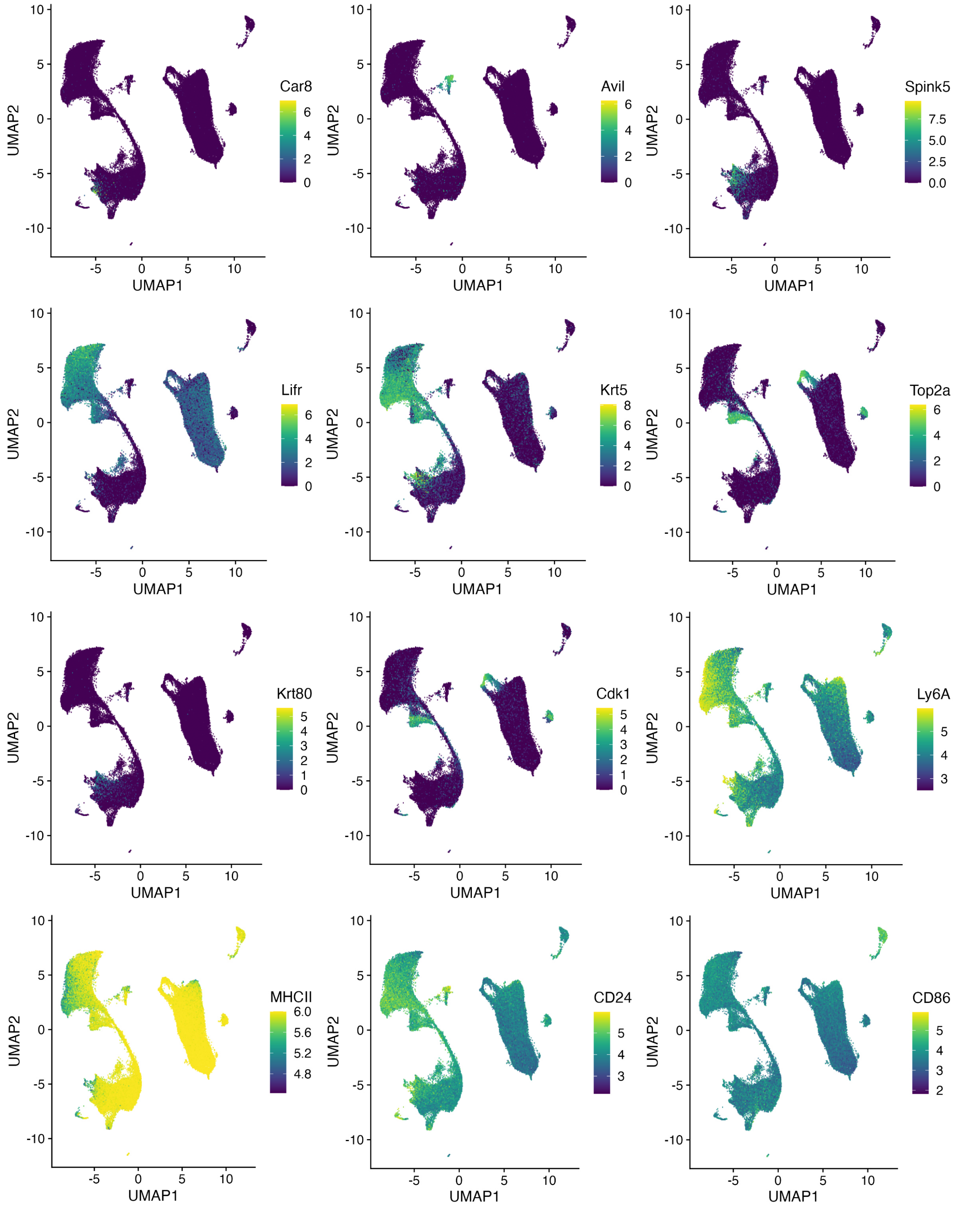


**Supplementary Figure 7.** Thymic cell type marker genes and surface marker expression (Ly6A, MHCII, CD24, CD86). Each panel shows a UMAP (as in Figure 2) coloured by expression level of the marker gene that illustrates specific cell types (*Car8, Avil, Spink5, Lifr, Krt5, Krt80*) and cell division state (*Top2a*, *Cdk1*).

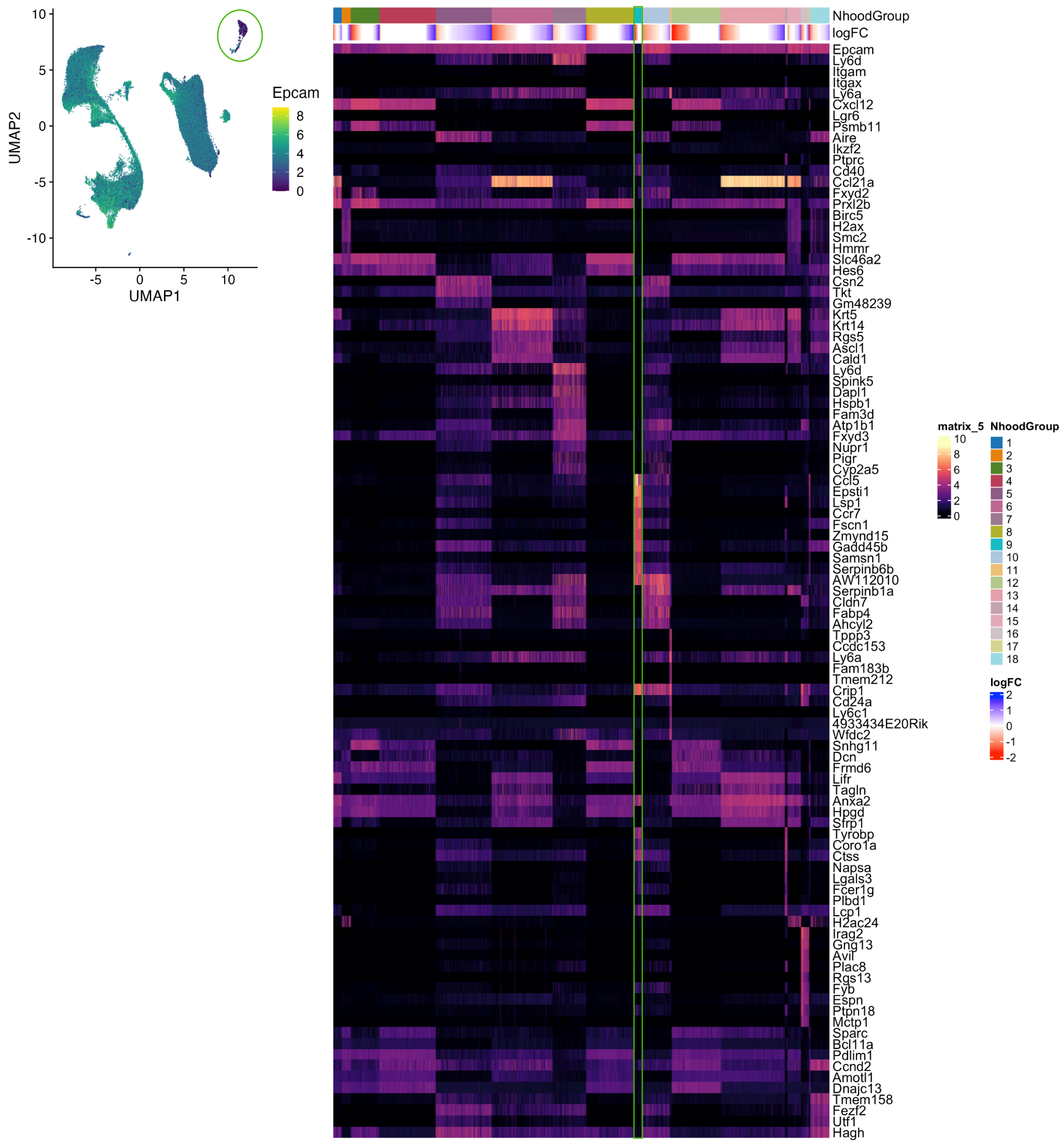


**Supplementary Figure 8.** Left panel highlights innate-like lymphoid cells (ILCs) that are sorted using surface EpCam+ staining but lack detectable *EpCam* mRNA. Right panel shows heatmap of marker genes across clusters. The column of ILCs are highlighted in green.

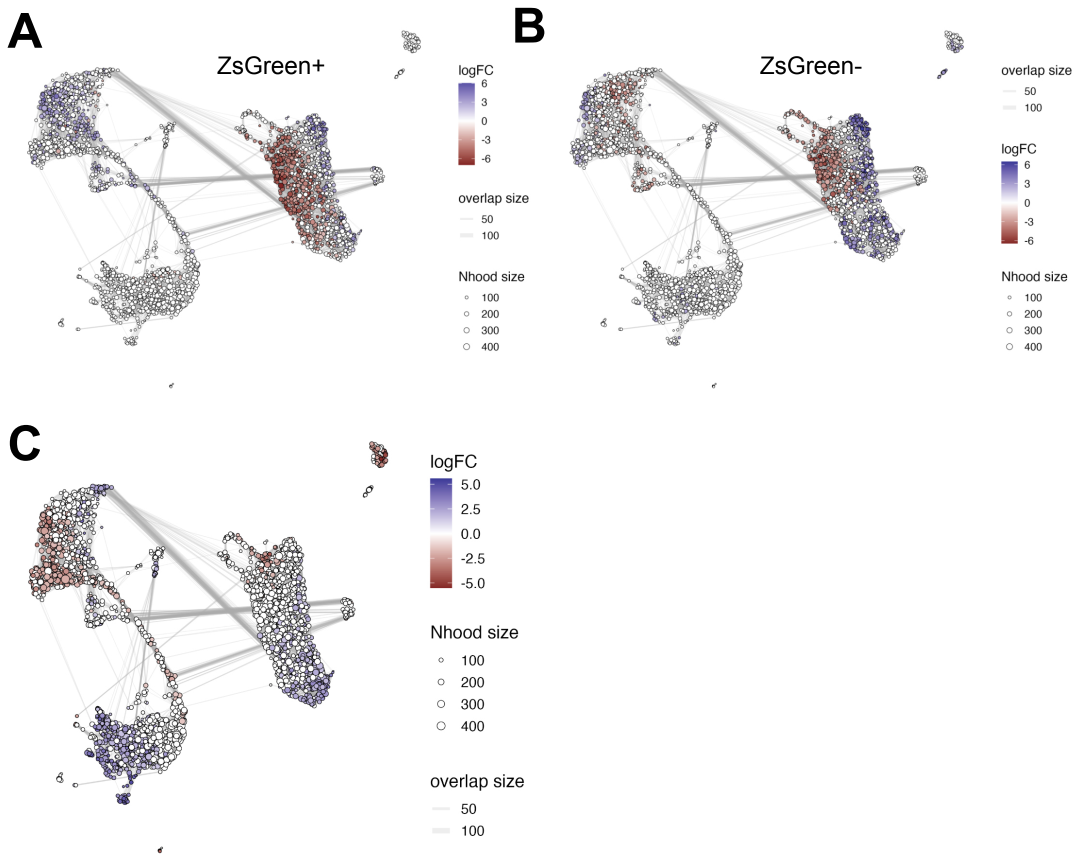


**Supplementary Figure 9.** (A-B) UMAP with differential abundance testing results overlaid showing the differential accumulation and depletion of TEC states from 2 to 10 days post dox treatment in ZsGreen+ (A) and ZsGreen- (B) sorted cells. Colour is log fold change between 2 and 10 days post-dox treatment. (C) UMAP with differential abundance testing results comparing ZsGreen+ and ZsGreen- sorting fractions, adjusted for post-dox treatment day. Points denote neighbourhood of cells and edges illustrate the number of cells shared between pairs of neighbourhoods.

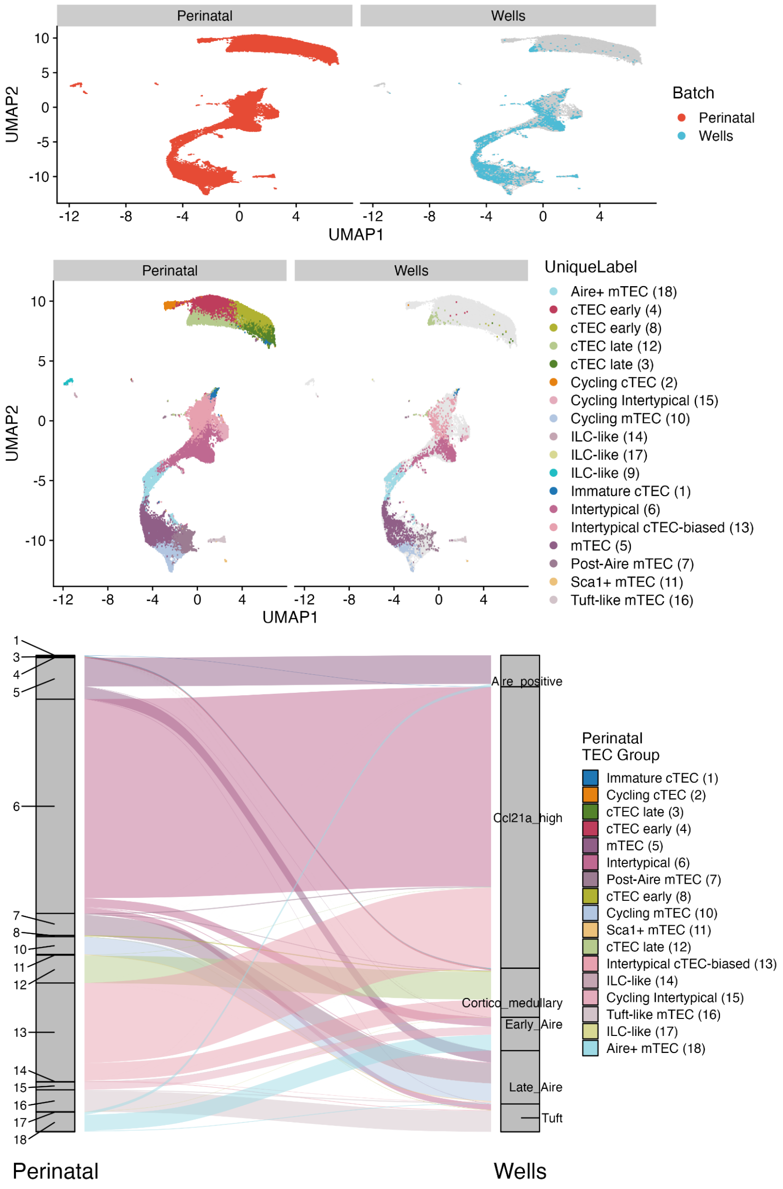


**Supplementary Figure 10.** UMAPs of cross-data set integration with Wells *et al.* scRNA-seq data coloured by data set (top panel) and cell type annotation (middle panel, as Figure 2E). Bottom panel shows an alluvium plot of cross-cell type label mapping from our data (left) to Wells *et al*. (right). Each line represents a single cell coloured by cell type (as in Figure 2E).

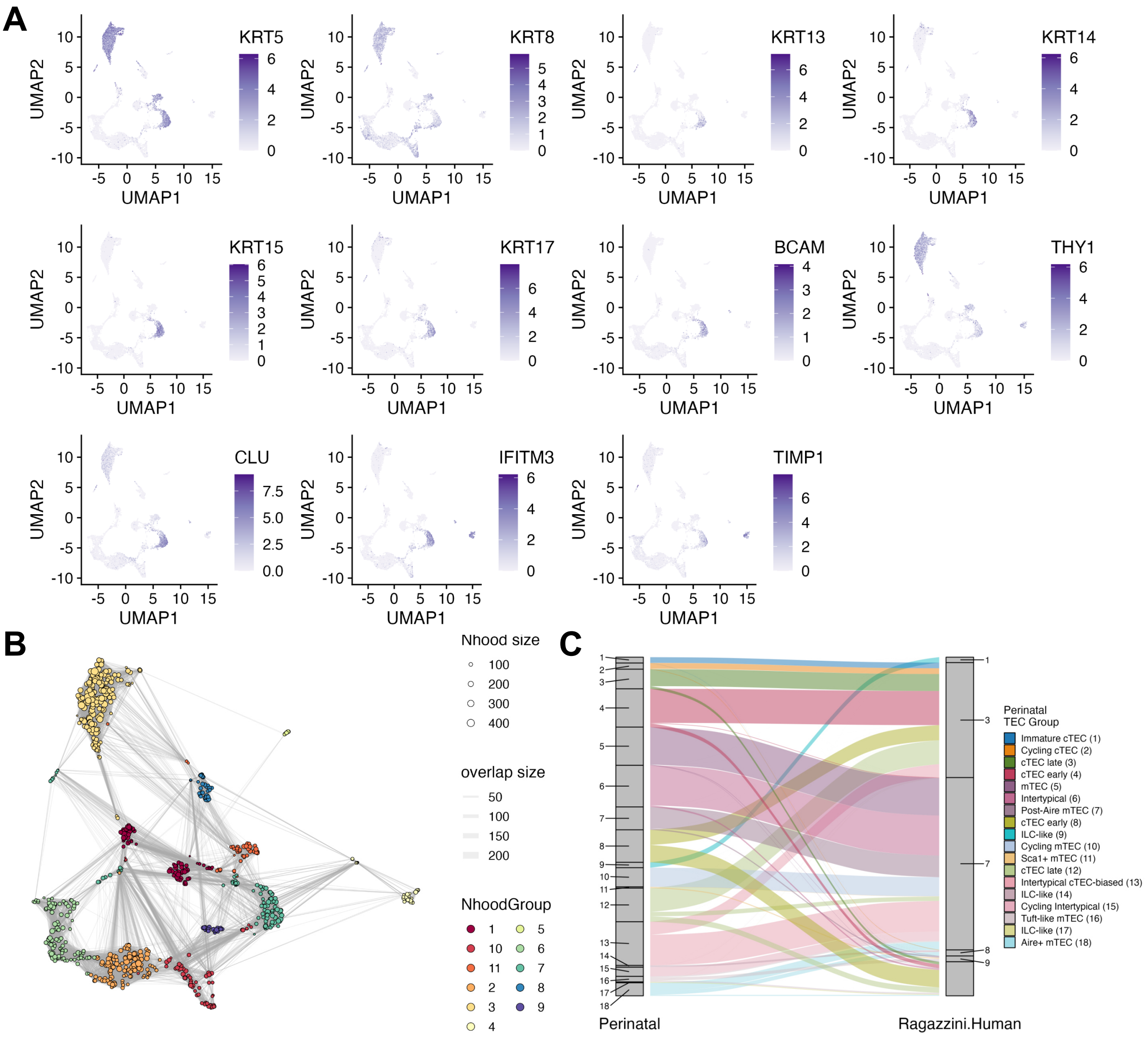


**Supplementary Figure 11.** (A) Gene expression of poly-keratin signature genes overlaid on a UMAP of data from Ragazzini et al. Each point is a single cell, and log normalized gene expression is shown by colour scale. (B) Neighbourhood UMAP of single-cell data from Ragazzini et al using UMAP co-ordinates as in (A). Points are neighbourhoods of cells, coloured by Louvain clustering on the neighbourhood adjacency matrix. Lines represent the number of cells shared by between neighbourhoods and point size corresponds to the number of cells. (C) Alluvium plot showing the cross-mapping of neighbourhoods between mouse perinatal TEC and human TEC from Ragazzini et al, coloured by the neighbourhood cluster as in Figure 2E.

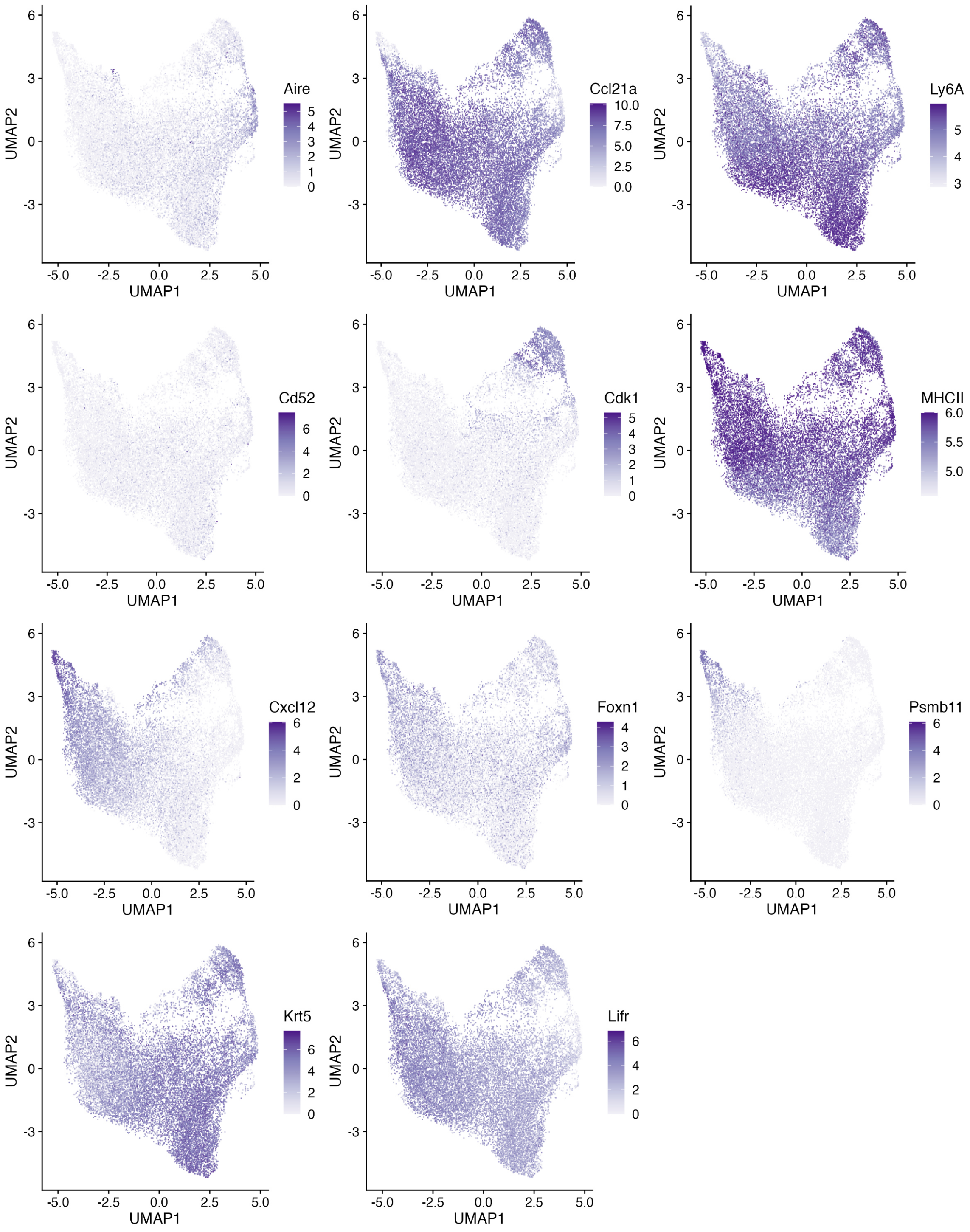


**Supplementary Figure 12.** UMAP of Intertypical subset of perinatal mouse TEC. Each point is a single cell with marker genes (*Aire, Ccl21a, Cd52, Cdk1, Cxc12, Foxn1, Psmb11, Krt5 and Lifr*) and CITE-protein (MHCII, Ly6A) measurements overlaid in colour.

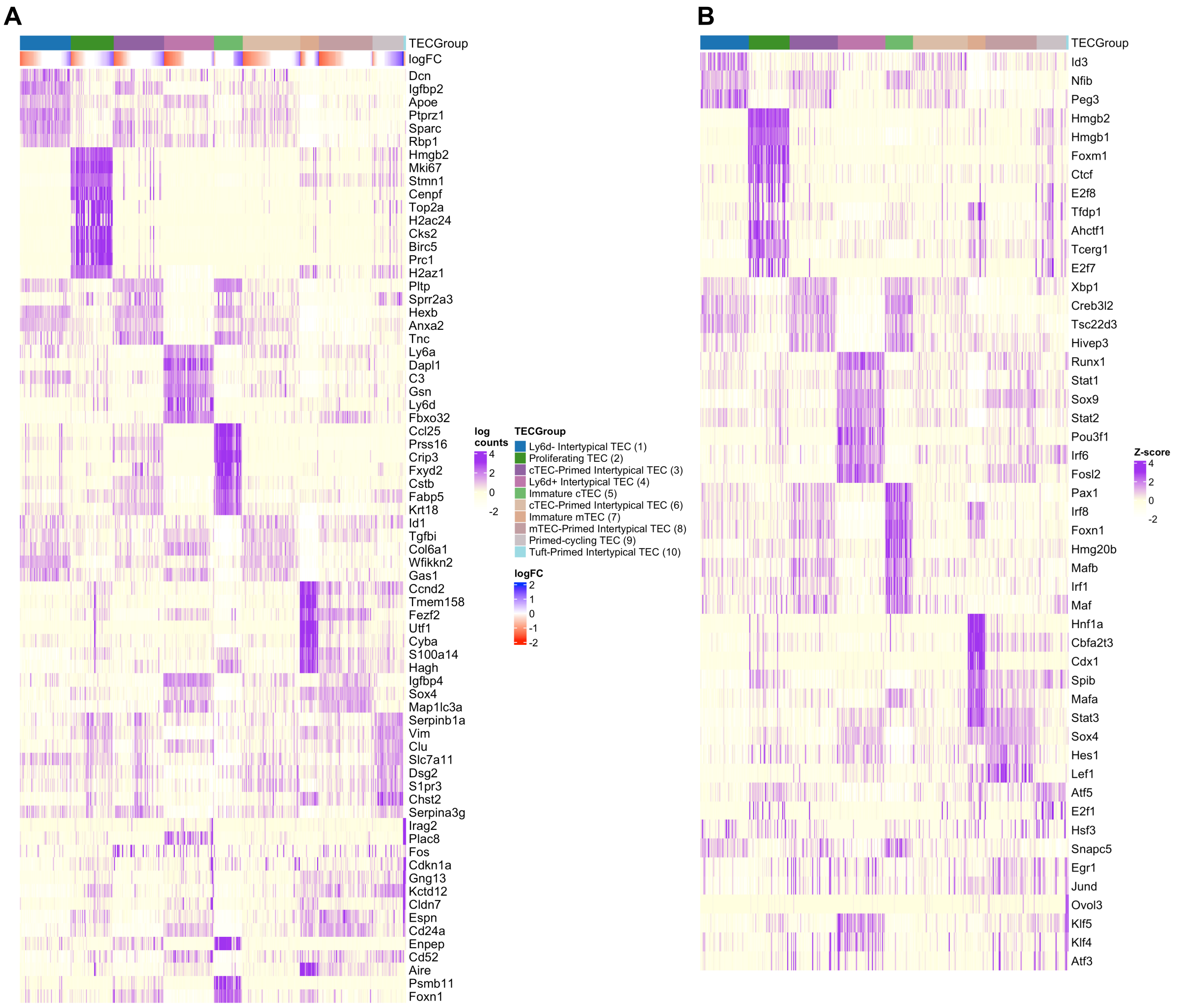


**Supplementary Figure 13.** Heatmaps of intertypical TEC neighbourhood subpopulations marker genes (A) and transcription factors (B). Each column represents a neighbourhood and each row a marker gene or marker transcription factor. Columns are ordered by neighbourhood cluster (NhoodGroup). Log fold-change denotes the difference between days 2 and 10, adjusting for ZsGreen sorting fraction. Gene expression values are row-standardised (Z-scores) to highlight the enrichment of marker genes in each neighbourhood cluster. *Foxn1*, *Aire* and *Cd52* were included as known markers of mTEC and cTEC lineages.

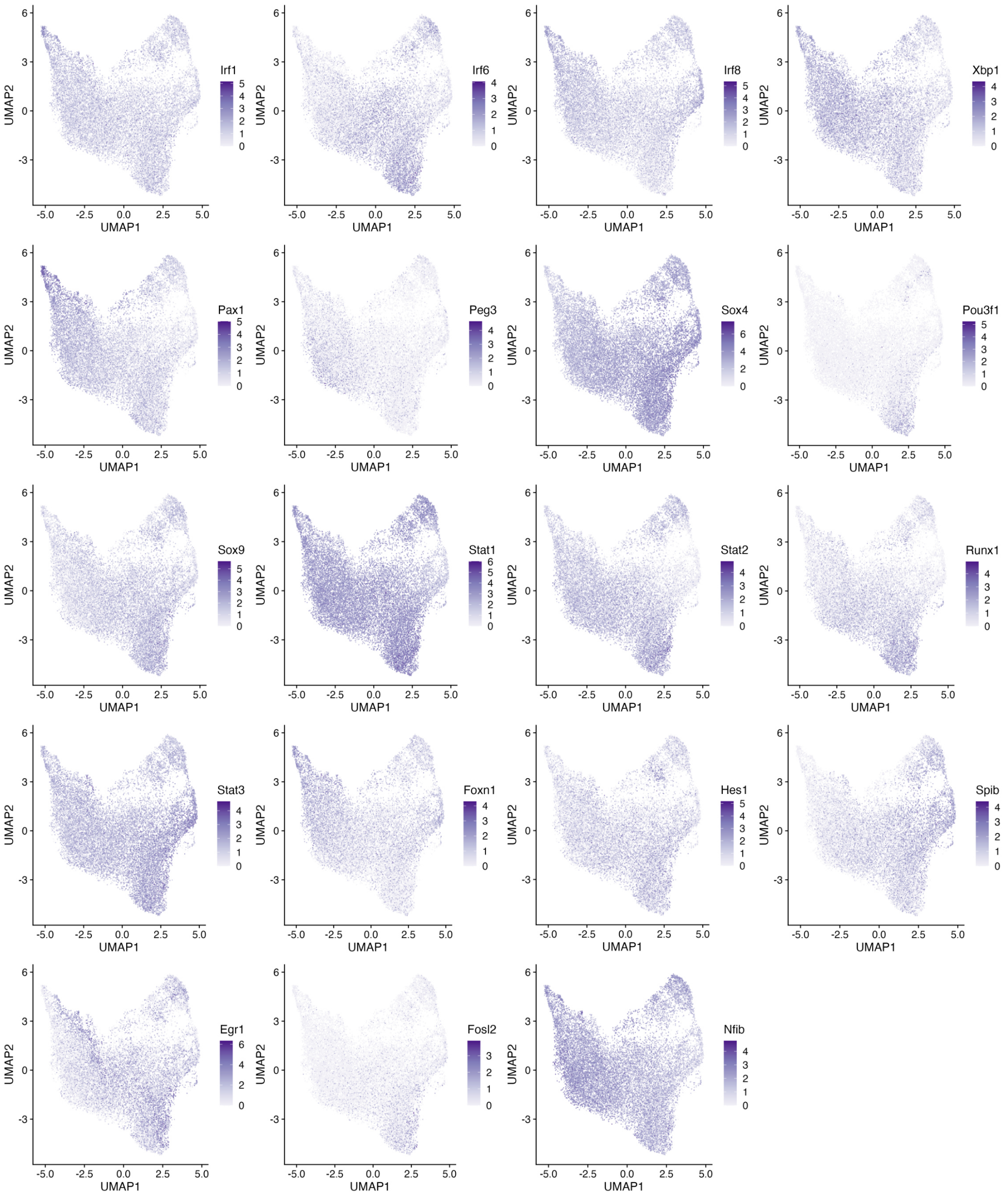


**Supplementary Figure 14.** Intertypical TEC sub-population characteristic transcription factor gene expression overlaid on the Intertypical TEC subset UMAP. Each point is a gene and each panel is a gene with annotated DNA activity according to Gene Ontology.

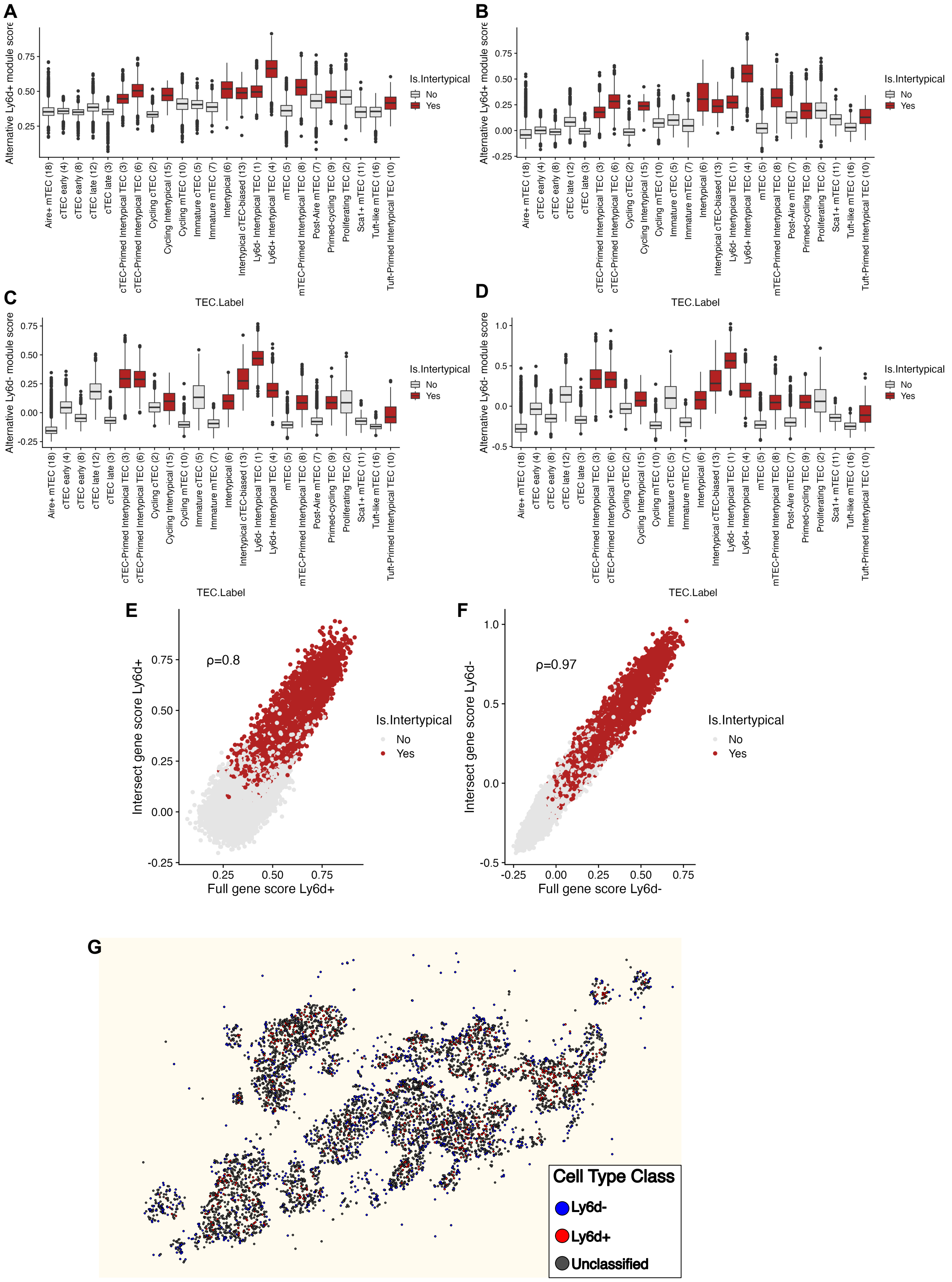


**Supplementary Figure 15.** Boxplots of modules scores (y-axis) of (A) Ly6d+ and (B) Ly6d- intertypical TEC gene sets before and after intersection with Xenium measured genes (C, D) across all TEC subtypes (x-axis). Boxes are coloured red for intertypical TEC subtypes and grey for all others. Boxes show the median and IQR with whiskers extending 1.5x the IQR and outliers beyond this threshold shown as individual points. (E) and (F) show concordance between module scores using the full and Xenium-measured genes sets for Ly6d+ (E) and Ly6d- (F). Each point is a single cell coloured by whether they are intertypical TEC (red) or not (grey). (G) Xenium spatial transcriptomics data showing intertypical TEC classification using module scores as either Ly6d+ (red), Ly6d- (blue) or unclassified (grey). Only cells classified as intertypical TEC are shown.

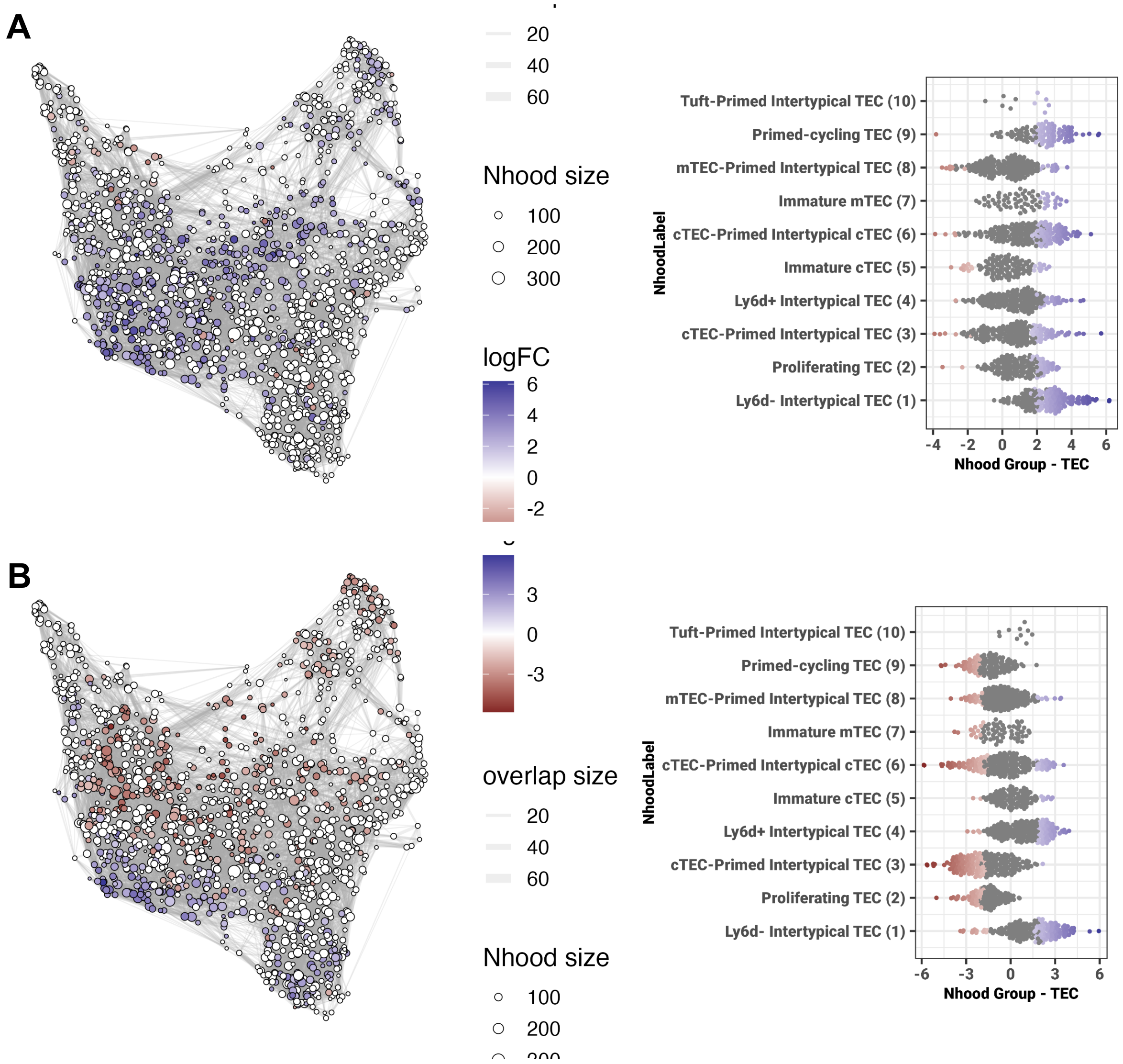


**Supplementary Figure 16.** Differential abundance testing results based on day post dox treatment in ZsGreen+ (A) and ZsGreen- (B) samples. Left shows neighbourhood UMAP coloured by log fold change from DA testing and right shows Beeswarm plots of log fold change (x-axis) versus cell type annotations (y-axis). UMAP points are neighbourhoods, with size denoting the number of cells and lines between neighbourhoods are proportional to the number of shared cells between pairs of neighbourhoods.

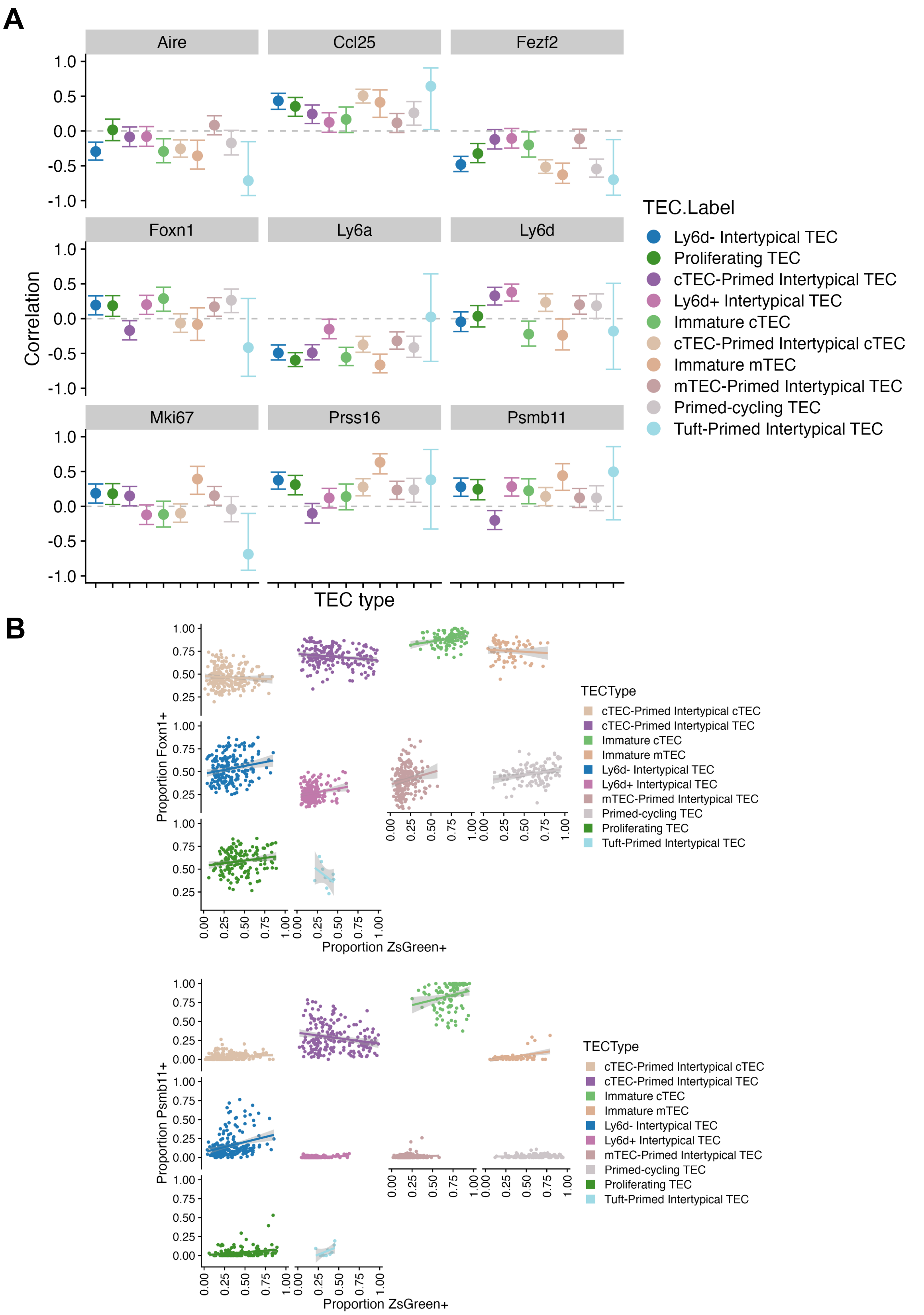


**Supplementary Figure 17.** (A) Correlation of Foxn1-regulated (*Psmb11*, *Prss16*, *Ccl25*) and TEC cell state marker gene expression (*Aire*, *Fezf2*, *Ly6a*, *Ly6d*, *Mki67*) vs. proportion of ZsGreen+ cells over neighbourhoods for intertypical TEC subsets (x-axis). Points show the average correlation with 95% confidence intervals. The horizontal grey dotted line shows a Pearson correlation of zero. (B) Scatter plots of neighbourhood proportions of Foxn1+ (top) and Psmb11+ (bottom) neighbourhoods vs. proportion of ZsGreen+ cells in each. Points are coloured by cell type annotation as in Figure 3A.

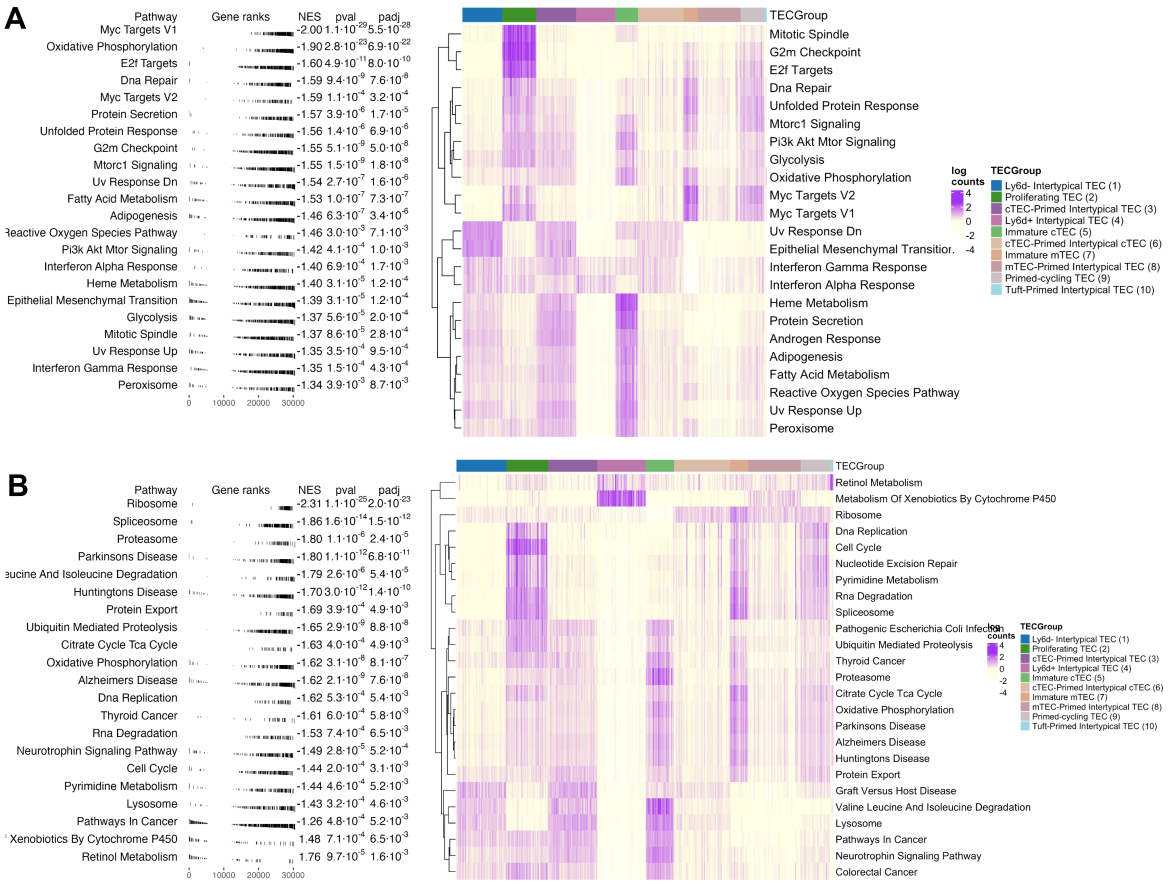

**Supplementary Figure 18.** Gene set enrichment analysis results of (A) Ly6d+ and (B) Ly6d- intertypical TEC marker genes. Left hand panel shows GSEA results for top enriched pathways and left heatmap shows average expression of genes in each gene set for each neighbourhood coloured by intertypical TEC annotation (columns).

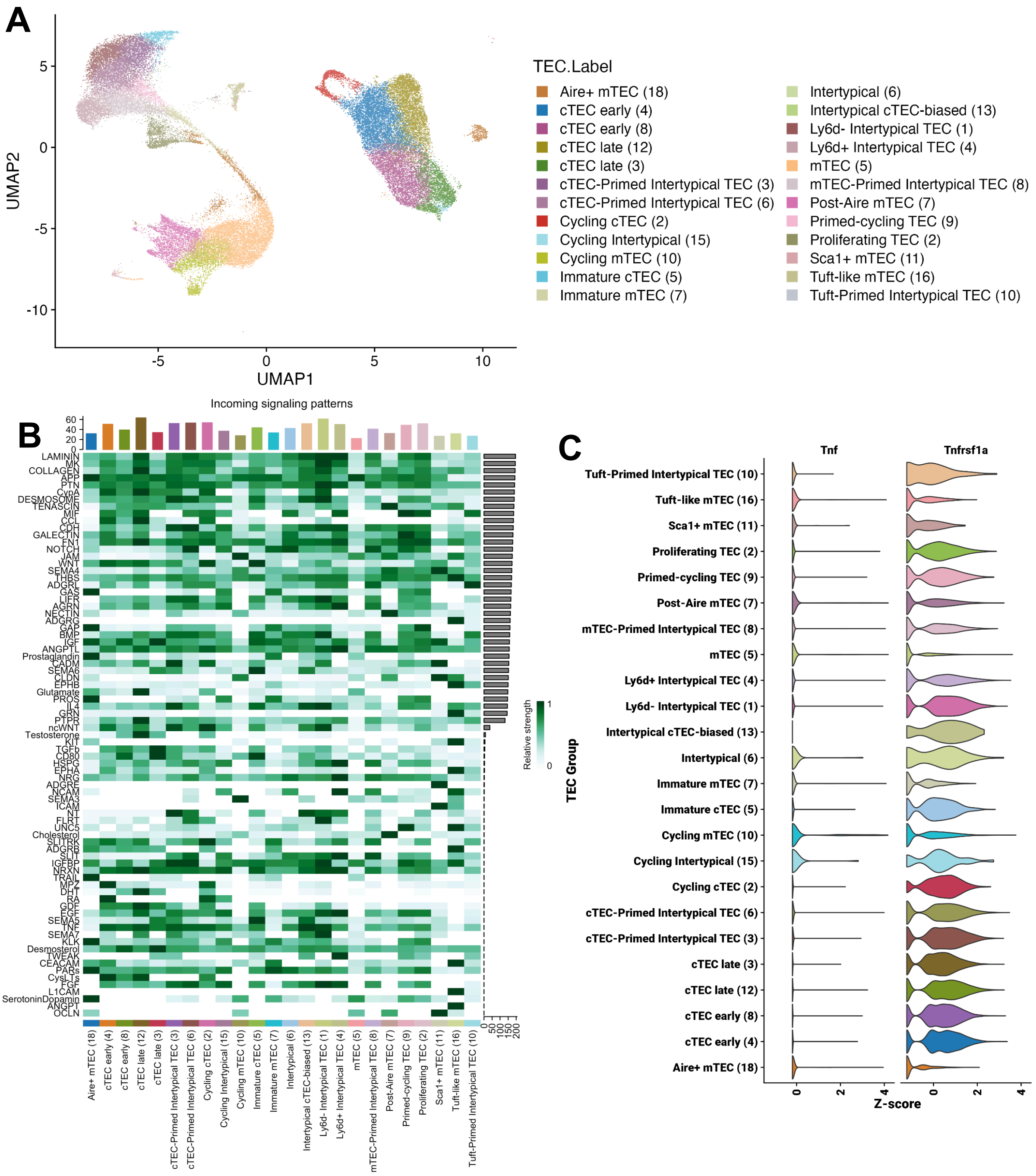


**Supplementary Figure 19.** (A) UMAP of combined TEC subtype annotations. (B) Inferred cell-cell signalling pathways active across all TEC subtypes in (A). Rows represent signalling pathways and columns are TEC subtypes. Heatmap cells are coloured based on the inferred strength of signalling by CellChat. (C) Violin plot of Tnf and Tnfrsf1a standardised (Z-score) single-cell expression across TEC subtypes (n=12 samples). Violin colours as in (A).

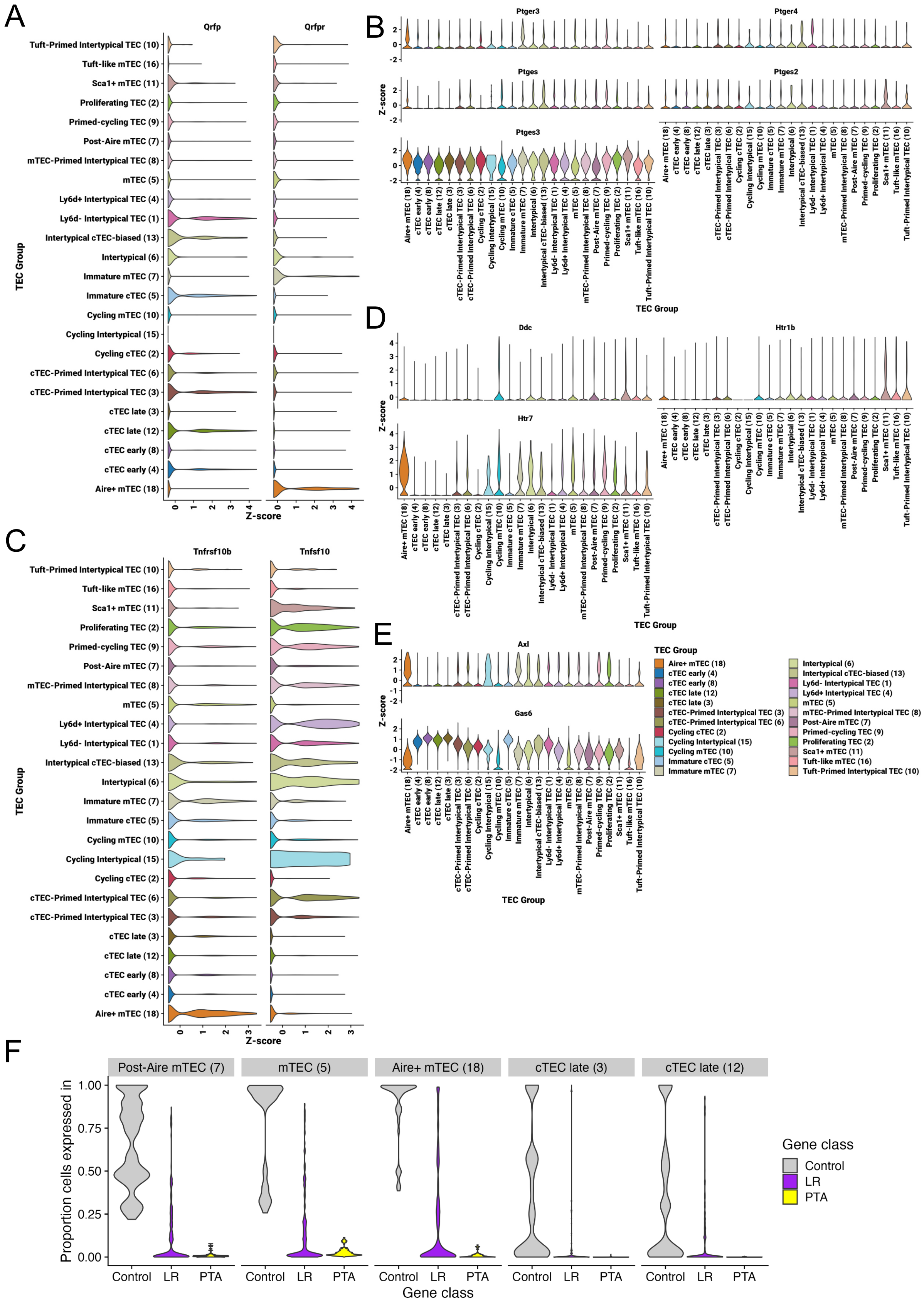


**Supplementary Figure 20**. (A-E) Violin plots of single-cell gene expression for the top 5 ligand receptor groups associated with incoming module 4 (Figure 4A-B). Each plot shows the Z-score across TEC subtypes. Colours according to panel E. (A) QRFP, (B) Prostaglandin, (C)TRAIL, (D) Dopamine/Serotonin, (E) GAS. (F) Violin plot of proportion of cells expressing each class of gene: control genes, ligand-receptor genes, or peripheral tissue antigen (PTA) genes.

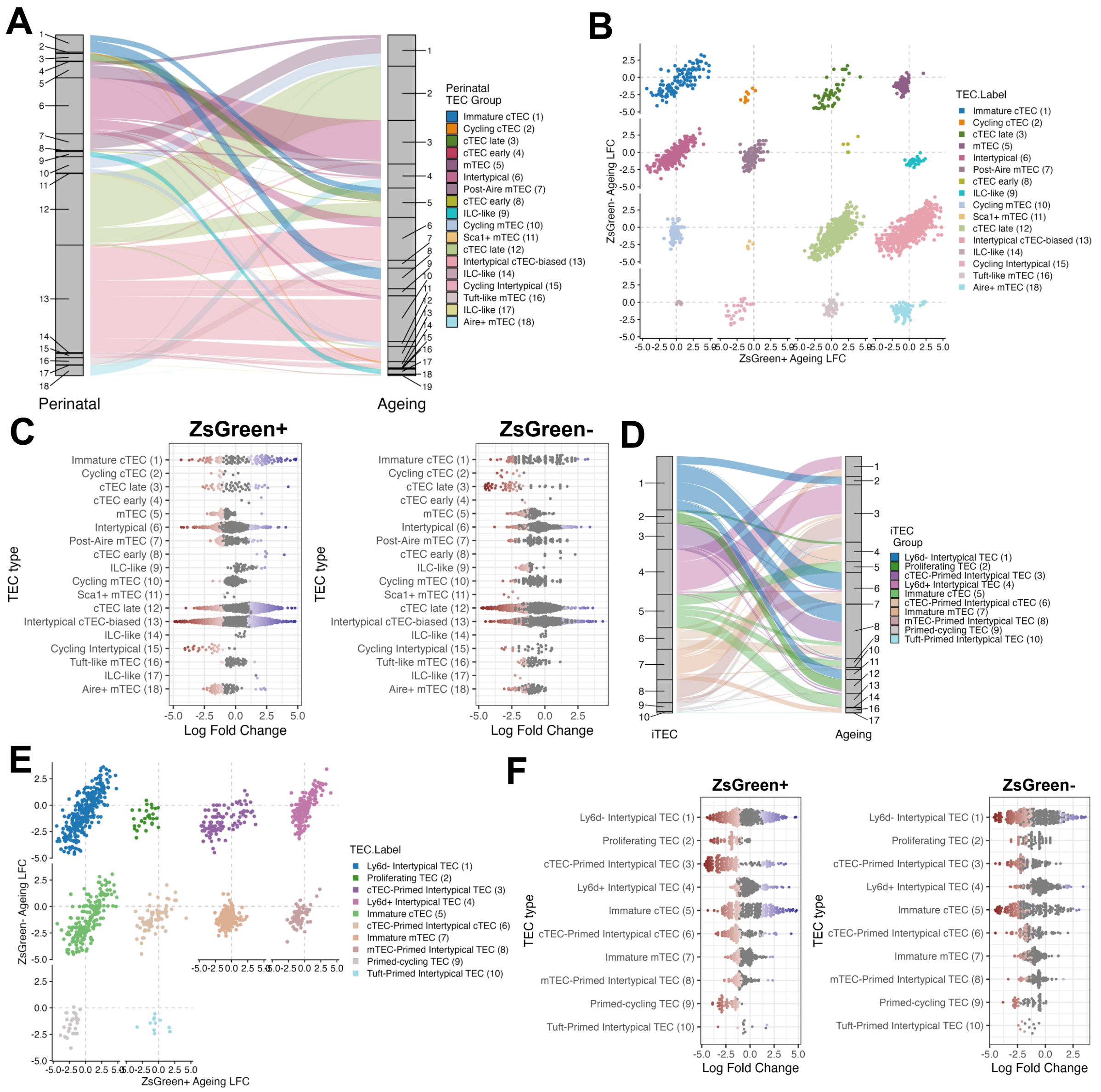


**Supplementary Figure 21.** (A) Cross-mapping neighbourhoods between perinatal and ageing data sets using maximal neighbourhood transcriptional correlation. Each line represents a neighbourhood, coloured by their annotation in the Perinatal data set. (B) Comparison of ageing DA testing log fold change results in the ZsGreen- (y-axis) and ZsGreen+ (x-axis) fractions. (C) Beeswarm plots of neighbourhood DA testing separately in ZsGreen+ (left) and ZsGreen- (right) sort fractions. DA testing compares neighbourhood abundance across ageing from 1 to 16 weeks old. Points are coloured by log fold change (x-axis) and annotated by Perinatal TEC subpopulation (y-axis), as in (A). (D) Cross-mapping neighbourhoods between perinatal intertypical TEC and ageing data sets using maximal neighbourhood transcriptional correlation. Each line represents a neighbourhood, coloured by their annotation in the Perinatal iTEC data set as in Figure 3A. (E) Comparison of ageing DA testing log fold change results in the ZsGreen- (y-axis) and ZsGreen+ (x-axis) fractions. (F) Beeswarm plots of neighbourhood DA testing separately in ZsGreen+ (left) and ZsGreen- (right) sort fractions. DA testing compares neighbourhood abundance across ageing from 1 to 16 weeks old. Points are coloured by log fold change (x-axis) and annotated by Perinatal intertypical TEC subpopulation (y-axis), as in (D).

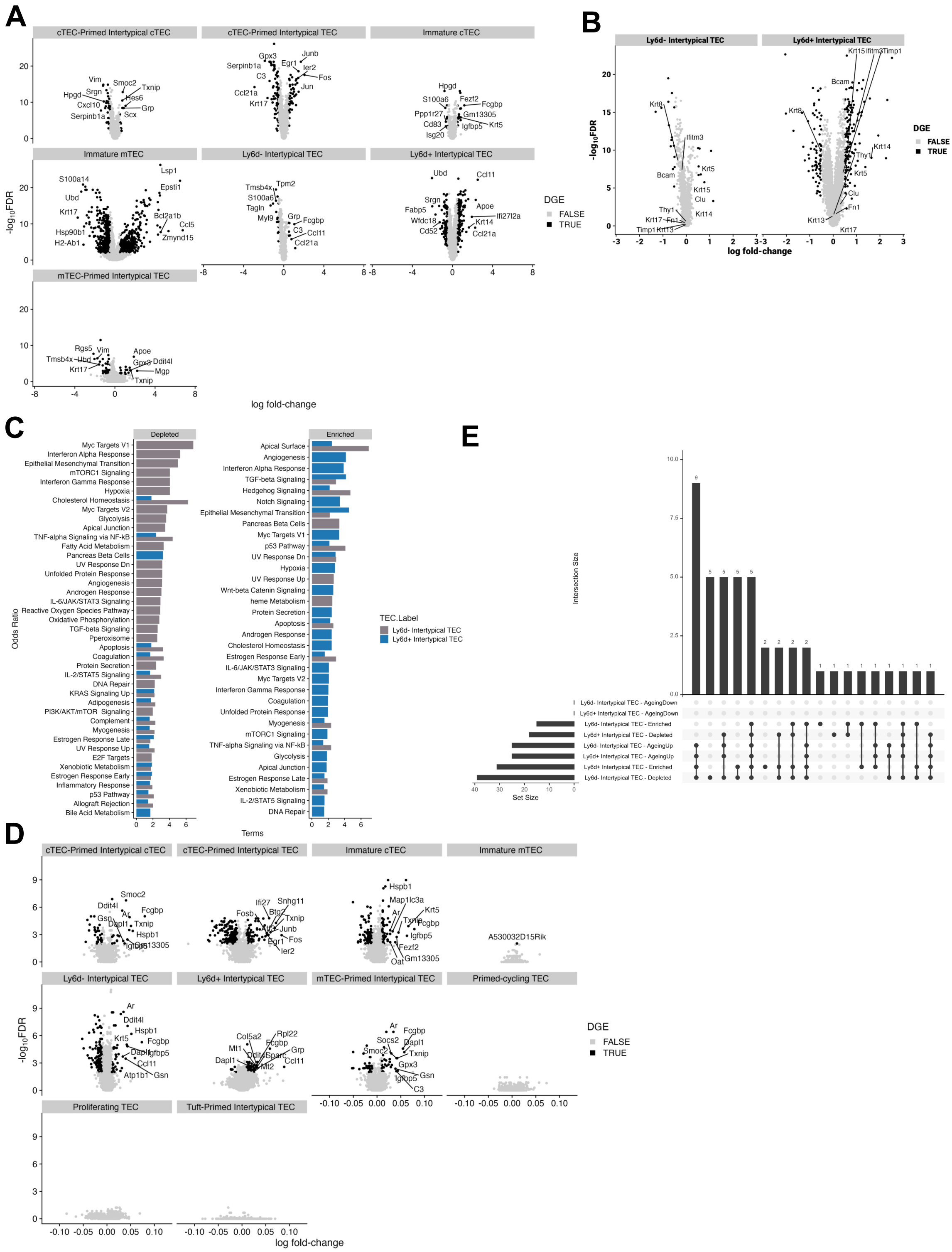

**Supplementary Figure 22.** (A) Volcano plots of differential gene expression changes between ageing enriched and depleted neighbourhoods within each TEC subpopulation. Log fold change is shown on the x-axis and -log10 FDR on the y-axis. Points are coloured black if they have FDR <1% and |log fold change| > 0.5. The top 10 DE genes for each TEC subpopulation are labelled. (B) As in (A) except genes are labelled according to marker genes for poly-keratin cells as in Ragazzini et al. (C) Gene set enrichment analysis results of MSigDB Hallmark pathways (y-axis) for genes up-regulated in ageing depleted (left) and enriched (right) neighbourhoods. X-axis shows log odds ratio of enrichment. (D) Volcano plots of ageing DEGs in each intertypical TEC sub-population; axes as in (A). Points are coloured black with |log fold change| > 0.1 and FDR < 1%. Labelled genes are the top 10 DGEs for each TEC sub-populaiton. (E) UpSet plot illustrating the small overlap between enriched pathways between DA and ageing differential expressed genes. Horizontal bars denote the number of pathways in each DE gene set and vertical bars denote the size of the intersection between each set of enriched pathways. Points and lines correspond to the set intersections.

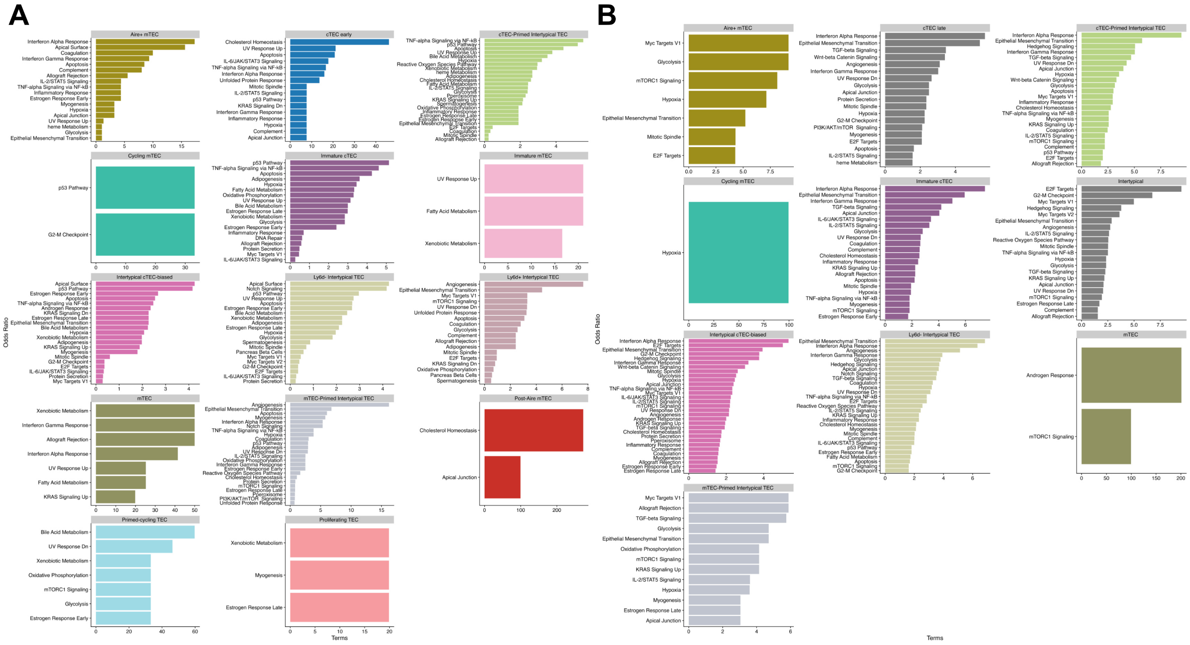


**Supplementary Figure 23.** Each panel shows the enriched MSigDB Hallmark gene sets that are (A) enriched or (B) depleted across ageing with a 10% FDR threshold.

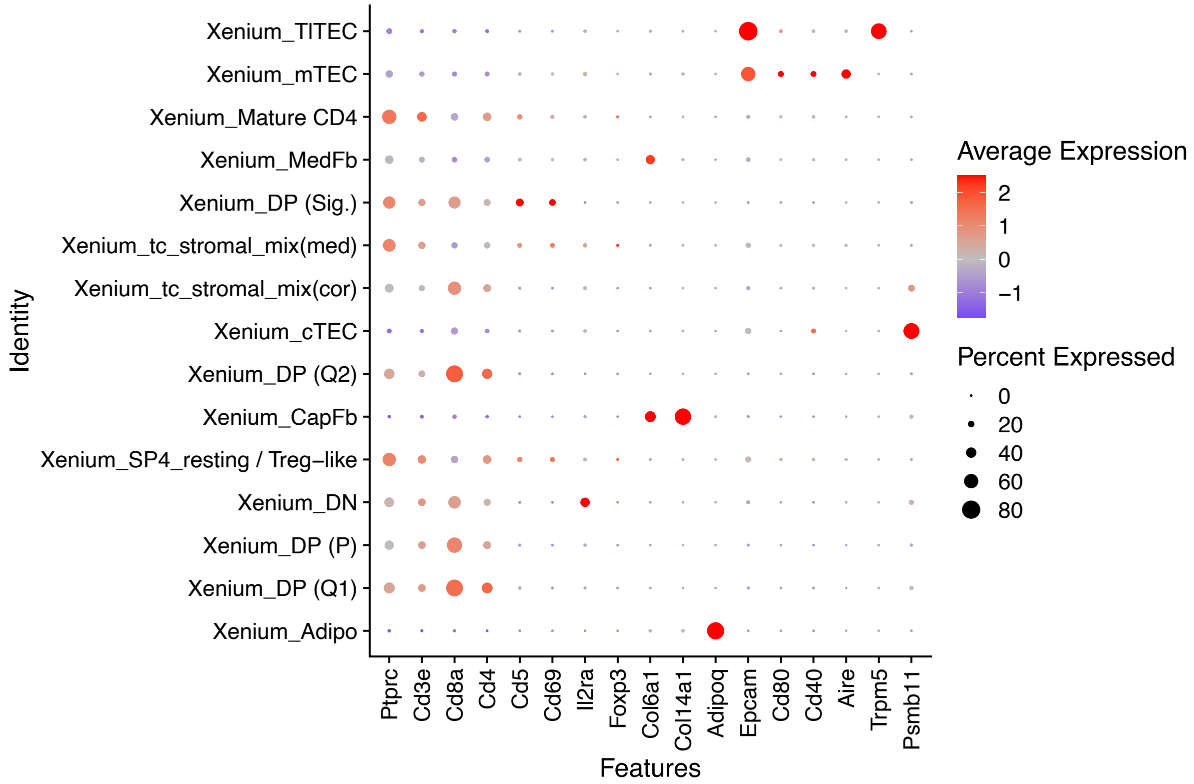


**Supplementary Figure 24.** Cell type annotation of Xenium data for histological compartment definition. Normalized expression profiles of selected genes across distinct cell types identified in the Xenium spatial transcriptomics dataset.
