## Supplementary Tables for "Ly6d expression delineates two putative postnatal thymus epithelial progenitor cells that are differentially affected by ageing"

***Supplementary Table 1:*** *list of antibodies/reagents used for flow cytometric phenotypic analysis and FACS cell sorting. Antibodies were diluted in FACS buffer (2% FCS in PBS), unless otherwise specified.*

| Marker | Clone | Conjugate | Source (Catalog #) | Usage |
| --- | --- | --- | --- | --- |
| AIRE | 5H12 | eFluor660 | eBioscience (50-5934-82) | 1:1000  BD Cytofix/Cytoperm^TM^ |
| CCL21 | Rabbit  polyclonal | Biotin | LSBio (LS-C104634) | 1:100  eBioscience^TM^ FoxP3 Staining buffer set |
| Streptavidin | N/A | BV650 | Biolegend (405232) | 1:500  eBioscience^TM^ FoxP3 Staining buffer set |
| CD45 | 30-F11 | PE-TexasRed | Invitrogen (MCD4517) | 1:500 |
| CD45 | 30-F11 | Alexa Fluor 700 | BioLegend (103128) | 1:400 |
| CD80 | 16-10A1 | PerCP-Cy5.5 | BioLegend (104722) | 1:500 |
| CD86 | GL-1 | BV650 | BioLegend (105036) | 1:500 |
| CD104 | 346-11A | PE | BioLegend (123609) | 1:500 |
| Dclk1 | Rabbit polyclonal | Unconjugated | Abcam (ab31704I) | 1:500  eBioscience^TM^ FoxP3 Staining buffer set |
| Anti-rabbit (H&L) | Goat polyclonal | Alexa Fluor 647 | Thermofisher (A-21244) | 1:1000  eBioscience^TM^ FoxP3 Staining buffer set |
| EpCAM (CD326) | G8.8 | BV421 | BioLegend (118225) | 1:500 |
| EpCAM (CD326) | G8.8 | PerCP-Cy5.5 | BioLegend (118220) | 1:500 |
| Ki67 | SolA15 | Alexa Fluor 700 | Invitrogen (56-5698-82) | 1:200  BD Cytofix/Cytoperm^TM^ |
| Ly51 | 6C3 | PE-Cy7 | BioLegend | 1:500 |
| MHC-II (I-A / I-E) | M5/114.15.2 | APC-Cy7 | BioLegend (107628) | 1:1000 |
| Sca1 (Ly-6A/E) | D7 | BV750 | Biolegend (108139) | 1:1000 |
| UEA1 | Lectin | Cy5 | Vector Laboratories (self-conjugated) | 1:500 |
| UEA1 | Lectin | DyLight594 | Vector Laboratories (DL-1067) | 1:500 |
| UEA1 | Lectin | Rhodamine | Vector Laboratories (RL-1062) | 1:500 |
| Zombie Violet^TM^ Fixable Viability Dye | | | Biolegend (423113) | 1:1000  PBS |

***Supplementary Table 2:*** *list of TotalSeq-A oligonucleotide-conjugated antibodies (sourced from BioLegend) used for Droplet sequencing.*

| Marker | Clone | Conjugate | HTO/ADT ID | Barcode Sequence | Usage |
| --- | --- | --- | --- | --- | --- |
| MHCI; CD45 | M1/42;  30-F11 | Anti-mouse Hashtag #1 TotalSeq-A Oligo | HashT1 (Q7005) | GTGAATAT | 1 μg/test |
| MHCI; CD45 | M1/42;  30-F11 | Anti-mouse Hashtag #2 TotalSeq-A Oligo | HashT2 (Q7006) | ACAGGCGC | 1 μg/test |
| MHCI; CD45 | M1/42;  30-F11 | Anti-mouse Hashtag #3 TotalSeq-A Oligo | HashT3 (Q7007) | CATAGAGT | 1 μg/test |
| MHCI; CD45 | M1/42;  30-F11 | Anti-mouse Hashtag #4 TotalSeq-A Oligo | HashT4 (Q7008) | TGCGAGAC | 1 μg/test |
| MHCI; CD45 | M1/42;  30-F11 | Anti-mouse Hashtag #5 TotalSeq-A Oligo | HashT5 (Q7015) | TCTCTACT | 1 μg/test |
| MHCI; CD45 | M1/42;  30-F11 | Anti-mouse Hashtag #6 TotalSeq-A Oligo | HashT6 (Q7016) | CTCTCGTC | 1 μg/test |
| MHCII  (I-A/I-E) | M5/114.15.2 | TotalSeq-A0117 Oligonucleotide | RPI1 | CGTGAT | 1 μg/test |
| Sca1 (Ly6A/E) | D7 | TotalSeq-A0130 Oligonucleotide | RPI3 | GCCTAA | 1 μg/test |
| CD24 | M1/69 | TotalSeq-A0212 Oligonucleotide | RPI4 | TGGTCA | 1 μg/test |
| CD86 | GL-1 | TotalSeq-A0200 Oligonucleotide | RPI10 | AAGCTA | 1 μg/test |

***Supplementary Table 3:*** *list of HTO (hashtag barcode) and ADT (CITE-seq) primers used for Droplet sequencing. Unique indexes are shown in red.*

| HTO/ADT ID | Oligonucleotide Sequence |
| --- | --- |
| HashT1 (Q7005) | CAAGCAGAAGACGGCATACGAGATGTGAATATGTGACTGGAGTTCAGACGTGTGC |
| HashT2 (Q7006) | CAAGCAGAAGACGGCATACGAGATACAGGCGCGTGACTGGAGTTCAGACGTGTGC |
| HashT3 (Q7007) | CAAGCAGAAGACGGCATACGAGATCATAGAGTGTGACTGGAGTTCAGACGTGTGC |
| HashT4 (Q7008) | CAAGCAGAAGACGGCATACGAGATTGCGAGACGTGACTGGAGTTCAGACGTGTGC |
| HashT5 (Q7015) | CAAGCAGAAGACGGCATACGAGATTCTCTACTGTGACTGGAGTTCAGACGTGTGC |
| HashT6 (Q7016) | CAAGCAGAAGACGGCATACGAGATCTCTCGTCGTGACTGGAGTTCAGACGTGTGC |
| HashT1 (Q7005) | CAAGCAGAAGACGGCATACGAGATGTGAATATGTGACTGGAGTTCAGACGTGTGC |
| RPI 1 | CAAGCAGAAGACGGCATACGAGATCGTGATGTGACTGGAGTTCCTTGGCACCCGAGAATTCCA |
| RPI 3 | CAAGCAGAAGACGGCATACGAGATGCCTAAGTGACTGGAGTTCCTTGGCACCCGAGAATTCCA |
| RPI 4 | CAAGCAGAAGACGGCATACGAGATTGGTCAGTGACTGGAGTTCCTTGGCACCCGAGAATTCCA |
| RPI 10 | CAAGCAGAAGACGGCATACGAGATAAGCTAGTGACTGGAGTTCCTTGGCACCCGAGAATTCCA |
| RPI 13 | CAAGCAGAAGACGGCATACGAGATTTGACTGTGACTGGAGTTCCTTGGCACCCGAGAATTCCA |
| RPI 11 | CAAGCAGAAGACGGCATACGAGATGTAGCCGTGACTGGAGTTCCTTGGCACCCGAGAATTCCA |

***Supplementary Table 4:*** *Marker gene lists of Ly6d+ and Ly6d- intertypical TEC from scRNA-seq data and Xenium spatial transcriptomics platform.*

| **Ly6d_pos** | **Ly6d_neg** | **Xenium_Ly6d_pos** | **Xenium_Ly6d_neg** |
| --- | --- | --- | --- |
| Fcgbp | Tagln | Ackr1 | Abca1 |
| Krt14 | Apoe | Acvr1 | Ackr3 |
| Krt5 | Igfbp2 | Adcy2 | Adam28 |
| Hspb1 | Gpx3 | Adgrg1 | Adamdec1 |
| Wfdc18 | Lifr | Agrn | Angpt1 |
| Ly6a | Hpgd | Ahr | Angptl4 |
| Igfbp4 | Mgp | Ankrd11 | Anxa2 |
| Krtdap | Anxa2 | Anxa6 | Aox3 |
| C3 | Gas1 | Aqp5 | Aplp2 |
| Gsn | Ptprz1 | Bcl11b | Arid5b |
| Fbxo32 | Dcn | Bmpr1b | Bace1 |
| Gm8113 | Wfikkn2 | Boc | Bcam |
| Cald1 | Oat | C3 | Bmp6 |
| Tagln | Ifit3 | Cacna1g | Bmpr1a |
| Clca3a2 | Col6a1 | Cadm3 | Calcrl |
| Dapl1 | Sparc | Casq2 | Casp12 |
| Tgfbi | Plscr2 | Ccl11 | Cav1 |
| Itga6 | Fzd2 | Cd276 | Ccn2 |
| Emp2 | Limch1 | Cdc42 | Ccnd3 |
| Acta2 | Hif1a | Celsr2 | Cd34 |
| Ifit3 | Osmr | Clca3a2 | Cdh3 |
| Gas1 | Csrp2 | Cldn1 | Cers4 |
| Col6a1 | Myl9 | Cntn2 | Chrd |
| Irf7 | Iigp1c | Col6a1 | Cldn1 |
| Ifi203 | Il6st | Col6a2 | Col6a1 |
| Htra1 | Mme | Col7a1 | Col6a2 |
| Parp14 | Id3 | Col9a3 | Cyp1b1 |
| Cyp2f2 | Wls | Cops9 | Dab2 |
| Fst | Ifitm2 | Cxadr | Dll1 |
| Fxyd3 | Bcam | Cxcl14 | Ecscr |
| Ifi202b | Itga1 | Dag1 | Efemp1 |
| Cavin3 | Rflnb | Dapk1 | Epha1 |
| Urah | Calcrl | Ddit4 | Ets2 |
| Ifi204 | C1s1 | Dgka | F3 |
| Smoc2 | Pdgfa | Dhx58 | Fat1 |
| Ifit1 | Slfn5 | Dkk3 | Fmod |
| Col6a2 | Ifit3b | Dpy30 | Fst |
| Myh9 | Acta2 | Dpysl2 | Fzd2 |
| Mndal | Lama3 | Dusp7 | Gas1 |
| Ifi44 | Ifit1 | Duxbl1 | Ghr |
| Dgkz | Crim1 | Dync1h1 | Glg1 |
| Phf11d | Pdpn | Emc2 | Gna14 |
| Map1lc3a | Cst3 | Emc6 | Gpr88 |
| Aopep | Vwa1 | Enpp1 | Hif1a |
| Ncam1 | Fst | Epb41l2 | Hoxa7 |
| Tacstd2 | Rcn3 | Ephb3 | Hpcal4 |
| Mpzl2 | Serpinb6b | Epn3 | Hpse2 |
| Runx1 | Myl6 | Esyt2 | Ifit1 |
| Shisa4 | Rasl11a | Eya4 | Ifit3 |
| Skint7 | Txndc17 | Ezr | Ifnar2 |
| Scx | Col6a2 | F3 | Igfbp2 |
| H2-T22 | Twsg1 | Fbxl7 | Igfbp3 |
| Rnf213 | Cdh3 | Fbxo32 | Il13ra1 |
| Robo1 | Pgm5 | Fst | Il13ra2 |
| Zbp1 | Slco2a1 | Furin | Il1r1 |
| Slfn2 | Sult5a1 | Fzd7 | Il33 |
| Slfn5 | Kcnma1 | Galnt14 | Il6st |
| Trim30a | Lox | Gas1 | Irs1 |
| Ifit3b | Kazald1 | Gbp3 | Itga1 |
| Oasl2 | Aplp2 | Gpr107 | Itga9 |
| Ccl21d | Angpt1 | Gpx2 | Kcnma1 |
| NA | Apoc1 | Grpel1 | Lama3 |
| Stat2 | Golim4 | Herc6 | Ldlrad4 |
| Adamts9 | Sp100 | Hmcn1 | Lgr4 |
| Mir205hg | Defb1 | Hspa1b | Lifr |
| Eya4 | Insig2 | Ifi204 | Lox |
| Cxcl14 | Cpne8 | Ifi208 | Lrp1 |
| Plec | Mgat3 | Ifi44 | Ltbp1 |
| Tnfsf10 | Kcnj15 | Ifit1 | Mest |
| Tpm1 | C1ra | Ifit3 | Mme |
| Lamb3 | Igfbp3 | Ifnlr1 | Moxd1 |
| Itgb4 | Serping1 | Il18r1 | Mrtfa |
| Gbp3 | Pros1 | Inpp4b | Musk |
| Picalm | Il1r1 | Irf7 | Npy1r |
| Il18r1 | Rragd | Itga6 | Nt5e |
| Pmepa1 | Ntf5 | Itgb4 | Ntf5 |
| Mill1 | Ecscr | Itgb5 | Osmr |
| Zbtb20 | F3 | Jag1 | Pak3 |
| Ltbp2 | Fmod | Kcnh1 | Pakap |
| Robo2 | Opcml | Kdm5a | Pdgfa |
| Ccl11 | Sec62 | Kif5c | Pdpn |
| Sdc3 | Rnf150 | Klc3 | Peg3 |
| Rigi | Tnik | Klf10 | Pik3r1 |
| Ddx60 | Wwc2 | Kras | Prkci |
| C1s1 | Rbfox1 | Lacc1 | Ptgdr |
| Mycbp2 | Angptl4 | Lama5 | Ptger4 |
| Slc27a3 | Arid5b | Lamb3 | Ptprz1 |
| Krt15 | Tmem64 | Lamtor5 | Rab27a |
| Dag1 | Rchy1 | Large2 | Rab38 |
| Boc | Timp2 | Lgr6 | Rragd |
| Zfp703 | Ston2 | Limk2 | Sdc2 |
| Klf13 | Pik3r1 | Lin7c | Serpina3n |
| Ptgfrn | Abca1 | Lrig1 | Serpini1 |
| Irf6 | Adam28 | Lrp1 | Smpd3 |
| Cxadr | Il33 | Malt1 | Spaca6 |
| Foxn3 | Scn7a | Map1lc3a | Spon1 |
| Kctd4 | Lgr4 | Mcam | Sulf2 |
| Pbx1 | Qrfp | Mef2a | Syne4 |
| Sult5a1 | Cd302 | Met | Tagln |
| Calml3 | Xdh | Mpc1 | Thbs2 |
| Herc6 | Pak3 | Mycbp2 | Tmem98 |
| Ddit4 | Rabac1 | Myh9 | Tnfsf12 |
| Xylt1 | Anxa8 | Ncam1 | Tns2 |
| Vcl | Trbc2 | Ncbp2 | Vsir |
| Sdc1 | G930009F23Rik | Ncor2 | Wls |
| Ccdc141 | Slc3a2 | Ndufa12 | Wnt5a |
| Nfe2l1 | Ctps1 | Nfe2l1 | Wnt5b |
| Sbno2 | Serpini1 | Nhlh2 | Wnt6 |
| Gbp7 | Pdzrn4 | Nod2 |  |
| Parp8 | Ctdsp2 | Oasl2 |  |
| Ankrd11 | Mfap4 | Ogt |  |
| Pou3f1 | Flrt3 | Oxtr |  |
| Anxa6 | Map3k7cl | P2ry1 |  |
| Oas3 | Casp12 | Pawr |  |
| Dtx3l | Bmpr1a | Pdgfb |  |
| Slc6a6 | Ypel3 | Picalm |  |
| Lacc1 | Atp1b3 | Piezo2 |  |
| Cpxm2 | Il13ra1 | Plcb4 |  |
| mt-Co3 | Fat1 | Plcg1 |  |
| Skint8 | Nrbp2 | Plec |  |
| Pdlim3 | Antxr1 | Plekha1 |  |
| Ifi211 | Sorcs2 | Plppr4 |  |
| Smad7 | Ets2 | Pml |  |
| Gbp5 | Sdc2 | Pou3f1 |  |
| Apol9a | Ifnar2 | Ppp1r2 |  |
| Nrbp2 | Mansc4 | Prrx1 |  |
| mt-Co1 | Rgma | Prss41 |  |
| Zfand5 | Vav3 | Rab7b |  |
| BC034090 | Ptger4 | Rac1 |  |
| Rgs12 | Peg3 | Rala |  |
| Crp | Nudt16 | Rarg |  |
| Plekha1 | Serpina3n | Rhoa |  |
| Lgals7 | Ccnd3 | Rhoj |  |
| Mx1 | Emb | Robo1 |  |
| Epb41l2 | Lurap1l | Robo2 |  |
| Stx16 | Tm9sf3 | Runx1 |  |
| Pdzd2 | Pde7a | Runx2 |  |
| Phldb2 | Tmem123 | Samhd1 |  |
| Sh3pxd2b | Mmd | Scn3b |  |
| Sectm1a | Ltbp1 | Scx |  |
| Parp9 | Lrp1 | Sdc1 |  |
| Agrn | Smpd3 | Sdhb |  |
| Serpinb5 | Avpr1a | Sdk2 |  |
| Hspa1b | Fat3 | Selenos |  |
| F3 | Pls3 | Sema3a |  |
| Dnase2a | Gxylt2 | Sema3f |  |
| Skil | Nckap5 | Sema6a |  |
| Slfn8 | Il13ra2 | Serpinb5 |  |
| Lrp1 | Thbs2 | Serpine1 |  |
| Fbxl7 | Esyt1 | Sfxn3 |  |
| Slc4a7 | Bace1 | Sh3rf2 |  |
| Samhd1 | AW011738 | Sin3b |  |
| Ogt | Tmem98 | Slc16a2 |  |
| Rnf150 | Tmem47 | Slc4a7 |  |
| Svil | Cyp1b1 | Slc9a1 |  |
| Dync1h1 | Ghr | Smad7 |  |
| Slc4a11 | Slc1a4 | Smdt1 |  |
| Clip1 | Sulf2 | Smoc2 |  |
| Galnt14 | Glg1 | Snx5 |  |
| Pdxdc1 | Ecrg4 | Sod2 |  |
| Synpo2 | Thsd4 | Srgap1 |  |
| Plek2 | Chl1 | Ssbp2 |  |
| Mef2a | Prkci | Stat2 |  |
| Scara3 | Clca3a1 | Sumo1 |  |
| Jag1 | Bcap29 | Svil |  |
| Fat2 | Ctsf | Synpo |  |
| Antxr1 | Cldn1 | Synpo2 |  |
| Sh3d19 | Ctnnd2 | Tacstd2 |  |
| Nlrc5 | Mn1 | Tagln |  |
| Tanc1 | Tnfsf12 | Tfap2a |  |
| Frmd4a | Cers4 | Tmem259 |  |
| Nop53 | Luzp2 | Tmem43 |  |
| Gprin3 | Heph | Tnfsf10 |  |
| Mitd1 | Hpse2 | Tox |  |
| Pnisr | Arhgef25 | Trem2 |  |
| Phip | Gna14 | Trim63 |  |
| Cadm4 | Eola1 | Trpm6 |  |
| Sdk2 | Rab27a | Trpv4 |  |
| Lgals8 | Cnksr3 | Ube2m |  |
| Elk3 | Unc79 | Vcl |  |
| Tmem132a | Col5a2 | Vdac2 |  |
| Col17a1 | Art4 | Vit |  |
| Pawr | Epha1 | Vsir |  |
| Lrp1b | Wnt5a | Wfdc18 |  |
| Irf2bpl | Pcolce | Yif1a |  |
| Ube2ql1 | Tspan4 | Zbp1 |  |
| Nedd9 | Maea | Zeb1 |  |
| Lmod1 | Dlc1 | Zfpm2 |  |
| Tns4 | Cd34 | Zup1 |  |
| Unc45a | Wnt6 |  |  |
| Actn4 | B4galt4 |  |  |
| Gpr87 | Rab30 |  |  |
| Pml | Cpq |  |  |
| Mbnl2 | Pex5l |  |  |
| Tnfrsf12a | Fam171b |  |  |
| Malt1 | Mfap2 |  |  |
| Sema3f | Ice2 |  |  |
| Bcl11b | Mbip |  |  |
| mt-Nd6 | Naaladl2 |  |  |
| Dhx58 | Syne4 |  |  |
| Kif5c | Ehd2 |  |  |
| Dapk1 | 2810433D01Rik | |  |
| Trim12a | Trim68 |  |  |
| Tor1aip1 | Insyn1 |  |  |
| Clca3a1 | Arhgap42 |  |  |
| Marf1 | Rab38 |  |  |
| Inpp4b | Nr1d2 |  |  |
| Rin2 | Adamts19 |  |  |
| Ppp1r12b | Dll1 |  |  |
| Numa1 | Rcan3 |  |  |
| Nipal2 | St8sia2 |  |  |
| Apol9b | Efemp1 |  |  |
| Furin | Spaca6 |  |  |
| Pde7a | Maml2 |  |  |
| Ssbp2 | Gm29797 |  |  |
| Sema6a | Arhgap22 |  |  |
| Trim8 | Tmem141 |  |  |
| Gpx2 | C1s2 |  |  |
| Camta1 | Plekhf1 |  |  |
| Dusp23 | Mest |  |  |
| Fam53b | Wasf3 |  |  |
| Fry | Tns2 |  |  |
| Limk2 | Eva1b |  |  |
| Lrig1 | Gpr88 |  |  |
| Scn3b | Plekhh2 |  |  |
| Proser2 | Trgc1 |  |  |
| Celsr2 | Otulinl |  |  |
| Vsnl1 | Nt5e |  |  |
| Pkp3 | Irs1 |  |  |
| Ncor2 | Col3a1 |  |  |
| Trim25 | Dab2 |  |  |
| Tgif1 | Moxd1 |  |  |
| Srgap1 | Ago4 |  |  |
| Srrm2 | Ldlrad4 |  |  |
| Slc27a1 | Lhfpl6 |  |  |
| Mgat5b | Entpd2 |  |  |
| Adgrg1 | Lnx1 |  |  |
| Tox | Tmem255a |  |  |
| Trem2 | Lmntd1 |  |  |
| Sema3a | Colec12 |  |  |
| Lama5 | Flrt2 |  |  |
| Parp12 | Prkx |  |  |
| Fnbp4 | Cables1 |  |  |
| Slitrk4 | Tdrp |  |  |
| Ifi213 | Bmp6 |  |  |
| Sobp | Hpcal4 |  |  |
| Klhl24 | C1rl |  |  |
| Plppr4 | Chrd |  |  |
| Klc3 | Ackr3 |  |  |
| Marchf8 | Nptx1 |  |  |
| Alpk2 | Itga9 |  |  |
| Ppp1r9b | Cfh |  |  |
| Lrig3 | Ogfod3 |  |  |
| Cldn1 | Spon1 |  |  |
| Mid1 | Vsir |  |  |
| Trp53i11 | Samd4 |  |  |
| Zfp282 | Plcxd1 |  |  |
| Fzd7 | Gm11627 |  |  |
| Arhgef5 | Slc27a6 |  |  |
| Synpo | Ms4a4d |  |  |
| Ahr | C1qtnf3 |  |  |
| Gm10600 | Mrtfa |  |  |
| Pxdc1 | Ccn4 |  |  |
| Trim63 | Fcor |  |  |
| Oplah | Adamdec1 |  |  |
| Ass1 | Gm17501 |  |  |
| Esyt2 | Cav1 |  |  |
| Slc9a1 | Podn |  |  |
| Col9a3 | Enox1 |  |  |
| Zeb1 | A730020M07Rik | |  |
| Xrn1 | Ptgdr |  |  |
| Klf10 | Necab1 |  |  |
| Ehd2 | Ccn2 |  |  |
| Nexn | Tmem100 |  |  |
| Ccnl2 | 2010009K17Rik | |  |
| Rarg | Gm9898 |  |  |
| Vps13b | Pdgfd |  |  |
| Dusp7 | Aox3 |  |  |
| Vit | Npy1r |  |  |
| Fam193a | Ptprm |  |  |
| Oas2 | Fam110c |  |  |
| Cep170b | Pid1 |  |  |
| Cyth3 | Musk |  |  |
| Gdpd5 | A630023A22Rik | |  |
| Large2 | B3gnt9 |  |  |
| Trabd2b | Gm16075 |  |  |
| Zcchc2 | Cdh7 |  |  |
| Akr1b8 | NA |  |  |
| Duoxa1 | NA |  |  |
| Pdgfb | BC107364 |  |  |
| Tmem259 | C2cd4d |  |  |
| Sez6l | Wnt5b |  |  |
| Heca | Fbxo27 |  |  |
| Zbtb1 | Hoxa7 |  |  |
| Zup1 | Gm48529 |  |  |
| Eva1c | NA |  |  |
| Mob3b | Gm7233 |  |  |
| Hip1 | Pde1a |  |  |
| Arhgef19 | A730056A06Rik | |  |
| Clca2 | Gm13052 |  |  |
| Serinc5 | Gm16701 |  |  |
| Gpr107 | Nrsn1 |  |  |
| Adamts17 | Gm15337 |  |  |
| Erf | NA |  |  |
| Fibin | Rhox4a |  |  |
| Cplx2 | Gm8104 |  |  |
| Anpep | Gm27040 |  |  |
| Helb | Gm39075 |  |  |
| Ltbp4 | Gm26831 |  |  |
| Slc16a2 | C1qtnf7 |  |  |
| Serpine1 | 5730471H19Rik | |  |
| Kcnh1 | Gm43425 |  |  |
| Arhgap23 | Gm42706 |  |  |
| Gm15867 | 4932411K12Rik | |  |
| Tmem43 | Gm14319 |  |  |
| Tmtc1 | Semp2l1 |  |  |
| Kdm5a | 1700065J11Rik | |  |
| Mapk6 | NA |  |  |
| Dkk3 | Or52e8 |  |  |
| Plekho1 | Gm47897 |  |  |
| Sfxn3 | Kctd16 |  |  |
| Aqp5 | Gm45833 |  |  |
| Kansl1 | Dhx58os |  |  |
| Rgs10 | Ddi1 |  |  |
| Vsir | D530033B14Rik | |  |
| Epn3 | Gm36642 |  |  |
| Sertad4 | Gm12023 |  |  |
| Hs6st1 | Pcdha9 |  |  |
| Tfap2a | Hoxa6 |  |  |
| Helz2 | Pakap |  |  |
| Fyb2 | Or5h17 |  |  |
| Ppp1r13l | Gm16347 |  |  |
| Rin1 | Gm26600 |  |  |
| Col7a1 | Pard3bos3 |  |  |
| Gm14003 | Gm34983 |  |  |
| Ephb3 | Gm31463 |  |  |
| Bmal1 | Gm10830 |  |  |
| Mcam | NA |  |  |
| Zfp532 | Gm37406 |  |  |
| Paox | Gm35453 |  |  |
| Plcg1 | Gm16043 |  |  |
| Met | Spata46 |  |  |
| Tnk2 | Vmn1r20 |  |  |
| Zcchc24 | 4930412F09Rik | |  |
| Cadm3 | Gm12256 |  |  |
| Flrt2 | Gm30489 |  |  |
| Dgka | Or5b120 |  |  |
| Mpv17l | Fthl17f |  |  |
| Zmat4 | Gm46515 |  |  |
| Pcyox1l | Gm11685 |  |  |
| Ifnlr1 | 2210418O10Rik | |  |
| Adcy2 | Gssos2 |  |  |
| Ifi206 | Gm28539 |  |  |
| Greb1l | Gm47030 |  |  |
| B3galt2 | Vmn2r30 |  |  |
| Arhgap24 | Gm12977 |  |  |
| Itgb5 | NA |  |  |
| Fam78b | Or6c69c |  |  |
| Metap1d | Tex13d |  |  |
| Srd5a1 | Gm37021 |  |  |
| Cd276 | Gm14330 |  |  |
| Sh3rf2 | NA |  |  |
| Acvr1 | Gm8094 |  |  |
| Ccdc3 | Pcdha2 |  |  |
| Nod2 | Gm15961 |  |  |
| Gm16233 | Gm12970 |  |  |
| Prrx1 | Gm28075 |  |  |
| Caskin2 | Gm31126 |  |  |
| P2ry1 | Gm42704 |  |  |
| E130114P18Rik | 1700044K03Rik | |  |
| Piezo2 | Jund |  |  |
| Prag1 |  |  |  |
| Col2a1 |  |  |  |
| A730020M07Rik | |  |  |
| Ccdc125 |  |  |  |
| Pitpnm2 |  |  |  |
| St3gal4 |  |  |  |
| Numbl |  |  |  |
| Aff3 |  |  |  |
| Trim14 |  |  |  |
| Tbc1d2 |  |  |  |
| Runx2 |  |  |  |
| Gm49087 |  |  |  |
| Bmpr1b |  |  |  |
| Atp6v1c2 |  |  |  |
| Hmcn1 |  |  |  |
| Prickle1 |  |  |  |
| Trpv4 |  |  |  |
| Ifi208 |  |  |  |
| Gjb4 |  |  |  |
| Rhoj |  |  |  |
| Lrrc32 |  |  |  |
| Rab7b |  |  |  |
| Bmerb1 |  |  |  |
| Tram2 |  |  |  |
| Trpm6 |  |  |  |
| Ackr1 |  |  |  |
| Cyp2d22 |  |  |  |
| Shroom4 |  |  |  |
| 3110099E03Rik | |  |  |
| Gm40849 |  |  |  |
| Porcn |  |  |  |
| Zfpm2 |  |  |  |
| Cntn2 |  |  |  |
| Arsj |  |  |  |
| Prss41 |  |  |  |
| Chd5 |  |  |  |
| A930015D03Rik | |  |  |
| Prkcb |  |  |  |
| Ifitm10 |  |  |  |
| Pcdhb21 |  |  |  |
| NA |  |  |  |
| Fhl3 |  |  |  |
| Hectd2 |  |  |  |
| Enpp1 |  |  |  |
| Gm43042 |  |  |  |
| Pcdhb9 |  |  |  |
| 4831440D22Rik | |  |  |
| Duox1 |  |  |  |
| Ifitm7 |  |  |  |
| Lgr6 |  |  |  |
| Cacna1g |  |  |  |
| Gm5084 |  |  |  |
| Vmn2r90 |  |  |  |
| Pcdhga9 |  |  |  |
| Oxtr |  |  |  |
| Pitpnm3 |  |  |  |
| A730049H05Rik | |  |  |
| Runx2os1 |  |  |  |
| Gm37035 |  |  |  |
| NA |  |  |  |
| NA |  |  |  |
| Gm12681 |  |  |  |
| Gm13293 |  |  |  |
| Casq2 |  |  |  |
| Gm10687 |  |  |  |
| Peak1os |  |  |  |
| NA |  |  |  |
| BC043934 |  |  |  |
| Nhlh2 |  |  |  |
| Gm3287 |  |  |  |
| Gm9821 |  |  |  |
| Kctd16 |  |  |  |
| Gm15775 |  |  |  |
| Gm16892 |  |  |  |
| NA |  |  |  |
| NA |  |  |  |
| Cpne1 |  |  |  |
| Gm9936 |  |  |  |
| NA |  |  |  |
| Ccdc121rt2 |  |  |  |
| Adad1 |  |  |  |
| Gm47900 |  |  |  |
| Gm28723 |  |  |  |
| Gm47040 |  |  |  |
| Sspnos |  |  |  |
| Gm27004 |  |  |  |
| Duxbl1 |  |  |  |
| Gm15908 |  |  |  |
| Cstdc7 |  |  |  |
| Gm42421 |  |  |  |
| NA |  |  |  |
| Gm2164 |  |  |  |
| 1700025N21Rik | |  |  |
| Pmepa1os |  |  |  |
| Or2y13 |  |  |  |
| NA |  |  |  |
| Or52e5 |  |  |  |
| Gm9955 |  |  |  |
| 4930521O11Rik | |  |  |
| Gm45188 |  |  |  |
| Gm20379 |  |  |  |
| Gm13276 |  |  |  |
| 4930542C12Rik | |  |  |
| NA |  |  |  |
| Gm16015 |  |  |  |
| 4930596D02Rik | |  |  |
| Gm29246 |  |  |  |
| Gm47471 |  |  |  |
| Gm39312 |  |  |  |
| Nup58 |  |  |  |
| 4921522P10Rik | |  |  |
| Gm45237 |  |  |  |
| Gm21969 |  |  |  |
| Msantd5f6 |  |  |  |
| Gm8271 |  |  |  |
| Gm43397 |  |  |  |
| Gm13063 |  |  |  |
| Gm47177 |  |  |  |
| NA |  |  |  |
| NA |  |  |  |
| 1700071G01Rik | |  |  |
| Gm35766 |  |  |  |
| Or5b123 |  |  |  |
| Gm28415 |  |  |  |
| Acss2os |  |  |  |
| Gm43894 |  |  |  |
| Gm43904 |  |  |  |
| Or2ak7 |  |  |  |
| Or5p63 |  |  |  |
| Or5ak24 |  |  |  |
| 1700007H22Rik | |  |  |
| Gm40293 |  |  |  |
| Mxra8os |  |  |  |
| C230029F24Rik | |  |  |
| 4930429C20Rik | |  |  |
| Gm48548 |  |  |  |
| Gm7276 |  |  |  |
| Gm45723 |  |  |  |
| Or2r11 |  |  |  |
| Gm28340 |  |  |  |
| Or12e8 |  |  |  |
| Gm2163 |  |  |  |
| Gm47501 |  |  |  |
| NA |  |  |  |
| Gm41118 |  |  |  |
| Vmn1r170 |  |  |  |
| 4933404K08Rik | |  |  |
| Gm15239 |  |  |  |
| Gm45785 |  |  |  |
| Mbd3l1 |  |  |  |
| Gm34333 |  |  |  |
| Ighv1-37 |  |  |  |
| Pfn3 |  |  |  |
| Gm48612 |  |  |  |
| Gm17078 |  |  |  |
| Gm15898 |  |  |  |
| 4930439A04Rik | |  |  |
| Gm28798 |  |  |  |
| Or2f1b |  |  |  |
| NA |  |  |  |
| Pgm3 |  |  |  |
| Traf4 |  |  |  |
| Yif1a |  |  |  |
| Ergic2 |  |  |  |
| Ier3ip1 |  |  |  |
| Lin7c |  |  |  |
| Mrpl36 |  |  |  |
| Hmox2 |  |  |  |
| Tmed4 |  |  |  |
| Selenot |  |  |  |
| Tmco1 |  |  |  |
| Med28 |  |  |  |
| Dph3 |  |  |  |
| Vma21 |  |  |  |
| Ndufb2 |  |  |  |
| Ncbp2 |  |  |  |
| Sin3b |  |  |  |
| Sdf2 |  |  |  |
| Lamtor5 |  |  |  |
| Pfdn1 |  |  |  |
| Cript |  |  |  |
| Coq7 |  |  |  |
| Derl1 |  |  |  |
| Adprh |  |  |  |
| Serp1 |  |  |  |
| Commd6 |  |  |  |
| Vbp1 |  |  |  |
| Ndufb7 |  |  |  |
| Sar1b |  |  |  |
| Tbca |  |  |  |
| Naa38 |  |  |  |
| Timm13 |  |  |  |
| Iscu |  |  |  |
| Trappc3 |  |  |  |
| Ndufa13 |  |  |  |
| Ppp1r11 |  |  |  |
| Chchd1 |  |  |  |
| Snx5 |  |  |  |
| Aurkaip1 |  |  |  |
| Emc2 |  |  |  |
| Coa3 |  |  |  |
| Rac1 |  |  |  |
| Sdhb |  |  |  |
| Hsbp1 |  |  |  |
| Mrpl20 |  |  |  |
| Cuta |  |  |  |
| Mrps21 |  |  |  |
| Pcbd2 |  |  |  |
| Emc7 |  |  |  |
| Glrx5 |  |  |  |
| Ndufs5 |  |  |  |
| Dpysl2 |  |  |  |
| Bzw1 |  |  |  |
| Srp9 |  |  |  |
| Dad1 |  |  |  |
| Ndufa12 |  |  |  |
| Ndufb8 |  |  |  |
| Mrps14 |  |  |  |
| Ubl5 |  |  |  |
| Yipf4 |  |  |  |
| Ndufs7 |  |  |  |
| Rnf11 |  |  |  |
| Ppdpf |  |  |  |
| Mrpl11 |  |  |  |
| Mrps33 |  |  |  |
| Mdh2 |  |  |  |
| Selenos |  |  |  |
| Ndufb11 |  |  |  |
| Rala |  |  |  |
| Sdhd |  |  |  |
| Ndfip2 |  |  |  |
| Ndufs2 |  |  |  |
| Rapgef5 |  |  |  |
| Uqcrc2 |  |  |  |
| Grpel1 |  |  |  |
| Emc6 |  |  |  |
| Atp5pf |  |  |  |
| D8Ertd738e |  |  |  |
| Uqcrb |  |  |  |
| Sys1 |  |  |  |
| Lsm6 |  |  |  |
| Kras |  |  |  |
| Rps27l |  |  |  |
| Arpc1a |  |  |  |
| Azin1 |  |  |  |
| Ndufb10 |  |  |  |
| Ndufc1 |  |  |  |
| Akr1a1 |  |  |  |
| Sumo1 |  |  |  |
| Brk1 |  |  |  |
| Vdac3 |  |  |  |
| Ufc1 |  |  |  |
| Tmem14c |  |  |  |
| Ube2m |  |  |  |
| Ndufa8 |  |  |  |
| Ndufb5 |  |  |  |
| Chmp5 |  |  |  |
| Uqcr10 |  |  |  |
| Tomm22 |  |  |  |
| Cops9 |  |  |  |
| Uqcrfs1 |  |  |  |
| Spcs1 |  |  |  |
| Slc25a3 |  |  |  |
| Swi5 |  |  |  |
| Atp5mf |  |  |  |
| Ost4 |  |  |  |
| Zfand6 |  |  |  |
| Mrpl18 |  |  |  |
| Arf1 |  |  |  |
| Atp5f1b |  |  |  |
| BC031181 |  |  |  |
| Bsg |  |  |  |
| Dpy30 |  |  |  |
| Ndufab1 |  |  |  |
| Plcb4 |  |  |  |
| Smdt1 |  |  |  |
| Naxe |  |  |  |
| Ndufa7 |  |  |  |
| Atp5f1a |  |  |  |
| Gabarapl2 |  |  |  |
| Eif1 |  |  |  |
| Rer1 |  |  |  |
| Gde1 |  |  |  |
| Cox7a2 |  |  |  |
| Atp5mc3 |  |  |  |
| Micos10 |  |  |  |
| Chchd2 |  |  |  |
| Vdac2 |  |  |  |
| Cox8a |  |  |  |
| Clta |  |  |  |
| Cdc42 |  |  |  |
| Oaz1 |  |  |  |
| Ndufb9 |  |  |  |
| Tmbim6 |  |  |  |
| Prdx1 |  |  |  |
| Ddt |  |  |  |
| Cox5a |  |  |  |
| Atp5f1c |  |  |  |
| Sod2 |  |  |  |
| Atp5pb |  |  |  |
| Rhoa |  |  |  |
| Mdh1 |  |  |  |
| Ppp1r2 |  |  |  |
| Etfa |  |  |  |
| Calm3 |  |  |  |
| Ndufc2 |  |  |  |
| Tmem176b |  |  |  |
| Nme1 |  |  |  |
| Srp14 |  |  |  |
| Dbi |  |  |  |
| Ezr |  |  |  |
| Suclg1 |  |  |  |
| Tspan3 |  |  |  |
| Mpc1 |  |  |  |
| Ndufa4 |  |  |  |
| Tmem176a |  |  |  |
| Calm1 |  |  |  |
| Prnp |  |  |  |
